# An Exact-Residue Atlas of Opioid Receptor Wiring and Rewiring across Ligand and Transducer Contexts

**DOI:** 10.64898/2026.08.14.744957

**Authors:** Manal A. Nael, Laxman M. Alakonda, Khaled M. Elokely

## Abstract

Opioid-receptor structures span four human receptor subtypes, diverse ligands, signaling partners, and experimental constructs. We curated 86 human opioid-receptor structures representing 84 independent experimental maps, with one unique experimental data set counted once for structure-level inference, and analyzed them using our in-house StrucMind platform. StrucMind constructs exact Ballesteros-Weinstein (BW) contact graphs, meaning residue-contact networks restricted to unambiguous generic BW positions. Relative to active transducer-bound structures, structures classified as inactive showed 2.49% lower mean contact similarity and 34.84% more rewired contacts, where rewiring is the static set of contacts gained or lost between two structures. Among 77 maps with a resolved selected-ligand site, changed contacts were 15.28% direct to the site, 40.13% adjacent at one graph edge, and 44.59% connected-distal at a finite graph distance greater than one. The deposited-water analysis identified 137 receptor-proximal waters. Sixty-one contacted at least two protein residues, including 38 that bridged at least two exact-BW residues; a separate ligand-contact branch contained 10 waters contacting both selected ligand and receptor, only 3 of which belonged to the 38-water set. None of 32 component-association tests survived global correction. For peptide versus small molecule, the smallest nominal *p* value among four outcomes corresponded to 6.18% lower shared-contact distance root-mean-square deviation (*p* = 0.00989 ; *q* = 0.3165, where *q* is the adjusted *p* value). The atlas supports bounded, testable hypotheses, not causal component, hydration, or efficacy mechanisms.

## Introduction

The mu, kappa, delta, and nociceptin/orphanin FQ opioid receptors (MOR, KOR, DOR, and NOP) are class A G protein-coupled receptors (GPCRs), seven-transmembrane signaling proteins that couple extracellular ligand binding to intracellular transducers, meaning proteins that relay receptor activation. Antagonist-bound structures, in which a ligand blocks agonist-driven activation, established inactive architectures of all four receptors.^1–4^ Later structures bound to agonists, ligands that promote receptor signaling, peptides, heterotrimeric G proteins composed of Gα, Gβ, and Gγ subunits, or β-arrestins, receptor-binding regulatory and signaling adaptors, resolved receptor-specific features of activation, ligand recognition, and signaling.^5–13^ Allosteric modulators, ligands that bind outside the primary, or orthosteric, ligand-binding pocket, further defined receptor modulation in specific systems.^14–16^ An experimental construct is the engineered receptor, fusion, partner, stabilizer, and component composition used for structure determination. These structures create a difficult comparison problem because receptor sequence, state, orthosteric ligand modality, meaning peptide versus small molecule, source-reported activity, signaling partner, construct, membrane-associated components, experimental method, and resolution vary concurrently.

Several mechanisms of interest are established in particular systems. Conserved internal waters participate in GPCR stabilization and activation.^17^ DOR structures and functional experiments define an allosteric sodium pocket.^18^ MOR ligand design has demonstrated water-mediated engagement of the sodium site.^19^ Cholesterol and membrane context can alter opioid-receptor signaling in defined systems.^20, 21^ A recent MOR study placed a cholesterol molecule beside the distinct BMS-986187 allosteric pocket and proposed a cooperative ligand/lipid mechanism.^15^ Membrane phosphoinositides, phosphorylated phosphatidylinositol lipids, regulate GPCR/β-arrestin complex assembly and dynamics in a receptor-dependent manner.^22^ Phosphatidylinositol 4,5-bisphosphate is abbreviated PI(4,5)P₂ or PIP₂. Structures of MOR coupled to *G_z_*, a Gα subtype, and MOR/β-arrestin 1 centered on transmembrane helix 1 (TM1)-dependent transducer-specific signaling and did not present a PIP₂-specific experiment.^13^ By contrast, the engineered KOR preparation containing the vasopressin V2 receptor tail (V2R-tail), β-arrestin 1, and short-chain PI(4,5)P₂, Protein Data Bank (PDB) entry 9ZZO, resolved a PIP₂-binding site and found that mutation of PIP₂-contacting residues reduced β-arrestin recruitment.^12^ A deposited-structure cohort cannot re-create those experiments. It can quantify how often their structural contexts are represented, where associated receptor contacts lie, how large observed differences are, and which contrasts have enough overlap to estimate a cohort-level association.

We assembled a human opioid-receptor cohort of 86 structures representing 84 independent maps, curated ligand and component roles, and applied deposited-water, protein-interface, single-structure, and exhaustive pairwise analyses. Ballesteros-Weinstein mapping assigns each transmembrane residue a helix number and a position relative to the most conserved residue on that helix; exact mapping here requires an unambiguous assignment from the G protein-coupled receptor database (GPCRdb). In a contact graph, residues are nodes and spatial contacts are undirected edges. Wiring is a single structure’s edge set, whereas rewiring is the symmetric difference, or contacts present in only one of two structures. The selected-ligand site is the union of exact-BW receptor residues within 4.5 Å of curated orthosteric and allosteric ligand atoms. Direct contacts touch the site, adjacent contacts are one graph edge from it, connected-distal contacts have a finite graph distance greater than one, and disconnected contacts have no finite path. A percentage-scaled difference is an observed difference or fitted coefficient divided by the absolute observed reference-group mean and multiplied by 100. Prevalence and percentage-scaled differences are reported with uncertainty; publication overlap means that both contrast levels occur within one publication; matching capacity is the number of exact-stratum pairs available; leave-one-out stability is the result after deleting one structure or publication and refitting; and *q* is the Benjamini-Hochberg false-discovery-rate-adjusted *p* value. The first argument is deliberately limited: recognized state-associated directions serve as a literature-directed positive control, not an independent replication. The second argument is a dataset-level contribution: a traceable exact-BW atlas broadens quantitative comparison across opioid receptors while preserving uncertainty and preventing system-specific mechanisms from becoming unsupported cohort-general claims.

Figure 1 links the two architectures that govern this study. The receptor architecture places each subtype’s sequence intervals and selected conserved class A motifs in the same BW frame. The methodological architecture traces cohort discovery, map-independent curation, exact residue mapping, contact-graph construction, all-to-all comparison, role-aware site annotation, spatial localization, water recurrence, statistical modeling, and claim classification. Each step points to its detailed SI section and machine-readable specification, and the figure marks the interpretation boundary imposed by static deposited structures. Figure 2 provides the corresponding three-dimensional (3D) structural context by superposing representative MOR, DOR, KOR, and NOP receptor cartoons and highlighting selected sodium-pocket and activation-switch positions.

**Figure 1.**
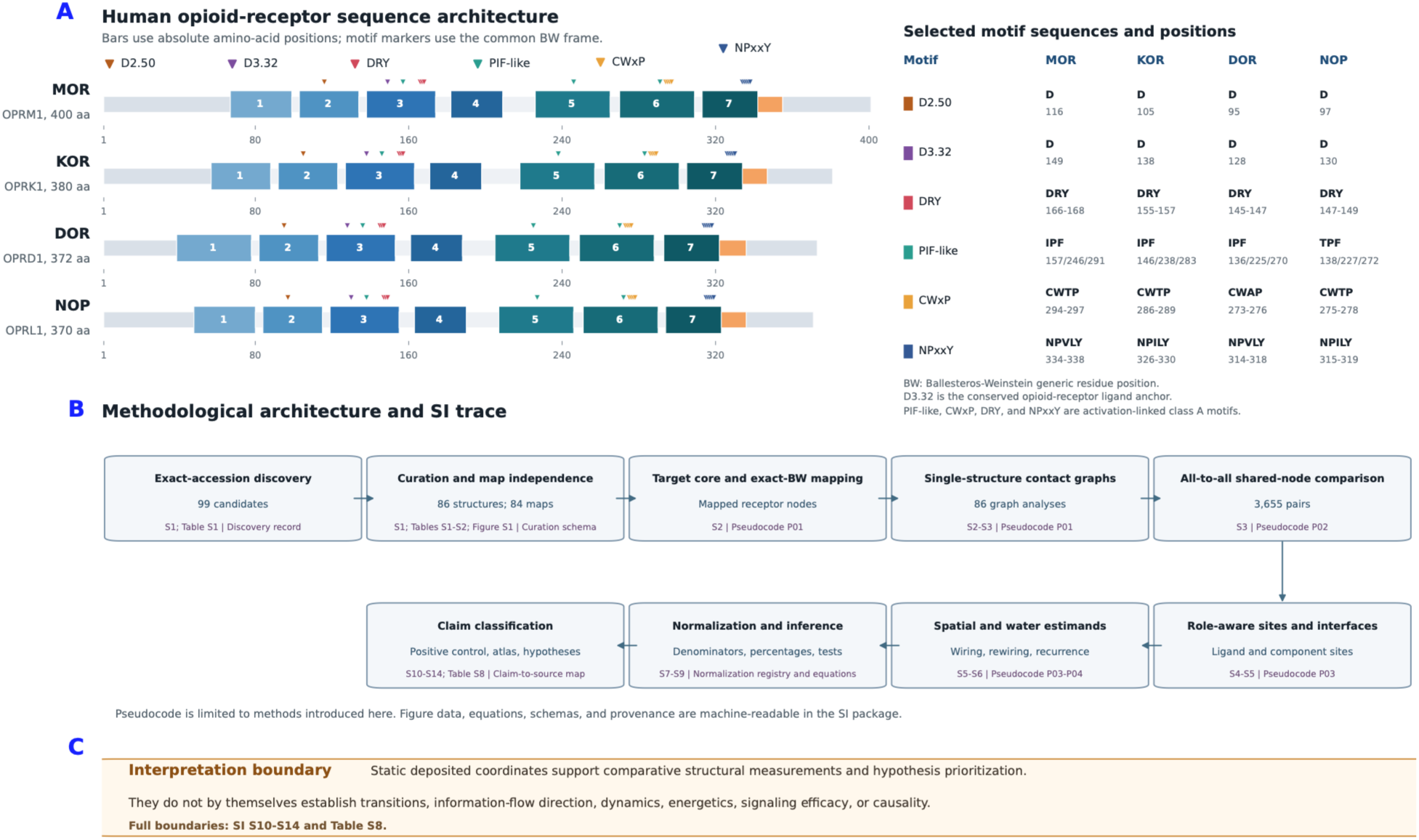
Opioid-receptor sequence architecture and analytical architecture. (A) Linear maps of human MOR (OPRM1, 400 residues), KOR (OPRK1, 380 residues), DOR (OPRD1, 372 residues), and NOP (OPRL1, 370 residues) show the N terminus, transmembrane helices 1 through 7, extracellular and intracellular loops, helix 8, and the C terminus. Markers identify D2.50, the conserved sodium-pocket aspartate; D3.32, the conserved opioid-receptor ligand anchor; the cytoplasmic DRY motif at 3.49 through 3.51; the PIF-like activation connector at 3.40/5.50/6.44; the CWxP motif at 6.47 through 6.50; and the NPxxY motif at 7.49 through 7.53. NOP carries T3.40 rather than I3.40, and DOR carries CWAP rather than CWTP. Absolute positions and segment intervals are receptor-specific, whereas BW labels provide the cross-receptor coordinate system.[25,27] (B) The workflow proceeds from exact-accession discovery and independent-map curation to target-core and BW mapping, our in-house StrucMind single-structure and all-to-all graph analyses, role-aware site annotation, spatial and water estimands, explicit normalization and denominator rules, uncertainty analysis, and bounded claim classification. Each box identifies its detailed SI section, table, figure, or machine-readable specification. Pseudocode P01 through P04 is restricted to methods introduced in this study. (C) Deposited coordinates support static structural comparisons and hypothesis prioritization; the full interpretation boundaries are in SI S10 through S14 and Table S8.

**Figure 2.**
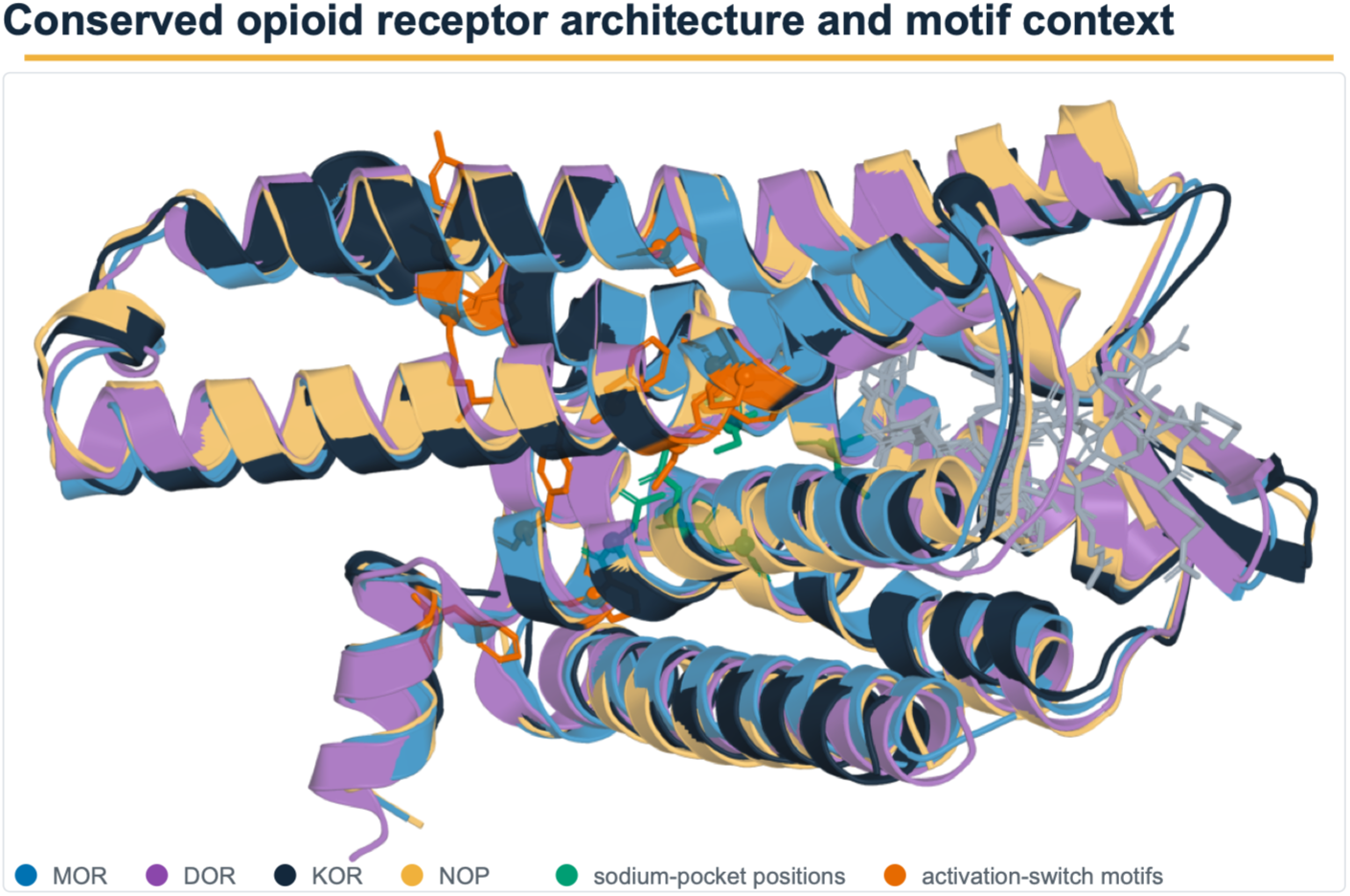
Three-dimensional structural context for conserved opioid-receptor architecture. Aligned receptor cartoons show representative MOR (8F7R, blue), DOR (8F7S, purple), KOR (8F7W, dark navy), and NOP (8F7X, gold) coordinates. Bound peptide ligands are shown as gray sticks. Teal and orange sticks and spheres mark selected sodium-pocket and activation-switch motif positions, respectively, on the KOR reference. The overlay illustrates shared architecture and motif context.

## Results and Discussion

### A curated cohort of 86 human opioid-receptor structures

The cohort registry, a provenance-preserving list of candidate PDB entries, began with 99 candidates. Organism, sequence, construct, and receptor-identity criteria excluded 13 entries, yielding 86 human structures: 35 MOR, 28 KOR, 18 DOR, and 5 NOP. Two pairs share an experimental map, 10TL with 9PU5 and 10TM with 9PUD. Selecting one representative per independent experimental map produced the 84-map primary panel, comprising 33 MOR, 28 KOR, 18 DOR, and 5 NOP maps. The primary panel contains 49 active-partner-bound, 16 inactive-like, 15 unclear, and 4 active-like structures. Active-partner-bound denotes an active annotation with a resolved signaling partner; active-like denotes an active annotation without a resolved partner; inactive-like denotes an inactive annotation or its prespecified title-rule fallback; and unclear denotes no qualifying state rule. Seventy-two maps were obtained by electron microscopy and 12 by X-ray diffraction. The expected-accession target core, defined as selected-chain polymer residues explicitly covered by the Structure Integration with Function, Taxonomy and Sequences (SIFTS) mapping to the intended UniProt receptor accession, retained 24,360 residues and excluded 715. Of those excluded residues, 489 mapped explicitly to non-target UniProt segments and 226 lacked an explicit mapping to the expected target accession. Exact-BW graph nodes form the narrower subset that also has an unambiguous BW label and the atom required by the contact rule.

The curated role annotations define three biologically consequential cases. First, the 84-map binding partition is 72 orthosteric-only, 9 orthosteric-plus-allosteric, 1 allosteric-only, and 2 with no resolved ligand. The adjusted binding-mode contrast therefore compares 9 orthosteric-plus-allosteric maps with 72 orthosteric-only maps. Second, in 9L60, chain P is classified as orthosteric dynorphin A(1-13), whereas MPAM-15 is allosteric.^16^ The primary coordinate-defined site contains 25 exact-BW dynorphin contacts and 10 MPAM-15 contacts, with a 35-position selected-ligand union. The Protein-Ligand Interaction Profiler (PLIP), a rule-based detector of noncovalent contacts in deposited structures, independently resolves 14 dynorphin exact-BW contacts. Third, PIO in 9ZZO is the deposited short-chain diC₈ form of PI(4,5)P₂ and is classified as the analysis category PIP₂-like lipid.^12^ Assembly presence means that the complete modeled Gα/Gβ/Gγ heterotrimer is present. Because those three assembly-presence vectors are identical in this cohort, Gβ-specific analysis was restricted to direct receptor/Gβ contact, defined by a 4.5 Å heavy-atom cutoff, within the 66 Gα-positive maps. Cohort composition and the analytical workflow are summarized in Table 1 and Figure 3, and representative deposited coordinate contexts are shown in Figure 4.

**Figure 3.**
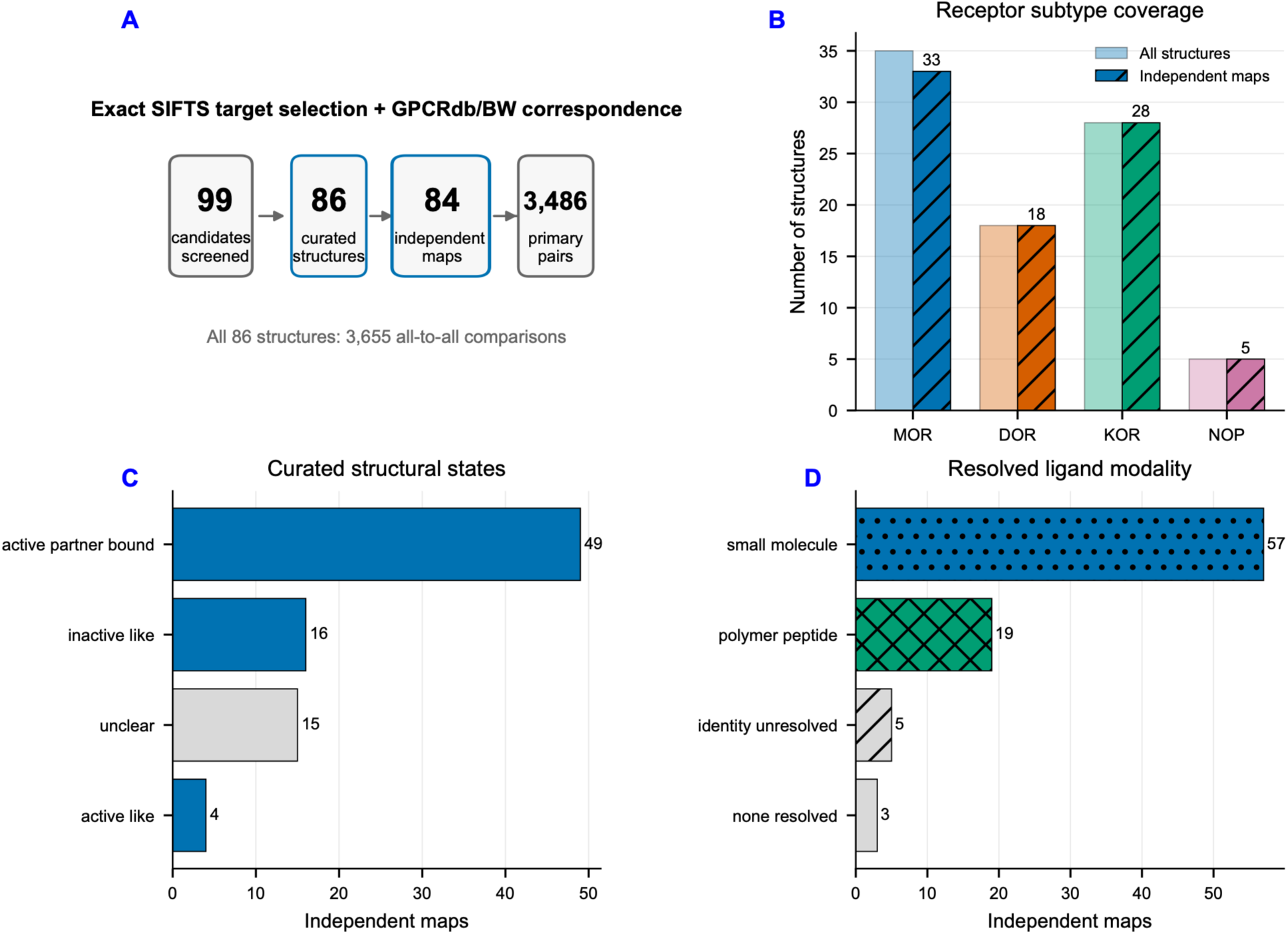
Cohort construction and analysis design. (A) Screening and exact-mapping workflow. Candidate count is the sum of 86 included structures and 13 curation-defined exclusions. Two shared-map model pairs reduce the primary analysis to 84 independent maps and 3,486 primary comparisons; all 86 structures yield 3,655 comparisons. (B-D) Subtype, structural-state, and ligand-modality composition.

**Figure 4.**
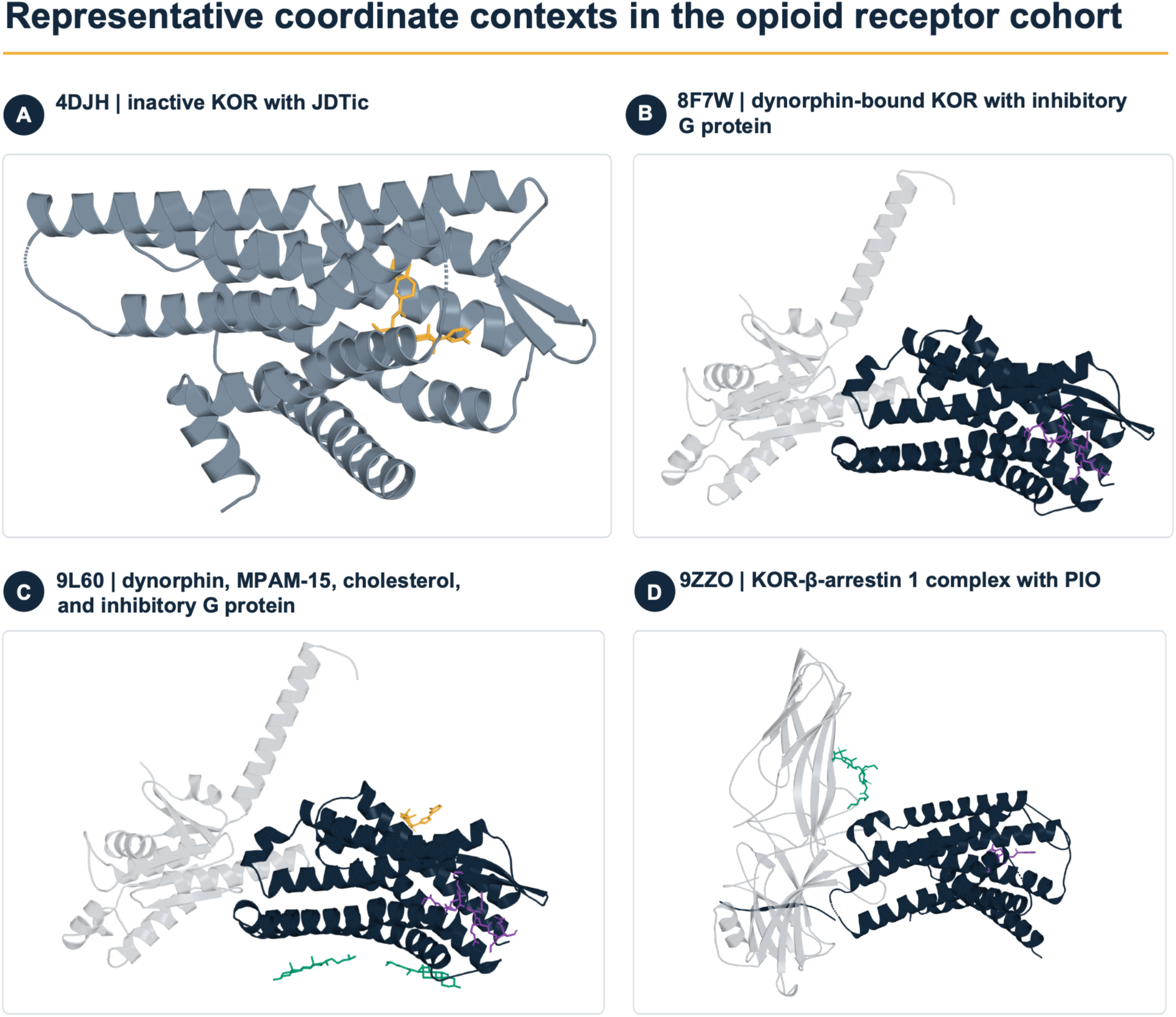
Three-dimensional examples of coordinate contexts represented in the opioid-receptor cohort. (A) Inactive KOR 4DJH with JDTic (gold). (B) Dynorphin-bound KOR 8F7W with the Gα subunit of an inhibitory G protein. (C) KOR 9L60 with dynorphin (purple), MPAM-15 (gold), two cholesterol molecules (teal), and Gα. (D) Engineered KOR/V2R-tail/β-arrestin 1 assembly 9ZZO with PIO, the deposited short-chain diC₈ form of PI(4,5)P₂ (teal), and orthosteric ligand MP1104 (purple). Receptors are dark navy except for slate 4DJH; partner proteins are light gray.

**Table 1.** Composition and analytical scope of the curated opioid-receptor atlas.

| Quantity | Full cohort | Primary independent-map panel | Interpretation |
| --- | --- | --- | --- |
| Structures or maps | 86 structures | 84 maps | Two shared-map relationships retained in provenance |
| MOR/KOR/DOR/NOP | 35/28/18/5 | 33/28/18/5 | NOP is the smallest subtype panel |
| Single-structure analyses | 86 | 84 used for structure-level models | Every curated structure analyzed |
| Unordered all-to-all pairs | 3,655 | 3,486 | Every pair enumerated once |
| Semantic-mode records | 14,620 | Not applicable | Four noninterchangeable modes per full-cohort pair |
| Primary Cα 8 Å shared-node exact-BW contact-comparison records | 3,976,728 | Not applicable | Gained, lost, and conserved edge evidence on nodes resolved in both structures |
| Full-edge-set sensitivity records | 4,026,103 | Not applicable | Includes 49,375 records with at least one endpoint unavailable in one structure |
| Curated binding modes | Not applicable | 72/9/1/2 | Orthosteric / orthosteric plus allosteric / allosteric only / none resolved |
| Selected-ligand union resolved | 79/86 | 77/84 | Unresolved sites remain unknown, not zero |

### Exhaustive exact-BW wiring and rewiring

All 86 single-structure analyses and 3,655 full-cohort pair comparisons completed. Under the primary alpha-carbon (Cα) 8 Å definition, receptor graphs contained 223 to 240 nodes and 965 to 1,128 edges. Across all pairs, the median receptor-core Cα root-mean-square deviation (RMSD), a rigid-body coordinate difference, was 1.306 Å. The median contact Jaccard distance, defined as one minus the fraction of union contacts shared by both structures, was 0.0718. Thus, Jaccard distance is the union-normalized rewiring burden, whereas rewired-contact count retains its raw count scale; both were reported with their distinct denominators. The two measures were strongly concordant (Spearman *ρ* = 0.912, where *ρ* is the rank-correlation coefficient). Cross-subtype comparisons had greater median RMSD than within-subtype comparisons, 1.375 versus 1.034 Å, and greater median contact distance, 0.0769 versus 0.0592. These are descriptive differences because subtype and receptor sequence cannot be separated observationally (Figure 5). Figure 6 shows structural examples at the upper ends of the contact Jaccard-distance and exact-BW Kabsch RMSD distributions; these examples visualize the metric endpoints but do not isolate mechanisms.

**Figure 5.**
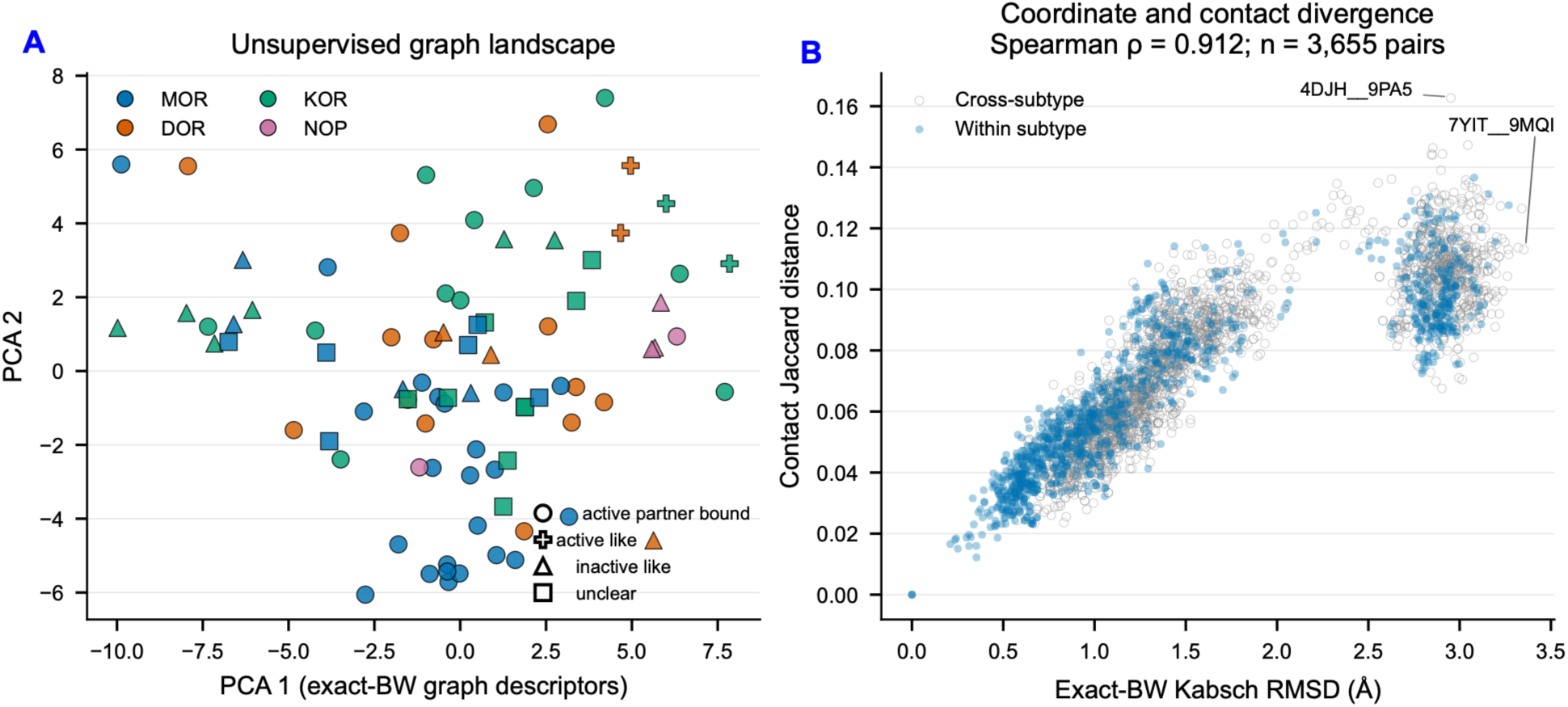
Exact-BW graph landscape across the opioid-receptor cohort. (A) Principal component analysis (PCA) projection for 84 independent maps. Color denotes subtype and shape denotes curated state; PCA is descriptive, not a validated state classifier. (B) Contact Jaccard distance versus coordinate RMSD for all 3,655 pairs. Within-and cross-subtype pairs use filled and open symbols. Spearman ρ = 0.912.

**Figure 6.**
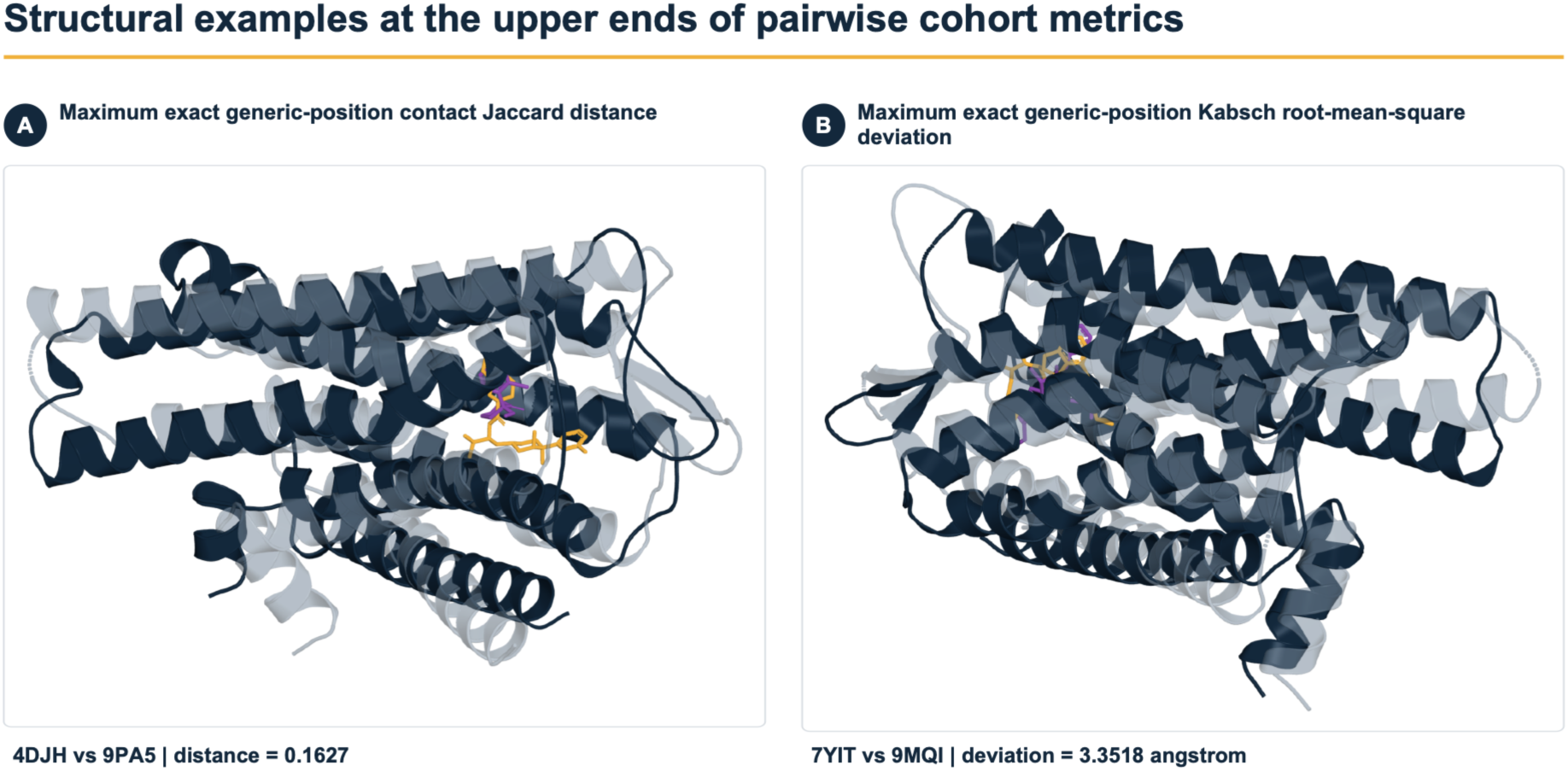
Three-dimensional examples at the upper ends of the pairwise cohort metrics. (A) Overlay of KOR 4DJH (slate) and MOR 9PA5 (dark navy), the pair with the maximum exact-BW contact Jaccard distance, 0.1627. (B) Overlay of KOR 7YIT (slate) and MOR 9MQI (dark navy), the pair with the maximum exact-BW Kabsch root-mean-square deviation, 3.3518 Å. Deposited ligands are shown as gold and purple sticks.

The primary graph was accompanied by Cα 6 Å, Cα 10 Å, beta-carbon (Cβ) 8 Å, minimum heavy-atom 4.5 Å, and minimum side-chain 5 Å definitions. Relative to Cα 8 Å, the median edge-set Jaccard similarity and median current-flow correlation were 0.678 and 0.865 for Cα 6 Å, 0.577 and 0.915 for Cα 10 Å, 0.876 and 0.937 for Cβ 8 Å, 0.858 and 0.896 for minimum heavy-atom 4.5 Å, and 0.313 and 0.495 for minimum side-chain 5 Å. Current-flow betweenness is an electrical-network graph centrality that measures how much effective flow passes through a node across source-target pairs. The sparse side-chain graph is therefore a sensitivity bound, not an independent confirmation. Exact-BW versus whole-core comparisons were most stable for global fingerprints and elastic-network summaries, which are harmonic-motion proxies, and less stable for local flow, graph curvature, perturbation-response proxies, and community assignments, which are algorithmic graph partitions (SI Figures S3 and S4).

### State directions provide a literature-directed workflow positive control

The literature-directed state analysis compared inactive-like with active-partner-bound structures across 16 estimable outcomes and applied Benjamini-Hochberg correction within that defined positive-control family. An endpoint mean is the mean, for one structure, of an outcome across every eligible pair incident on that structure. Six outcomes retained *q* < 0.05 (Table 2; Figure 7). Relative to the observed active-partner-bound mean, inactive-like structures had 2.49% lower endpoint-mean contact Jaccard similarity, 34.84% more rewired contacts, 34.78% more adjacent changed contacts, 25.05% more direct changed contacts, 15.15% greater endpoint-mean shared-contact distance RMS deviation, and 1.67% fewer exact-BW contact pairs. In a full endpoint leave-one-out analysis, meaning that one structure was removed and the analysis was recomputed, every eligible structure was deleted in turn, all pairwise records involving that structure were removed, all remaining endpoint means were recomputed, and the model was refit. All 378 outcome-specific refits completed, and each of the six retained coefficients preserved its sign in 100% of deletions. Percentage estimates in Table 2 include two-sided 95% confidence intervals (CIs). Figure 8 supplies a representative inactive-to-active KOR overlay and mapped G protein α-subunit interface context for structural orientation only.

**Figure 7.**
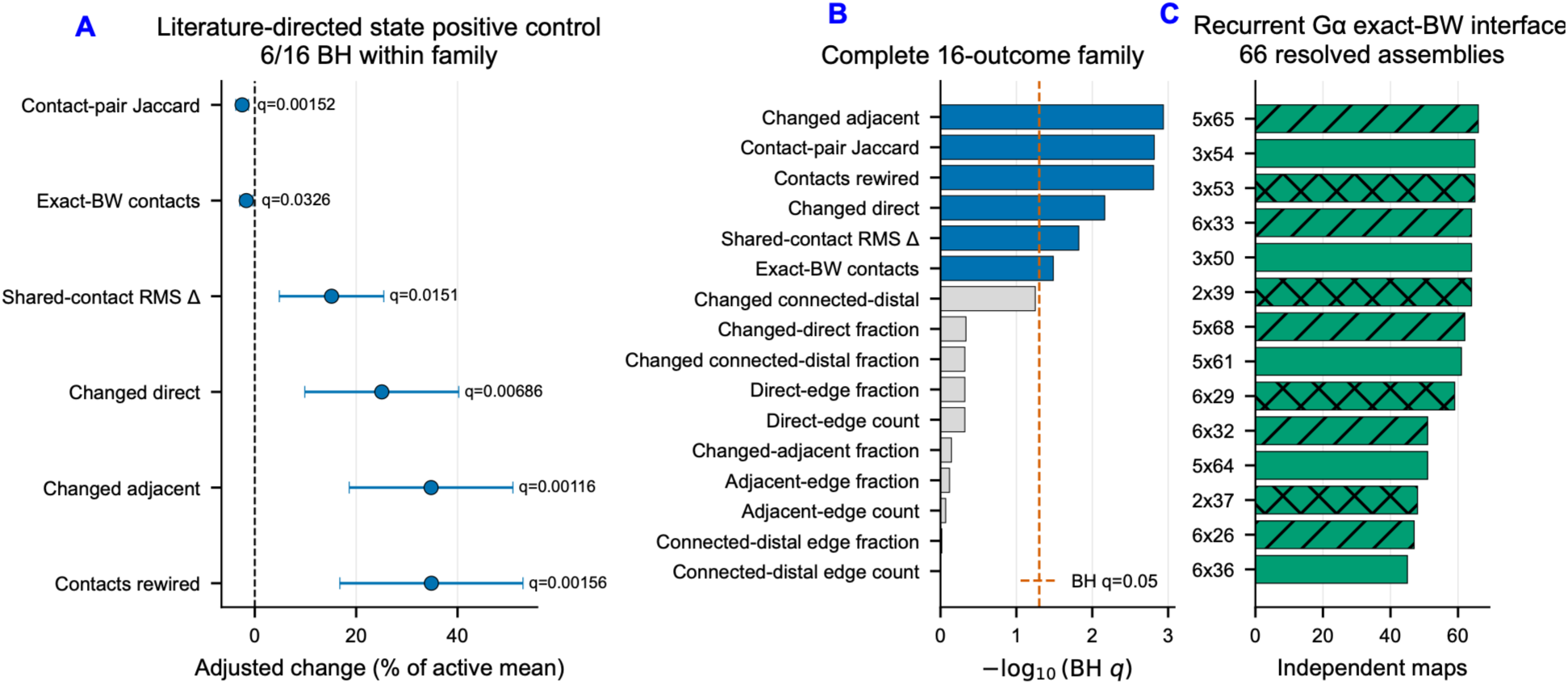
Literature-directed state positive control and the defined Gα interface. (A,B) The literature-directed inactive-like versus active-partner-bound positive-control family contains 16 prespecified outcomes; 6 pass Benjamini-Hochberg (BH) correction within this validation family. Adjusted changes relative to the active mean include Jaccard −2.49%, rewired contacts +34.84%, changed-adjacent +34.78%, changed-direct +25.05%, shared-contact RMS distance +15.15%, and exact-BW contacts −1.67%; each has 100% full endpoint leave-one-structure-out sign consistency. The directions agree with published inactive-versus-active and transducer-bound structures. This is a within-cohort positive control, not independent replication or a general component effect. (C) Exact-BW Gα sites are counted across 66 resolved G-protein maps.

**Figure 8.**
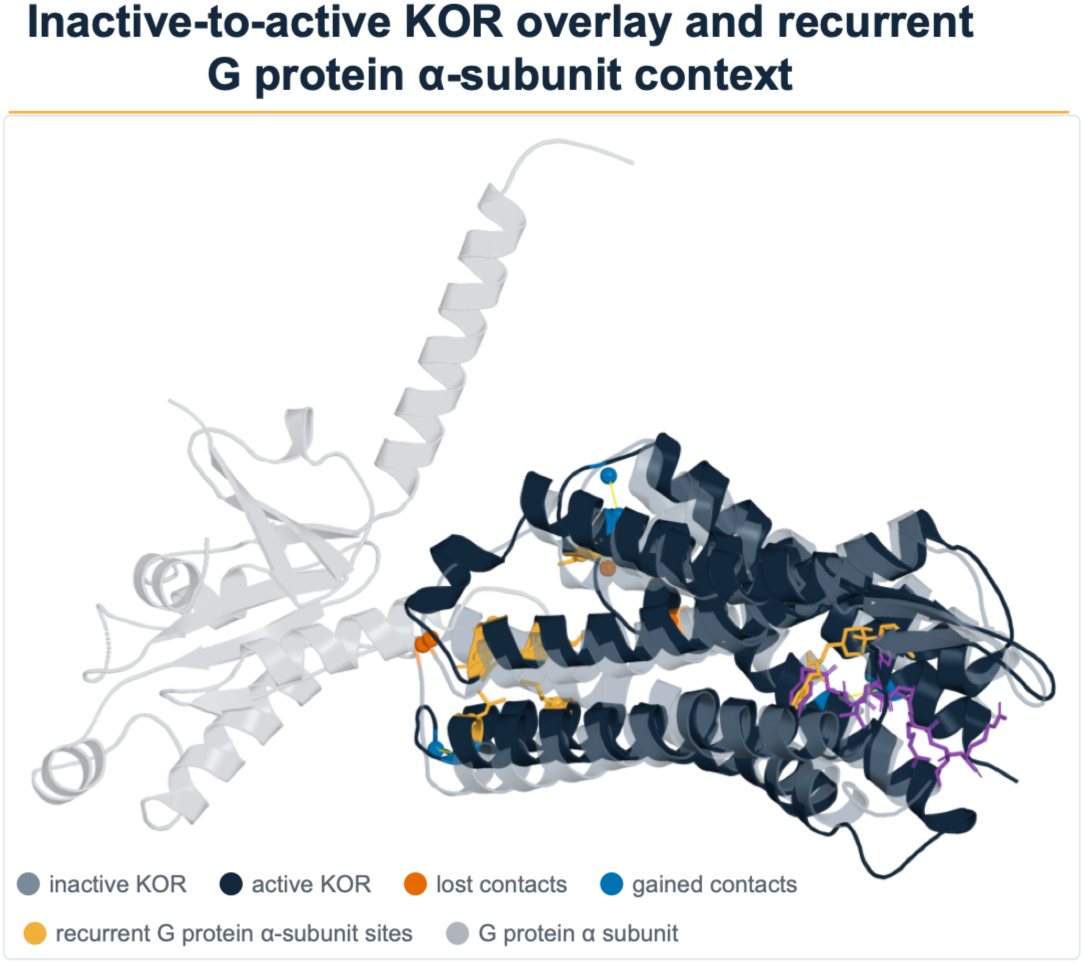
Three-dimensional context for the state positive control and recurrent Gα interface. Inactive-like KOR 6VI4 (slate) is overlaid with active, G protein-bound KOR 7Y1F (dark navy); Gα is light gray. Orange and blue spheres and dashed links show representative contacts lost and gained, respectively, between the two receptor graphs. Gold spheres on 7Y1F mark six recurrent exact-BW Gα-interface positions: 5x65 in 66 of 66 G protein maps; 3x53 and 3x54 in 65; and 2x39, 3x50, and 6x33 in 64. Co-resolved ligands are shown as gold and purple sticks.

**Table 2.** Literature-directed state positive control, inactive-like minus active-partner-bound.

| Outcome | Adjusted difference | Difference as % of active-partner-bound mean (95% CI) | $q$ | Model $n$ | Full endpoint leave-one-out sign consistency |
| --- | --- | --- | --- | --- | --- |
| Endpoint-mean contact Jaccard similarity | -0.02323 | -2.49% (-3.74%, -1.24%) | 0.00152 | 65 | 100% |
| Endpoint-mean rewired-contact count | +25.50 | +34.84% (+16.78%, +52.90%) | 0.00156 | 65 | 100% |
| Endpoint-mean changed-adjacent count | +10.70 | +34.78% (+18.64%, +50.92%) | 0.00116 | 59 | 100% |
| Endpoint-mean changed-direct count | +2.94 | +25.05% (+9.87%, +40.24%) | 0.00686 | 59 | 100% |
| Endpoint-mean shared-contact distance RMS deviation | +0.0480 Å | +15.15% (+4.84%, +25.45%) | 0.0151 | 65 | 100% |
| Exact-BW contact-pair count | -17.63 | -1.67% (-2.97%, -0.38%) | 0.0326 | 65 | 100% |

These directions are consistent with activation-linked receptor-core and intracellular-surface differences reported in opioid-receptor structures.^5–10^ Because the cohort includes structures that founded those models, this is a literature-directed workflow positive control. It is not independent replication or confirmation of a particular activation mechanism.

### A cohort-wide spatial atlas broadens quantitative comparison

Selected-ligand exact-BW sites were resolved in 77 of 84 independent maps, 91.7%, permitting all 2,926 comparisons among those maps. Under the Cα 8 Å graph, the pooled selected-ligand-centered static wiring was 13.39% direct, 32.53% adjacent, and 54.08% connected-distal. An estimand is the precisely specified quantity targeted by an analysis. The primary pair estimand induced each structure’s contact graph on the exact-BW nodes available in both structures. Among 239,736 shared-node-restricted changed contacts, 15.28% were direct, 40.13% adjacent, and 44.59% connected-distal. The disconnected fraction was zero because every selected-ligand-positive receptor graph was connected under this definition. A full-edge-set sensitivity contained 281,166 changed contacts and gave 13.19%, 36.31%, and 50.49%, respectively. Of those 281,166 full-edge changed-contact records, 41,430, or 14.74%, had at least one endpoint outside the pair’s shared-node intersection. Here, distal means connected-distal rather than an unresolved path.

Prior MOR studies have reported activation-linked conformational propagation and experimentally supported allosteric modulation.^6, 14^ In the BMS-986187 system, long-range communication pathways connecting the positive allosteric modulator site with DAMGO and the G-protein interface were inferred from molecular-dynamics trajectories by information-theory analysis.^15^ The present analysis adds a harmonized opioid-family measurement: exact-BW, direction-aware assignments for gained, lost, and conserved contacts across every eligible pair, with unresolved sites kept missing and source records provided for every summary. Pair ordering supplies computational source and destination labels only; it does not imply biological direction, signal propagation, or causality (Figure 9). The coordinate-defined direct, adjacent, and connected-distal classes around dynorphin and MPAM-15 in 9L60 are illustrated in Figure 10.

**Figure 9.**
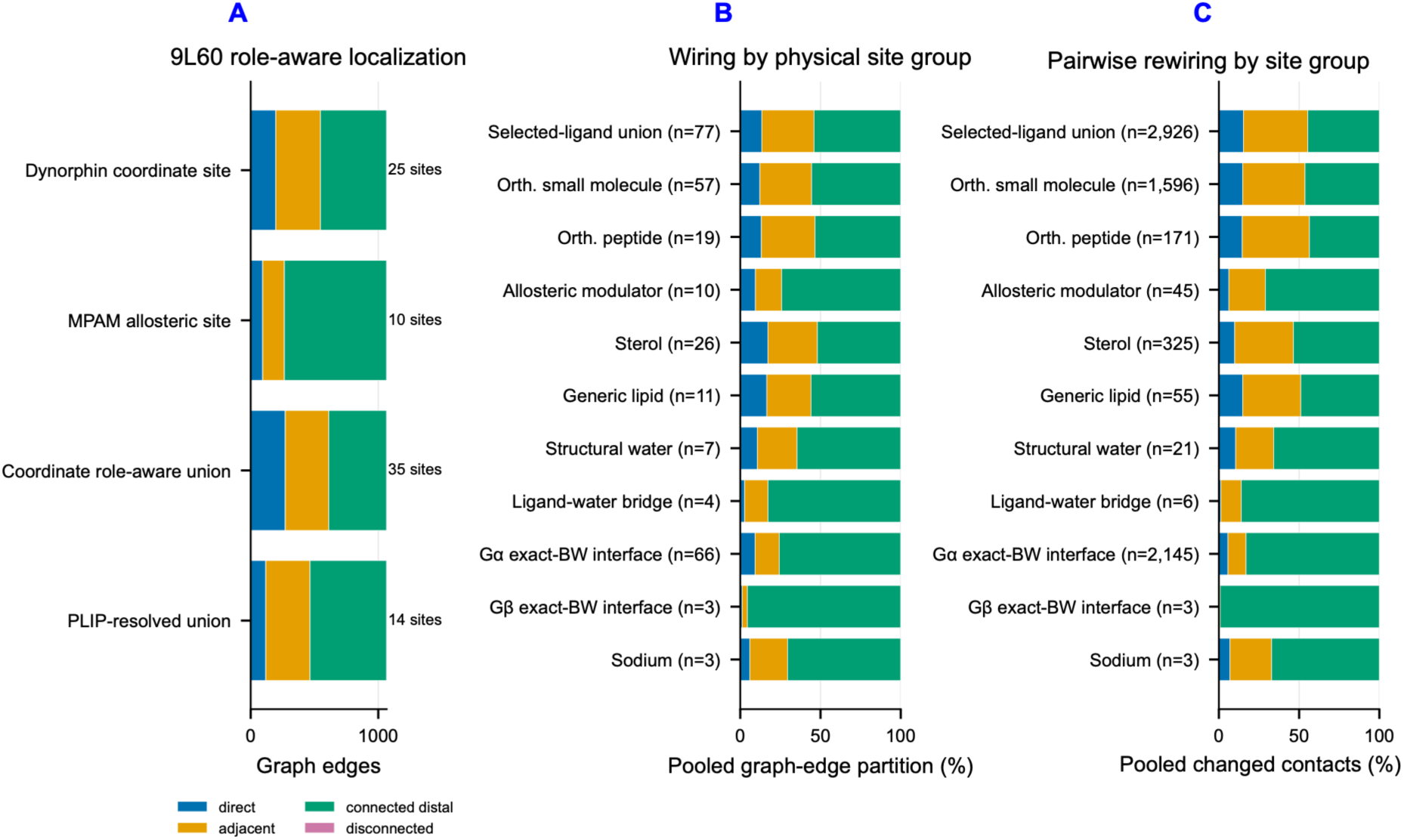
Coordinate role-aware direct, adjacent, connected-distal, and disconnected wiring. (A) KOR 9L60 contains a 25-site dynorphin footprint and a 10-site MPAM allosteric footprint; the coordinate union contains 35 exact-BW sites and partitions 1,066 edges into 271 direct, 341 adjacent, 454 connected-distal, and 0 disconnected. Its PLIP-resolved union contains 14 sites because the allosteric target is unresolved in that layer. (B,C) Pooled wiring and shared-node rewiring percentages keep the four classes disjoint. Direct/adjacent/connected-distal percentages are 10.73/24.66/64.61 and 10.44/23.67/65.89 for structural water; 2.76/14.57/82.67 and 1.27/12.72/86.01 for ligand-water bridges; 17.25/30.82/51.93 and 9.88/36.61/53.51 for sterol; 16.56/27.47/55.97 and 14.87/36.13/49.00 for generic lipid; 9.45/16.39/74.16 and 6.22/22.79/70.99 for allosteric sites; and 9.21/15.12/75.67 and 5.67/11.27/83.06 for Gα. Disconnected is explicitly 0 for these graphs. The allosteric group contains 10 maps/45 both-positive pairs; the selected-ligand union contains 77/2,926.

**Figure 10.**
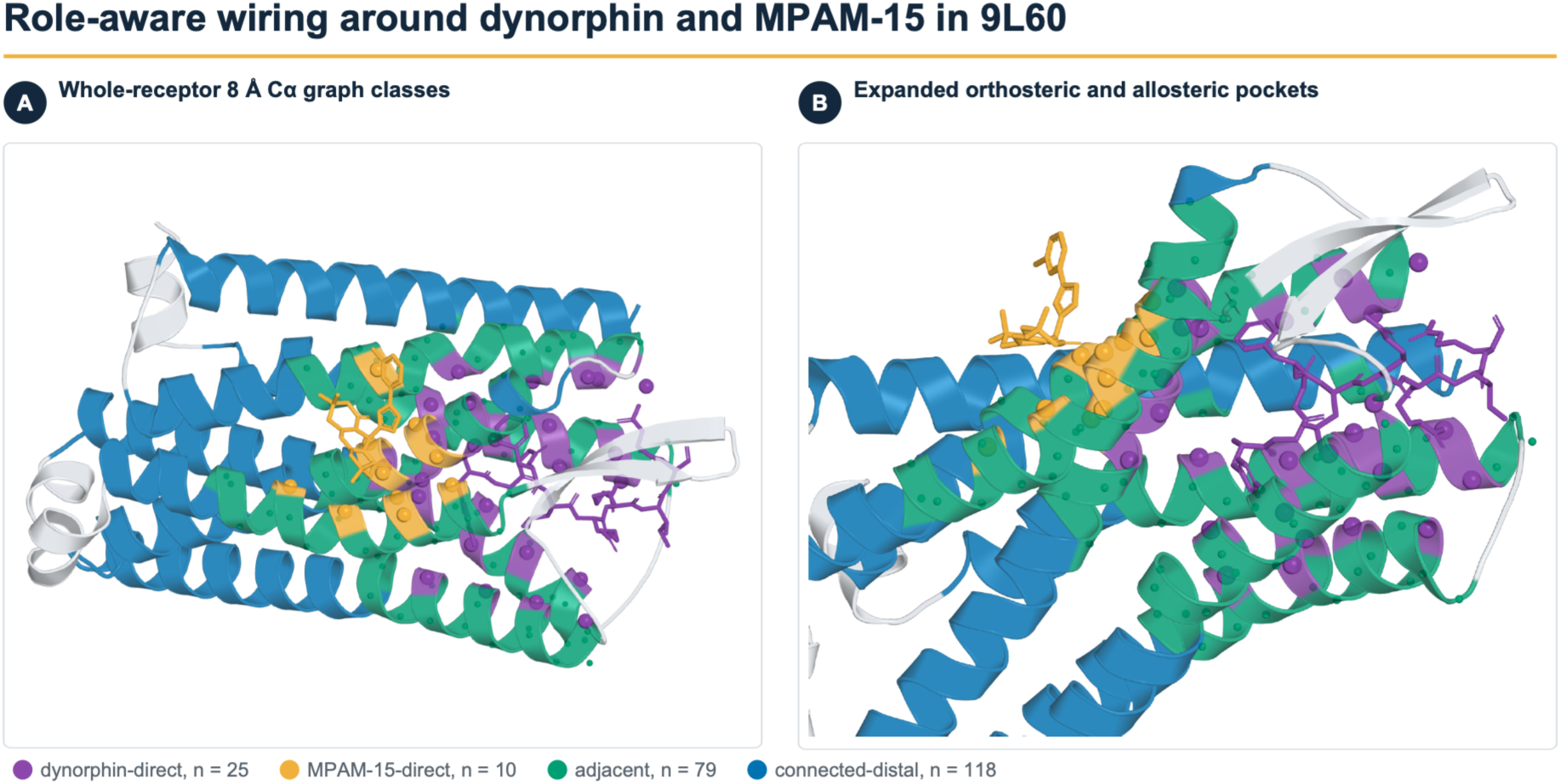
Three-dimensional role-aware wiring around dynorphin and MPAM-15 in KOR 9L60. (A) Whole-receptor view of the mutually exclusive node classes in the 8 Å Cα contact graph. (B) Expanded view of the orthosteric and allosteric pockets. Dynorphin and its 25 direct receptor sites are purple; MPAM-15 and its 10 direct sites are gold; 79 adjacent sites are teal; and 118 connected-distal sites are blue.

### Component prevalence and percentage-scale structural context

In this section, receptor-contacting means that at least one component nonhydrogen atom lies within 4.5 Å of the designated receptor. A structural water is a deposited receptor-proximal water that contacts at least two protein residues at the stated polar cutoff. Receptor-contacting sterol occurs in 26/84 maps, 31.0%; generic lipid in 11/84, 13.1%; deposited receptor-proximal water in 9/84, 10.7%; two-protein-residue structural water in 7/84, 8.3%; a resolved heterotrimeric G-protein assembly in 66/84, 78.6%; a receptor-contacting allosteric modulator in 10/84, 11.9%; orthosteric peptide in 19/76 resolved-modality maps, 25.0%; and orthosteric-plus-allosteric binding in 9/81 resolved binding-mode maps, 11.1%. Within the 66 Gα-positive maps, 11, or 16.7%, contain direct receptor/Gβ contact at 4.5 Å. The PIP₂-like analysis category and β-arrestin each occur only in 9ZZO. Receptor-contacting sodium occurs in 3/84 maps, and a selected-ligand/water/receptor bridge occurs in 4/84 (Figure 11). Representative deposited component and partner-interface relationships are shown in Figure 12.

**Figure 11.**
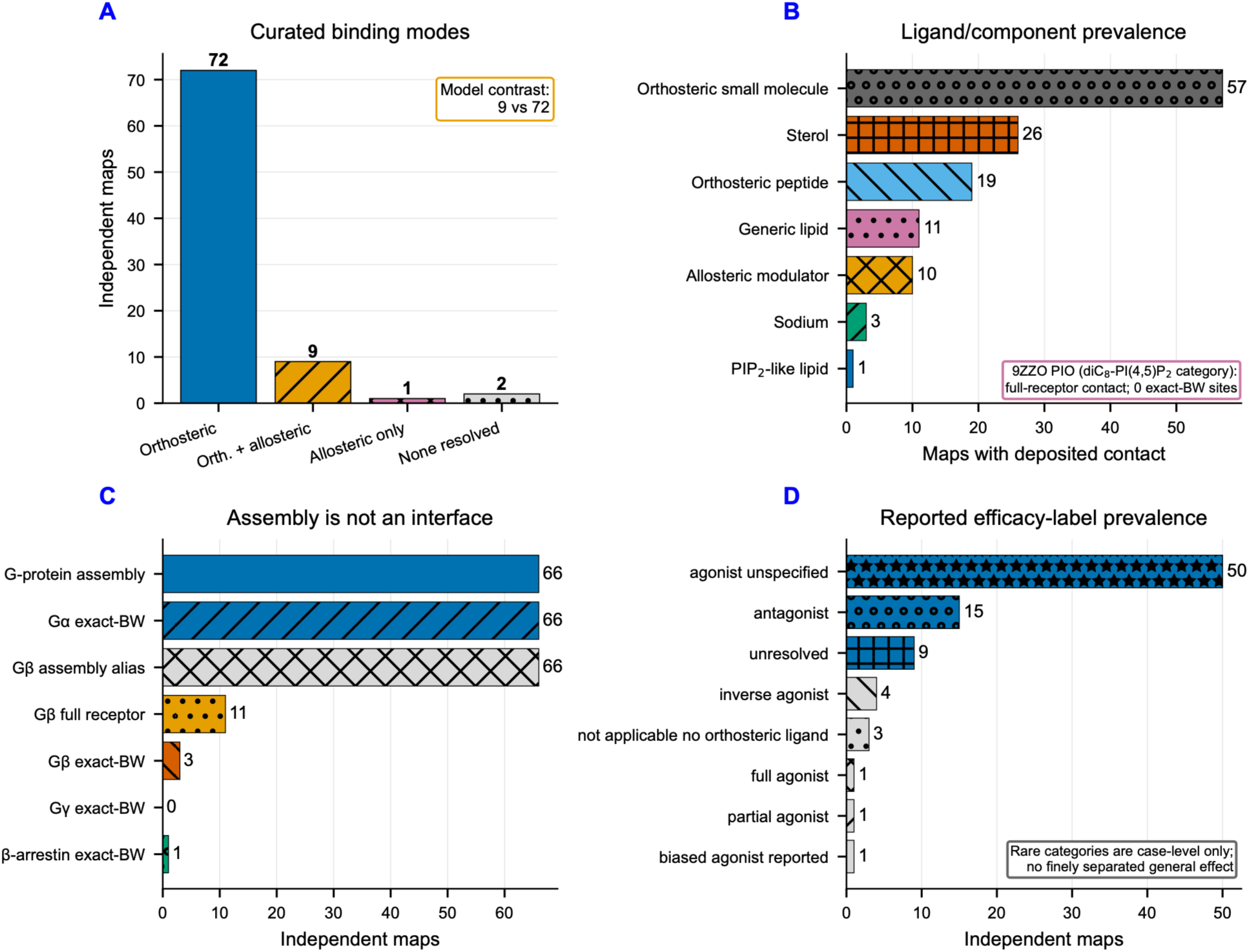
Curated ligand, component, and protein-partner context. (A) Curated binding modes comprise 72 orthosteric, 9 orthosteric-plus-allosteric, 1 allosteric-only, and 2 none-resolved maps. Physical allosteric contacts occur in 10 maps; the inferential comparison is 9 versus 72. (B) Component prevalence separates assembly, full-receptor contact, and exact-BW contact. 9ZZO is an engineered KOR/V2R-tail/β-arrestin 1 assembly prepared with short-chain diC₈-PI(4,5)P₂; PDB ligand PIO is assigned to the "PIP₂-like lipid" analysis category. PIO makes a full-receptor contact but has no exact-BW core site within 5 Å; PIO, MPAM, and 9L60 dynorphin records are included in the figure source data. (C) Gβ/Gγ assembly co-presence aliases the 66-map G-protein heterotrimer vector and is not a Gβγ-specific exposure; Gβ contacts the full receptor in 11 maps but reaches an exact-BW interface in only 3, while Gγ reaches none. β-arrestin has one exact-BW-positive map. (D) Efficacy labels are descriptive: 50 agonist-unspecified and 15 antagonist maps dominate, while inverse agonist (4), biased agonist (1), partial agonist (1), and full agonist (1) are too sparse for finely separated general effects.

**Figure 12.**
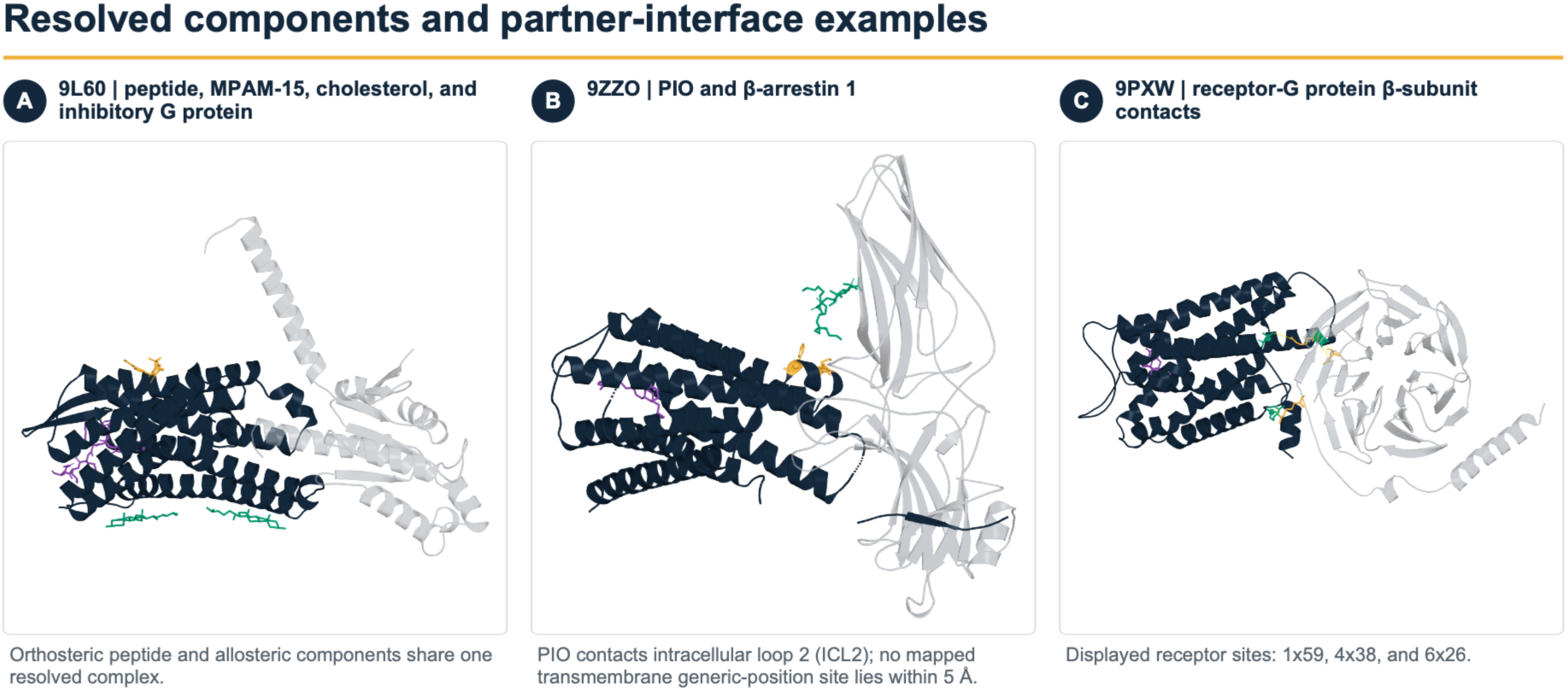
Three-dimensional examples of resolved components and protein-partner interfaces. (A) KOR 9L60 with orthosteric dynorphin (purple), MPAM-15 (gold), two cholesterol molecules (teal), and Gα (light gray). (B) Engineered KOR/V2R-tail/β-arrestin 1 complex 9ZZO with MP1104 (purple) and PIO (teal; defined in Figure 4). Gold spheres mark the PIO-contacting intracellular-loop-2 residues Lys165 and Phe169; PIO has no mapped transmembrane exact-BW site within 5 Å. (C) MOR-G protein complex 9PXW with naloxone (purple). Teal receptor sites 1x59, 4x38, and 6x26 contact the gold Gβ residues Arg46, Arg52, and Asp333, respectively, within the 4.5 Å heavy-atom cutoff; Gβ is light gray.

Site-centered localization describes where the receptor graph lies relative to each resolved site. For sterol sites, static wiring is 17.25% direct, 30.82% adjacent, and 51.93% connected-distal; shared-node changed contacts are 9.88%, 36.61%, and 53.51%, respectively. Generic-lipid sites give 16.56/27.47/55.97% static wiring and 14.87/36.13/49.00% shared-node changed contacts. Allosteric-modulator sites give 9.45/16.39/74.16% and 6.22/22.79/70.99%. Gα interfaces give 9.21/15.12/75.67% and 5.67/11.27/83.06%. These distributions answer where contacts occur relative to a deposited site. They do not identify an effect caused by that component.

An exposed map contains the named feature under its operational definition; a reference map does not. A bootstrap interval is obtained by repeatedly resampling the observed map summaries with replacement. Unadjusted spatial composition was evaluated only in the 77 maps with a resolved selected-ligand site (Table 3; SI Table S5). Sterol-contacting maps had a 14.92% higher mean changed-direct fraction and a 9.20% lower changed connected-distal fraction than reference maps. Structural-water-positive maps had a 15.99% lower changed-direct fraction and an 18.03% higher changed connected-distal fraction; deposited-water-positive maps showed the same direction at -13.65% and +12.18%. Generic-lipid-contact maps had a 17.03% lower changed-direct fraction and a 15.75% higher changed connected-distal fraction. Resolved G-protein maps had a 16.42% higher changed-direct fraction and a 13.85% lower changed connected-distal fraction. Allosteric-contact maps had a 17.70% higher changed-direct fraction and a 13.56% lower changed connected-distal fraction. The unadjusted intervals reported in SI Table S5 use 5,000 structure-level resamples within exposure groups. These unadjusted percentages describe deposited-model composition and possible magnitude, not an isolated component effect.

**Table 3.** Percentage-scale component context and evidential boundary.

| Context | Full-scope prevalence | Unadjusted spatial n (E/R) | Observed percentage pattern | Minimum-p multivariable-adjusted result, exploratory display | Interpretation |
| --- | --- | --- | --- | --- | --- |
| Receptor-contacting sterol | 26/84 (31.0%) | 23/54 | Changed direct +14.92%; connected-distal -9.20% | Endpoint-mean shared-contact distance RMS deviation -7.28% (95% CI -17.56%, +3.01%); $q = 0.6970$ | Descriptive; adjusted direction unclear |
| Receptor-contacting generic lipid | 11/84 (13.1%) | 9/68 | Changed direct -17.03%; connected-distal +15.75% | Endpoint-mean shared-contact distance RMS deviation -5.76% (-13.39%, +1.87%); $q = 0.6970$ | Limited overlap |
| Deposited receptor-proximal water | 9/84 (10.7%) | 9/68 | Changed direct -13.65%; connected-distal +12.18% | Endpoint-mean shared-contact distance RMS deviation +3.42% (-3.69%, +10.52%); $q = 0.7251$ | Coverage-limited |
| Two-protein-residue structural water | 7/84 (8.3%) | 7/70 | Changed direct -15.99%; connected-distal +18.03% | Endpoint-mean shared-contact distance RMS deviation +7.67% (-0.21%, +15.55%); $q = 0.4511$ | Exploratory, replication-limited |
| Resolved G-protein assembly | 66/84 (78.6%) | 60/17 | Changed direct +16.42%; connected-distal -13.85% | Endpoint-mean shared-contact distance RMS deviation +1.58% (-14.08%, +17.25%); $q = 0.9737$ | Partner/state/construct bundle |
| Direct Gβ contact within Gα | 11/66 (16.7%) | Not in unadjusted spatial table | No separate assembly exposure | Exact-BW contact-pair count +0.31% (-1.22%, +1.84%); $q = 0.9737$ | No association detected |
| Orthosteric peptide | 19/76 (25.0%) | 19/57 | Static direct +24.28%; connected-distal -10.17% | Endpoint-mean shared-contact distance RMS deviation -6.18% (-10.82%, -1.54%); $q = 0.3165$ | Bounded exploratory hypothesis |
| Orthosteric plus allosteric | 9/81 (11.1%) | 9/67 | Changed direct +20.05%; connected-distal -15.84% | Endpoint-mean contact Jaccard similarity +0.47% (-0.24%, +1.19%); $q = 0.6970$ | Publication-limited |
| Receptor-contacting allosteric modulator | 10/84 (11.9%) | 10/67 | Changed direct +17.70%; connected-distal -13.56% | Not a separate primary contrast | Descriptive physical-contact context |

E/R denotes exposed/reference.

One multivariable-adjusted ordinary least-squares model with type 3 heteroskedasticity-consistent (HC3) covariance was defined for each combination of eight deduplicated biological contrasts and four outcomes, giving 32 tests in one global Benjamini-Hochberg family. None had *q* < 0.05. For compact display, Table 3 shows the outcome with the smallest nominal *p* value among each contrast’s four defined outcomes; this is an exploratory selection rule, and all 32 rows remain in the multiplicity family. For peptide versus small molecule, the selected outcome was endpoint-mean shared-contact distance RMS deviation, which was 0.02082 Å lower, or 6.18% of the observed small-molecule mean (95% CI, -10.82% to -1.54%; *p* = 0.00989 ; *q* = 0.3165). The adjusted coefficient remained negative in every structure and publication leave-one-out refit. Eighteen exact-stratum matched pairs had a mean peptide-minus-small-molecule difference of -0.01889 Å. Within-publication contrasts were available in five discordant publications and agreed with the adjusted direction in three. A discordant publication is one that contains maps from both contrast levels. These checks support a testable, multiplicity-qualified hypothesis, not a discovery-level effect. For structural-water-positive maps, the minimum-*p* outcome under the shared-node primary estimand was endpoint-mean shared-contact distance RMS deviation, estimated as 7.67% higher (95% CI, -0.21% to +15.55%; *p* = 0.0564 ; *q* = 0.4511). Only four matched pairs and one discordant publication support that contrast. The within-Gα Gβ-contact estimate was near zero: +0.31% exact-BW contacts (95% CI, -1.22% to +1.84%; *q* = 0.9737).

### Deposited-water estimands form structural and ligand-contact branches

With a deposited water-oxygen fractional occupancy of at least 0.7, receptor proximity at most 5.0 Å, and a 3.5 Å nitrogen-or-oxygen (N/O) polar-contact cutoff, 137 receptor-proximal deposited waters occurred in 9 maps (Figure 13; Table 4). Eighty-five waters in 8 maps contacted at least one exact-BW receptor residue. The structural branch contained 61 waters in 7 maps that contacted at least two protein residues; 52 contacted at least two designated-receptor residues, 44 were members of the 61-water core and contacted at least one exact-BW residue, and 38 were core waters that bridged at least two exact-BW residues. A separate ligand-contact branch contained 10 selected-ligand/water/receptor waters in 4 maps. Three of those 10 also belonged to the 38-water core exact-BW bridge set, whereas 7 were outside the two-protein-residue structural core. All 137 retained waters had deposited occupancy 1.0, so the tested occupancy thresholds did not change membership.

**Figure 13.**
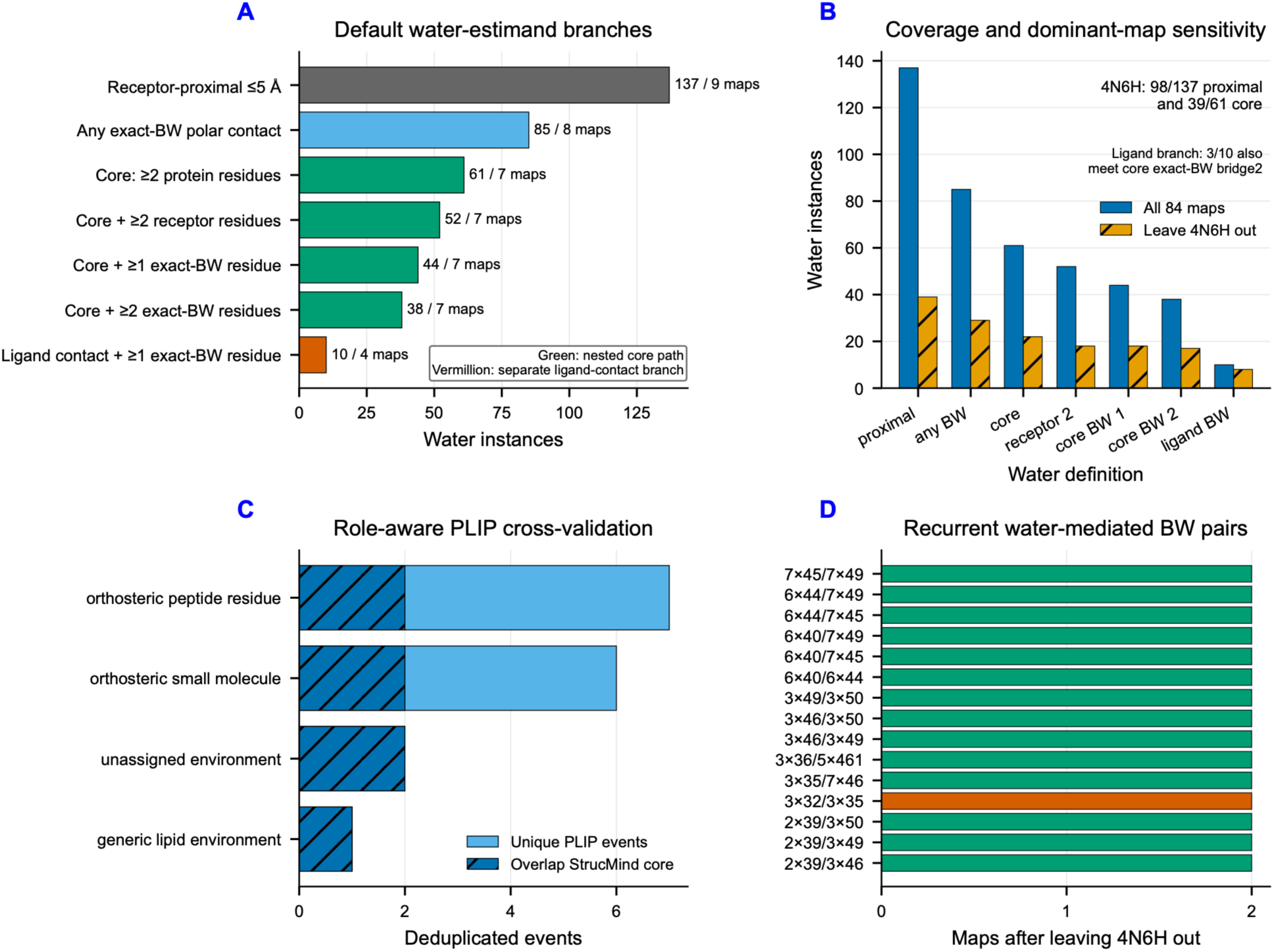
Defined water-estimand branches and role-aware cross-validation. At occupancy ≥0.7 and polar distance ≤3.5 Å, 137 receptor-proximal waters occur in 9 maps. The nested core path contains 61 waters with at least two protein-residue contacts, 52 with at least two designated-receptor-residue contacts, 44 core waters touching at least one exact-BW residue, and 38 core waters bridging at least two exact-BW residues. A separate receptor-contact branch contains 85 waters touching at least one exact-BW residue. The ligand-contact branch contains 10 ligand-water-receptor bridges in 4 maps; all 10 touch an exact-BW residue, but only 3 also belong to the 38-water core exact-BW bridge set. Excluding 4N6H leaves 39 proximal, 29 any-exact-BW-contact, 22 core, 17 core exact-BW bridge, and 8 ligand-bridge waters. PLIP events are role-aware and deduplicate to 16 structure-water-BW events; they are secondary cross-validation, not a ligand-effect matrix. The 3×32/3×35 water-mediated pair occurs in 3/84 processed maps, 3.57%, and 3/7 exact-BW-bridge-positive maps, 42.86%; it spans DOR 4N6H, KOR 4DJH, and NOP 4EA3 and remains in two maps after 4N6H exclusion.

**Table 4.** Default deposited-water estimands and branch overlap.

| Group and set | Definition | Water instances | Positive maps | % of 137 root waters |
| --- | --- | --- | --- | --- |
| Root, $P$ | Receptor-proximal deposited water | 137 | 9 | 100.0% |
| Exact-BW branch, $B_{any}$ | $P$ with at least one exact-BW polar partner | 85 | 8 | 62.0% |
| Structural branch, $C$ | $P$ with at least two protein-residue polar partners | 61 | 7 | 44.5% |
| Structural branch, $R_2$ | $C$ with at least two designated-receptor partners | 52 | 7 | 38.0% |
| Structural plus exact BW, $C_{B1}$ | $C$ with at least one exact-BW partner | 44 | 7 | 32.1% |
| Structural plus exact BW, $C_{B2}$ | $C$ with at least two exact-BW partners | 38 | 7 | 27.7% |
| Ligand-contact branch, $L$ | $P$ with selected-ligand and exact-BW receptor polar partners | 10 | 4 | 7.3% |
| Branch overlap | $L \cap C_{B2}$ | 3 | 2 | 2.2% |

Seven of the 10 ligand-contact waters are outside $C$ . Thus, $L$ is not a nested continuation of the structural branch.

Relative to resolved structural-water sites, static wiring is 10.73% direct, 24.66% adjacent, and 64.61% connected-distal; shared-node changed contacts are 10.44%, 23.67%, and 65.89%. The four ligand-bridge-positive maps are still more distal by this graph definition: static wiring is 2.76/14.57/82.67%, and shared-node changed contacts are 1.27/12.72/86.01%, for direct/adjacent/connected-distal strata. The multivariable structural-water contrast instead selected shared-contact RMS distance as its minimum-*p* outcome: +7.67% (95% CI, -0.21% to +15.55%; *p* = 0.0564; *q* = 0.4511). These distributions and the adjusted estimate nominate a distal structural context for follow-up, but they do not distinguish a causal hydration effect from resolution, method, state, construct, or publication selection. The seven-map exposure, one discordant publication, and global *q* value preclude a cohort-general hydration claim.

At a 1.0 Å coordinate radius, complete linkage, which requires every within-cluster separation to be at most the radius, and single linkage, which joins waters connected by chains of within-radius links, each yielded six recurrent exact-BW-bridging water clusters across at least two maps; each yielded three after excluding 4N6H. Topological recurrence, defined as the same unordered exact-BW residue pair being bridged in at least two independent maps, identified 25 recurrent BW pairs, of which 15 remained after excluding 4N6H. The 3×32/3×35 bridge occurred in 3/84 maps overall, 3.57%, and 3/7 exact-BW-bridge-positive maps, 42.86%, spanning DOR 4N6H, KOR 4DJH, and NOP 4EA3; the KOR/NOP recurrence persists without 4N6H. The 84-map denominator describes cohort-wide recurrence, whereas the 7-map denominator conditions on maps capable of contributing a *C*_B2_ topology. This topology is an exploratory candidate for targeted simulation or mutagenesis, not evidence of a conserved dynamic water wire. Figure 14 shows the recurring deposited-water geometry in the three contributing receptor subtypes.

**Figure 14.**
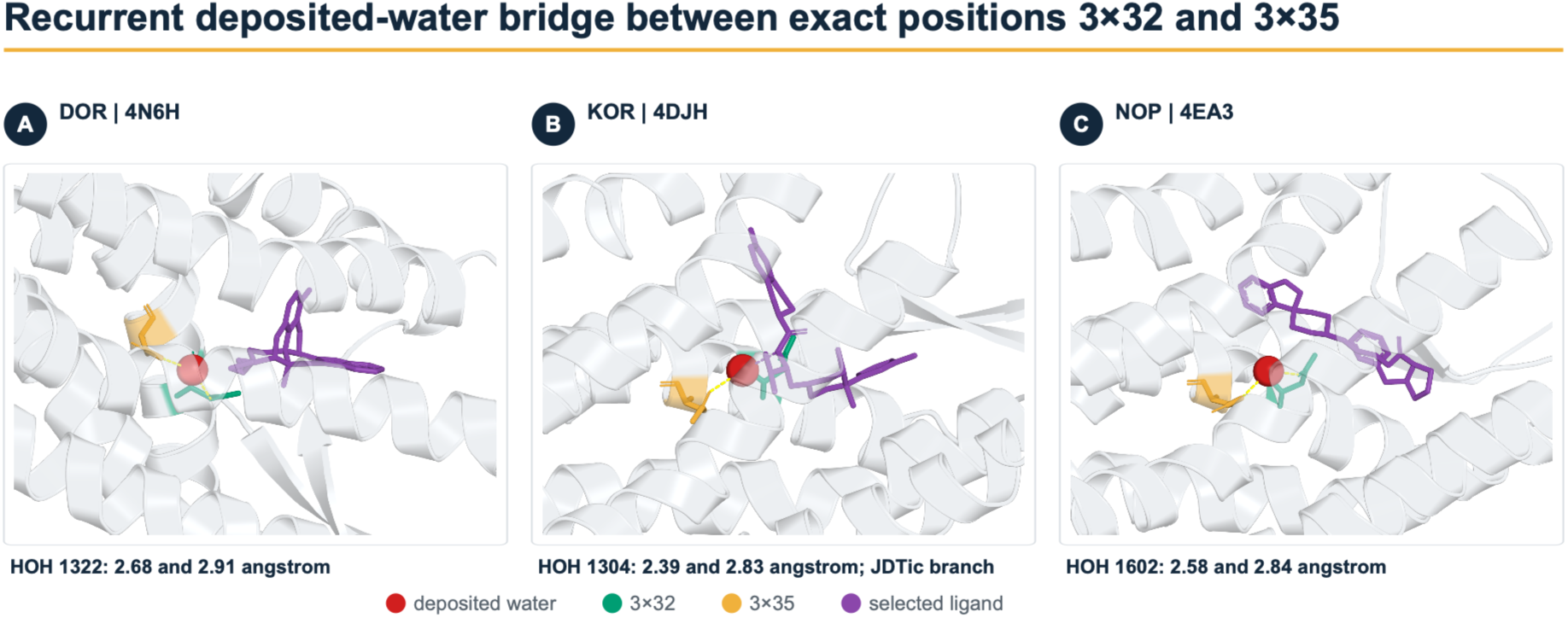
Three-dimensional context for the recurrent deposited-water bridge between exact positions 3×32 and 3×35. (A) DOR 4N6H, water HOH 1322, with water-to-residue distances of 2.68 and 2.91 Å. (B) KOR 4DJH, water HOH 1304, with distances of 2.39 and 2.83 Å. (C) NOP 4EA3, water HOH 1602, with distances of 2.58 and 2.84 Å. Exact positions 3×32 and 3×35 are teal and gold, respectively; deposited waters are red spheres; selected ligands are purple; protein-water links are blue; and the JDTic-water link in 4DJH is purple. Only 4DJH belongs to the selected-ligand water branch. The geometry recurs in 3 of 84 processed maps, 3.57%, and 3 of 7 exact-BW-bridge-positive maps, 42.86

DOR 4N6H contributes 71.5% of receptor-proximal waters and 63.9% of structural-core waters. Removing it leaves 39 receptor-proximal waters, 29 with any exact-BW contact, 22 structural-core waters, and 17 core exact-BW bridges. PLIP supplies 16 unique structure/water/BW events, of which 7, or 43.8%, overlap the two-protein-residue core; PLIP and the coordinate-defined water analysis are related but nonidentical estimands. Coordinate-defined recurrent clusters were contributed only by X-ray structures. Exact-BW topological recurrence additionally included a mixed-method DOR pair, X-ray 4N6H and cryogenic electron microscopy (cryo-EM) 9YDP, at 3×54/5×64. Primary studies establish conserved GPCR water networks,^17^ a DOR sodium pocket,^18^ and water-mediated MOR sodium-site engagement^19^ in particular experimental systems. The present result is a static deposited-water coordinate and topology atlas. It does not estimate solution hydration, residence time, energetics, ligand-efficacy effects, or a causal dynamic water network.

### Protein interfaces, sparse components, and activity classes

Resolved G-protein assembly is present in 66/84 maps. Because the Gα, Gβ, and Gγ assembly vectors are identical to the resolved G-protein vector, comparing assembly presence would only compare coupled with uncoupled deposited structures. Direct receptor/Gβ contact is narrower: 11 maps at 4.5 Å and 15 at 5.0 Å, with three maps contributing an exact-BW Gβ interface. No Gγ residue met the primary 4.5 Å receptor-contact criterion. The defensible Gβ-specific question is the within-Gα contact contrast, for which no association was detected.

β-Arrestin is present in one map, 9ZZO. In that engineered KOR/V2R-tail preparation, short-chain PI(4,5)P₂ (PDB ligand PIO), β-arrestin 1, Fab30, an antibody fragment, Nb32, a nanobody used to stabilize the complex, and other stabilizing elements form one inseparable exposure. PIO contacts the full receptor within 5.0 Å but does not contact an exact-BW core residue within 5.0 Å. The atlas therefore records the published engineered preparation rather than providing an independent β-arrestin or PIP₂ comparison.^12^

The ten receptor-contacting allosteric-modulator maps are 8K9L, 8Y71, 8Y72, 8Y73, 9L60, 9PU5, 9XC6, 9XDQ, 9XDR, and 9XF4. Nine also contain an orthosteric ligand, and one is allosteric-only. These maps include the established MPAM-15, BMS-986122, and BMS-986187 systems.^14–16^ Their percentage profiles are useful structural context, but the adjusted binding-mode contrast has only one discordant publication and cannot establish a generic allosteric-modulator effect. The same boundary applies across mechanisms established in specific systems: sodium,^18^ phosphoinositide and arrestin,^12, 13, 22^ and cholesterol or lipid context.^15, 20, 21^ A mechanism established in one system is not a cohort-general association.

Activity labels are too imbalanced for finely separated efficacy claims. Among 84 maps, 50 are unspecified agonists, 15 antagonists, 4 inverse agonists, 1 biased agonist, 1 partial agonist, 1 full agonist, 3 have no orthosteric ligand, and 9 remain unresolved. These are source-reported pharmacological labels retained without cross-assay normalization; no common response threshold was imposed. These labels are preserved for structure-specific queries, but the cohort cannot identify general full, partial, inverse, or biased efficacy effects. Recent MOR dynamics and DOR partial-agonism studies provide system-specific mechanistic evidence; the present deposited-coordinate analysis does not substitute for them.^11, 23^

### Claim classification against prior literature

The results separate into four claim levels. **Literature-directed positive control** comprises only the state-associated result, whose direction is consistent with inactive, active, and transducer-bound structures but is not independent replication. **Dataset-level quantitative contribution** comprises the curated 86-structure/84-map resource, 3,655 all-to-all comparisons, exact-BW contact records, curated ligand and component roles, spatial direct/adjacent/connected-distal framework, percentage-scale reporting, and deposited-water recurrence atlas. The water, sodium, lipid, G-protein, and arrestin observations are qualitative context or structure-specific recapitulations, not confirmation of their mechanisms. **Exploratory hypotheses** comprise the peptide-associated RMS geometry and recurrent 3×32/3×35 water topology. **Not identified as cohort-general effects** comprises hydration, cholesterol, generic lipid, PIP₂, β-arrestin, sodium, G-protein assembly, Gβ contact, generic allosteric modulation, and finely separated activity classes. Failure to identify a general association does not imply that a component is biologically inactive.

### Study Scope and Interpretation Boundaries

Every input is a deposited static model subject to construct design, stabilization, resolution, component selection, and publication bias. Component presence is not experimental addition, and apparent absence can mean unresolved or not modeled. Pairwise outcomes reuse structures; endpoint means reduce but do not eliminate dependence. Unadjusted percentage contrasts show scale but can reflect state, subtype, method, mapping coverage, exact-BW mapped-residue count, ligand, partner, or publication context. Multivariable-adjusted models control identified covariates, not unmeasured selection. Across the 84 independent-map endpoint means under the primary shared-node estimand, contact Jaccard similarity and rewired-contact count are almost perfectly redundant (Pearson *r* = −0.997460, where *r* is the linear-correlation coefficient) and are not independent support. Static coordinates cannot establish kinetics, thermodynamics, solution occupancy, conformational populations, information-flow direction, signaling efficacy, clinical activity, or causality.

## Conclusions

### From a Fragmented Structural Record to a Position-Exact Design Map

This study transforms the deposited human opioid receptor structural record from a collection of individually interpreted structures into a common, position-exact comparison space. Exact Ballesteros-Weinstein mapping, shared-node pairwise analysis, and complete single-structure and all-pair enumeration make it possible to distinguish conserved contacts from context-dependent wiring and rewiring without relying on selected structural exemplars. Recovery of established state-associated organization provides a literature-directed positive control, demonstrating that the framework recognizes known opioid receptor biology. The central advance, however, is the ability to localize structural conservation and variation relative to orthosteric ligands, allosteric ligands, deposited waters, molecular components, and transducer interfaces within one internally consistent atlas.

The ligand-facing analysis identifies a recurrent coordinate-defined direct-contact spine that includes Asp3×32, Tyr3×33, Met3×36, Trp6×48, and Tyr7×42 across representative structures of all four receptor subtypes. These positions provide a family-wide reference for receptor engagement. They should not be interpreted as experimentally validated affinity or potency determinants, but they provide a rational starting set of interactions to preserve while testing chemical variation elsewhere. The biological picture that follows is therefore not one of four unrelated receptors. It is a shared structural scaffold whose local neighborhoods and distal contact organization are reconfigured by receptor sequence, ligand geometry, conformational state, protein partner, and molecular environment.

### Residue-Level Routes toward Selectivity and Potency

The atlas supports a layered ligand-design strategy. The recurrent direct-contact spine can be treated as a conservative pharmacophoric framework, while subtype-variable positions within or beside the ligand-facing surface provide candidate selectivity vectors. Position 2×63 is a particularly informative example because it presents chemically distinct environments across the family: Asn in MOR, Val in KOR, Lys in DOR, and Asp in NOP. In the role-aware KOR 9L60 graph, the adjacent position 4×55 is likewise chemically distinct, with Ala in MOR, Gly in DOR, Ser in KOR, and Val in NOP. These positions offer concrete directions for matched substituent scans, stereochemical changes, peptide side-chain substitutions, and structure-guided extension of a conserved ligand core.

A practical small-molecule strategy would retain interactions with the recurrent direct-contact spine while varying one substituent at a time toward positions such as 2×63 or an adjacent subtype-variable neighborhood such as 4×55. Such analogs should be tested across MOR, KOR, DOR, and NOP rather than only against the intended receptor. This would distinguish increased target engagement from loss of activity at competing subtypes and would reveal whether selectivity results from one dominant interaction or the cooperative effect of several smaller differences.

For peptides and peptidomimetics, the dynorphin footprint in 9L60 provides an explicit extended engagement surface spanning TM2, TM3, TM5, TM6, and TM7. Side-chain scanning, truncation, cyclization, backbone constraint, N-methylation, and peptidomimetic substitution can therefore be directed toward defined portions of that footprint rather than applied empirically. The peptide-associated shared-contact geometry remains a multiplicity-qualified hypothesis, so it should motivate prospective experiments rather than be presented as a general rule that peptides possess a particular potency or efficacy advantage.

Potency optimization can be pursued through preservation of favorable direct interactions, ligand preorganization, reduction of unfavorable desolvation, and deliberate manipulation of water-mediated contacts. None of these outcomes is measured by the structural atlas itself. Affinity, residence time, potency, and efficacy must be established in standardized experiments. The value of the atlas is that it specifies which chemical changes should be tested and which receptor positions should be monitored when those properties are measured.

### One TM3 Axis Integrates Orthosteric, Allosteric, and Water-Aware Design

The most compelling integrative structural example centers on Asp3×32 and Asn3×35 of TM3. In KOR structure 9L60, orthosteric dynorphin A contacts Asp3×32, whereas the simultaneously bound allosteric modulator MPAM-15 contacts Asn3×35 and occupies a distinct TM2 to TM4 surface involving positions 2×52; 3×30, 3×31, 3×34, and 3×35; and 4×50, 4×54, 4×57, 4×58, and 4×61. This co-occupied structure provides an exact-residue template for separating orthosteric and allosteric design variables. Matched dynorphin and MPAM-15 analogs can first be used to determine how each site contributes to binding cooperativity and signaling under controlled conditions. The resulting geometry can then be evaluated for bitopic feasibility by varying linker length, rigidity, attachment point, and polarity while retaining the untethered parent ligands as controls. The structure demonstrates co-occupancy, but it does not establish that the two ligands can be joined productively.

The same TM3 pair participates in the strongest deposited-water candidate. A deposited water bridges Asp3×32 and Asn3×35 in DOR 4N6H, KOR 4DJH, and NOP 4EA3. In the KOR 4DJH structure, that water also contacts the antagonist JDTic. The recurrence in KOR and NOP persists after removal of the water-rich DOR structure, making the topology a particularly clear prospective test. It remains a static deposited-water observation, not an essential water, a dynamic water wire, or evidence of a hydration-mediated allosteric pathway. No corresponding MOR instance was identified in the deposited-water subset, and the contributing examples are limited to X-ray structures.

This topology nonetheless supports four distinct medicinal-chemistry experiments. A ligand can be designed to preserve the deposited bridge, recruit a water through an added donor or acceptor, displace the water with a steric or hydrophobic group, or replace its interactions with a ligand functional group that directly engages the Asp3×32 and Asn3×35 environment. Matched JDTic analogs that retain, remove, or reposition the water-facing vector would provide the cleanest initial test. Their evaluation should combine high-resolution structures, conservative perturbation of Asn3×35, explicit-solvent simulation, hydration free-energy analysis, binding affinity, kinetics, and receptor function. Concordance across these measurements would promote the topology from a static candidate to a water-aware design element. Discordance would be equally informative and should retire the proposed mechanism.

### Common Biology and Receptor-Specific Wiring

The atlas supports a model in which opioid receptors share a conserved activation scaffold but encode receptor identity within that scaffold. The common architecture includes the Asp3×32 recognition region, the Asp2×50-centered sodium pocket, the DRY motif, the PIF-like triad, CWxP, NPxxY, and the helix 8 anchor. These elements should not be viewed as independent modules. Asn3×35 belongs to the established sodium-pocket architecture while also appearing in the MPAM-15 site and the recurrent deposited-water topology. Asn7×49 contributes to the sodium-pocket region and initiates NPxxY. Such positions are structural crossroads in the deposited atlas, although their functional coupling remains to be tested.

Subtype differences also occur within otherwise conserved motifs. NOP contains Thr3×40 at a PIF-like position where MOR, KOR, and DOR contain Ile. DOR contains Ala6×49 within the CWxP region where the other subtypes contain Thr. Position 7×51 within the NPxxY span is Val in MOR and DOR but Ile in KOR and NOP. These substitutions illustrate that receptor-specific biology is not encoded only in peripheral loops or an outer selectivity shell. It is also embedded within the conserved helical machinery. They are candidate modulators of receptor-specific conformational behavior, not established determinants of signaling or pharmacology.

At the intracellular surface, recurrent Gα-contact positions include 2×39, Arg3×50, 3×53, 3×54, 5×65, and 6×33 across the receptor family. Arg3×50 connects the classical DRY motif directly to the repeatedly observed transducer-facing surface. In the 9L60 ligand-centered graph, these positions are connected-distal from the ligand union. This makes them experimentally addressable reporters for testing whether a ligand-dependent contact pattern is accompanied by a change at the transducer interface. Their graph relationship does not establish a physical signaling route or the direction of information transfer.

The single engineered KOR β-arrestin structure provides a more tentative extension of this picture. Its arrestin interface overlaps part of the recurrent Gα-facing surface at positions including 2×39, Arg3×50, 3×54, 5×65, and 6×33, while extending toward positions such as 5×68, 6×29, 6×32, and 7×56. This supports a testable common-core plus partner-specific-extension model. It does not establish a general mechanism of transducer selectivity or biased signaling because the arrestin context is represented by one engineered preparation.

The same structure also exposes an important boundary of the present exact-BW framework. Its PIP₂-like lipid contacts ICL2 residues Lys165 and Phe169 but does not contact the exact transmembrane BW core. Exact-BW mapping provides reproducibility across receptor subtypes, but it does not capture every loop, tail, or membrane-facing determinant. A next-generation atlas should therefore extend the current core analysis to harmonized intracellular-loop, extracellular-loop, helix 8, terminal-tail, and membrane-facing surfaces.

### From Structural Hypotheses to Measured Pharmacology

The atlas now supports a focused design-test-measure cycle. The first priority is to synthesize matched analogs that preserve the recurrent direct-contact spine while varying engagement of a single subtype-variable direct or adjacent position. These compounds should be paired with reciprocal receptor mutations and evaluated across the complete opioid receptor family. A position becomes a credible selectivity determinant only when ligand and receptor perturbations produce reciprocal, reproducible changes under the same assay conditions.

The second priority is to test the TM3 dual-site and water hypotheses directly. Orthosteric ligand, allosteric modulator, and bitopic candidate series should be evaluated with and without perturbation of the Asp3×32 and Asn3×35 environment. The recruit, preserve, displace, and replace water strategies should be compared in matched chemistry rather than inferred from unrelated ligands. Binding, kinetics, potency, efficacy, and transducer recruitment should be measured using shared reference compounds and harmonized assay conditions. Any signaling-bias claim should use declared reference ligands and an appropriate operational model.

The third priority is to separate receptor state, construct, transducer, and membrane context experimentally. Structures of the same receptor and ligand should be determined in transducer-free, G-protein-bound, and arrestin-bound conditions within one construct lineage. A particularly informative membrane experiment would use the MOR 9PU5 context as the starting point and cross DAMGO with or without BMS-986187 against cholesterol-containing and cholesterol-depleted membranes in the same nanodisc system. This factorial design would test the proposed ligand-lipid interaction without assigning causality from heterogeneous deposited structures.

For structural biology, this work provides a family-wide framework for selecting residues, waters, interfaces, and missing experimental contexts. For computational biology, it demonstrates how exact mapping, complete-pair analysis, and explicit graph-distance strata can turn a heterogeneous structural archive into a reproducible hypothesis engine. For medicinal chemistry, it converts broad concepts such as selectivity, allostery, hydration, and distal coupling into specific ligand modifications, receptor mutations, and controlled comparisons. The atlas does not determine which intervention will succeed. Its power is that it narrows the experimental search space, makes the competing hypotheses explicit, and defines observations capable of proving them wrong.

## Glossary

**Independent experimental map.** One unique experimental data set counted once for structure-level inference; entries sharing the same Electron Microscopy Data Bank map contribute one designated representative.

**Exact-BW position.** A receptor residue with one unambiguous GPCRdb Ballesteros-Weinstein generic label.

**Expected-accession target core.** Selected-chain polymer residues explicitly mapped by SIFTS to the intended UniProt receptor accession.

**Contact graph.** An undirected network whose nodes are mapped receptor residues and whose edges are residue pairs satisfying a stated spatial-contact rule.

**Wiring and rewiring.** Wiring is the edge set within one structure. Rewiring is the static symmetric difference between two edge sets and does not imply a physical transition.

**Primary shared-node estimand.** The pairwise quantity calculated after inducing both receptor graphs on exact-BW positions resolved in both structures.

**Selected-ligand site.** The union of exact-BW receptor residues within 4.5 Å of atoms from curated orthosteric or allosteric ligands.

**Spatial strata.** Direct means graph distance 0 from a resolved site; adjacent means distance 1; connected-distal means finite distance greater than 1; disconnected means no finite path.

**Endpoint mean.** The mean of all eligible pairwise outcomes incident on one structure within the stated analysis scope.

**Deposited water.** A water oxygen represented in a deposited coordinate model; it is not a measurement of bulk or dynamic hydration.

**Structural water and ligand-contact water.** Structural water contacts at least two protein residues under the stated polar-distance rule. Ligand-contact water contacts a selected ligand and at least one exact-BW receptor residue.

**Recurrence.** Reappearance of a coordinate cluster or exact-BW bridge topology in at least two independent maps under the stated rule.

**Exposure and reference.** Exposure denotes presence of a prespecified structural feature; reference denotes its absence within the eligible analysis scope.

**Normalization and standardization.** Normalization divides a quantity by a stated structural, contact, or reference denominator to create a ratio or relative scale. Standardization subtracts a stated mean and divides by a stated standard deviation. Raw counts remain unnormalized unless explicitly identified otherwise.

**Percentage scaling.** A percentage-scaled contrast divides an observed difference or fitted coefficient by the absolute observed reference-group mean and multiplies by 100. An unadjusted percentage contrast uses the observed exposed-minus-reference mean difference; it is not raw-scale data.

**Literature-directed positive control.** A prespecified result expected from structures already represented in the literature, used to evaluate workflow directionality rather than to claim independent replication.

**Adjusted q value.** A *p* value adjusted for the stated Benjamini-Hochberg false-discovery-rate family.

**Interpretation boundary.** The strongest claim justified by the estimand, cohort coverage, multiplicity control, and observational design.

## Methods

### Cohort discovery, curation, and map independence

All numerical analyses were performed with our in-house StrucMind platform.^24^ For every method introduced in this study, the SI supplies human-readable pseudocode and machine-readable parameters, equations, schemas, provenance links, and validation records. The Research Collaboratory for Structural Bioinformatics Protein Data Bank (RCSB PDB) was queried separately for exact reference-sequence-accession matches to P35372 for MOR, P41145 for KOR, P41143 for DOR, and P41146 for NOP. The exact discovery queries, response identifiers, query date, and absence of method, resolution, and release-date filters are supplied in the machine-readable discovery record. All entry hits were retained at discovery, producing 99 unique candidates. RCSB PDB coordinate models and metadata supplied experimental and entity annotations,^25^ GPCRdb supplied receptor annotations,^26^ the Protein Data Bank in Europe Structure Integration with Function, Taxonomy and Sequences resource (PDBe SIFTS) supplied residue-level UniProt correspondence,^27^ and exact generic residue labels followed the Ballesteros-Weinstein scheme.^28^ For accession *a* and candidate chain *c*, the inclusive mapped union span was

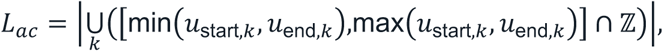

where *k* indexes SIFTS intervals and ℤ denotes integer UniProt positions. Candidate receptor-chain records were ranked by decreasing *L_ac_*, decreasing interval-length-weighted sequence identity, agreement with the GPCRdb preferred chain, agreement with curated-chain evidence, and then lexical author-chain and label-chain identifiers. Inclusion required source taxonomy identifier 9606, *L*_target,*c*_ ≥ 250, and *L*_target,*c*_ at least as large as the mapped union of every alternative accession on *c* . Mapping coverage and identity were retained as quality-control variables rather than additional hard gates.

The selected receptor copy in the asymmetric unit, the deposited crystallographic or reconstruction coordinate unit, supplied the primary receptor graph. Biological assembly 1, the deposited assembly used for environmental context, supplied component, partner, lipid, ion, and water coordinates. Ligand role was separated from component identity: named short opioid peptides were orthosteric only when polymer identity and receptor contact supported that role, and allosteric components were excluded from the orthosteric field. Shared experimental maps remained in the registry but contributed one explicitly designated representative to structure-level models. The tied pairs 10TL/9PU5 and 10TM/9PUD share EMD-71869 and EMD-71871, respectively; 9PU5 and 9PUD were retained because they are the complete receptor-*G_i_* entries. The receptor graphs were identical within each pair. The full cohort therefore contains 86 structures, and the primary panel contains 84 independent-map representatives.

State assignment used GPCRdb annotation when available. GPCRdb Active entries with a populated signaling-protein field were classified as active-partner-bound, Active entries without that field as active-like, and Inactive entries as inactive-like. If GPCRdb state was unavailable, a fixed, case-insensitive title rule classified titles containing inactive, antagonist, or inverse agonist as inactive-like; otherwise titles containing a G-protein or arrestin partner token as active-partner-bound; otherwise titles containing active or agonist as active-like; and all remaining entries as unclear. GPCRdb annotation took precedence in conflicts. The independent-map panel contains 49 active-partner-bound, 16 inactive-like, 4 active-like, and 15 unclear maps. The state models compared only the 49 active-partner-bound and 16 inactive-like maps.

## Exact-BW receptor graphs and single-structure analysis

Coordinate residues were assigned to the expected receptor accession only by exact selected-chain SIFTS correspondence and then to canonical GPCRdb/BW identifiers. No alternative coordinate-chain mapping or heuristic segment-fraction label entered the primary graphs. A graph node required an exact TM1 through TM7 BW label and the atom set required by its contact rule. This produced 19,957 primary Cα graph-node records across 86 structures, a strict subset of the 24,360-residue expected-accession target core. Exact BW labels were one-to-one within every selected chain; any duplicate would have failed graph construction rather than being chosen by an additional rule. Coordinates were taken from the first deposited model. DOR 4N6H was the only selected-receptor coordinate file with alternate locations, meaning multiple deposited coordinates for the same atom; its A and B conformers had equal occupancy, and conformer A was selected. For structure *s* and contact rule *m*, the undirected graph was 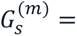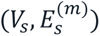. For the primary Cα rule,

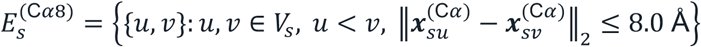

The five sensitivity graphs used Cα cutoffs of 6 and 10 Å, Cβ at 8 Å with Cα used for glycine, minimum inter-residue heavy-atom distance at 4.5 Å, and minimum side-chain distance at 5.0 Å. No sequence-separation exclusion was applied. Single-structure outputs included graph size, edge identity, connectedness, fingerprints, current-flow summaries, curvature, elastic-network summaries, perturbation-response summaries, and community assignments. The primary biological claims use the Cα 8 Å exact-BW graph; alternate rules and the whole receptor-core domain are sensitivity analyses.

## All-to-all structural alignment and contact comparison

Every unordered structure pair was enumerated once. For structures *a* and *b*, *M_ab_* denotes their shared exact-BW positions. Mapped Cα coordinates were superposed by proper-rotation Kabsch alignment.^29^ The pairwise RMSD was

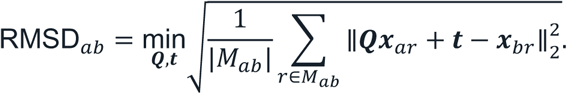

Here, ***x****_ar_* and ***x****_br_* are the mapped Cα coordinate vectors, ***Q*** is constrained to a proper rotation with det(***Q***) = 1, and ***t*** is a translation vector.

For structure *s*, *E_s_* contained every primary contact whose two endpoints had exact-BW assignments in that structure. The primary pair estimand first induced both graphs on their shared exact-BW node set *M_ab_*:

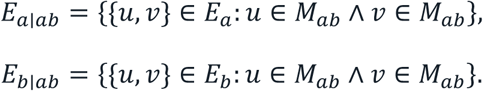

The conserved and union sets were

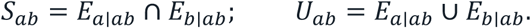

Contact Jaccard similarity and rewired-contact burden were

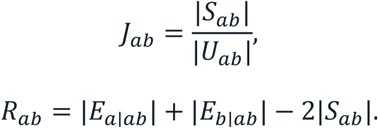

The shared-contact distance RMS deviation used the same conserved edges:

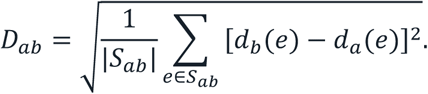

For a conserved edge *e*, *d_a_*(*e*) and *d_b_*(*e*) are the contact-rule-specific distances between its residue endpoints in structures *a* and *b*. No graph had an empty union; *D_ab_* was missing if a pair had no conserved edge. Within the universe *U_ab_*, gained edges were 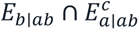, and lost edges were 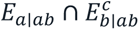 . A full-edge-set sensitivity comparison allowed mapping availability to contribute to the symmetric difference. Let *N*_full_ and *N*_shared_ be the pooled full-edge and shared-node changed-contact totals within one declared scope. Mapping-availability retention and removal were

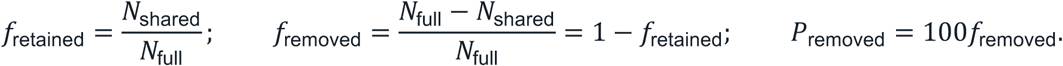

They were undefined when *N*_full_ = 0. The lexical pair order defines *a* and *b* only for reproducible bookkeeping. It has no temporal or mechanistic interpretation. For a pairwise outcome *Y_i_*_j_, let *P_i_* be the set of eligible unordered pairs incident on structure *i*. The structure-level endpoint mean was

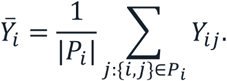

In the complete 84-map panel, each endpoint mean averages 83 pairs. Under a scope restriction or leave-one-out deletion, endpoint means are calculated from the eligible pairs. For the endpoint-outcome redundancy analysis, *x_i_* was endpoint-mean contact Jaccard similarity and *y_i_* was endpoint-mean rewired-contact count for independent map *i*. Pearson correlation across the *N* = 84 complete paired endpoint means was

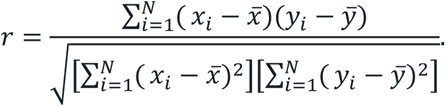

This correlation quantifies outcome redundancy and is not independent biological evidence. An ancillary semantic comparison encoded contacts either in GPCR-specific family bins or in generic universal bins. Burden mode weighted bins by their contact counts, whereas binary mode weighted occupied bins equally. The four resulting settings, family burden, family binary, universal burden, and universal binary, were retained separately for all 3,655 full-cohort pairs, generating 14,620 records. They were not used as evidence for the primary biological claims.

## Ligand, component, and protein-interface annotation

Ligand role, assembly presence, receptor contact, exact-BW interface, and curated biological role were maintained as separate variables. The primary selected-ligand site was coordinate-defined: an exact-BW receptor residue was direct when any nonhydrogen atom of that receptor residue lay within 4.5 Å of any nonhydrogen atom of a role-selected orthosteric or allosteric ligand. PLIP interactions were used as secondary interaction evidence and as an independent target-resolution comparator, not as the primary site definition.^30^ Component and polymer interfaces used the analogous minimum heavy-atom distance rule at 4.0, 4.5, and 5.0 Å; 4.5 Å defined primary receptor-contact exposures. Chain-pair residue contacts were resolved back to receptor residue labels before exact-BW mapping. Gβ contact was tested only within Gα-positive structures because Gα, Gβ, and Gγ assembly presence is aliased with resolved G-protein assembly.

## Site distance and spatial strata

Let *L_s_* be a resolved exact-BW site in structure *s*. The graph distance from node *v* to the site and the corresponding edge distance were

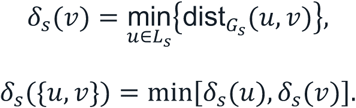

Direct edges have *δ* = 0, adjacent edges have *δ* = 1, connected-distal edges have finite *δ* > 1, and disconnected edges have no finite path. Lost edges use the source-structure graph and site when resolved, otherwise the destination graph and site. Gained edges use the destination graph and site when resolved, otherwise the source graph and site. Conserved edges use the most proximal supported classification across resolved endpoint sites. Pairs with no resolved endpoint site are omitted. The primary selected-ligand pair analysis further requires both endpoint sites to be resolved. Pooled static and changed-contact fractions for class *k* were

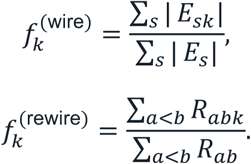

*E_sk_* is the subset of structure *s* edges assigned to class *k*, and *R_abk_* is the number of changed contacts assigned to *k* for pair {*a*, *b*}. Their reported percentage forms were 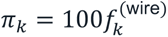 and 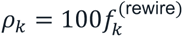. A fraction and its percentage were undefined when their corresponding pooled denominator was zero. Thus, reported localization fractions are pooled edge or changed-contact fractions, not unweighted averages of per-map percentages.

### Deposited-water definitions and recurrence

Let *w* be a deposited water oxygen, *o*(*w*) its occupancy, *d_R_*(*w*) its minimum distance to any designated-receptor heavy atom, *A*(*w*) the set of protein residues with N or O atoms within 3.5 Å, *R*(*w*) the subset of designated-receptor residues in *A*(*w*), *B*(*w*) the subset of exact-BW receptor residues in *A*(*w*), and *L*_sel_(*w*) an indicator of a selected-ligand N or O atom within 3.5 Å. At the default settings,

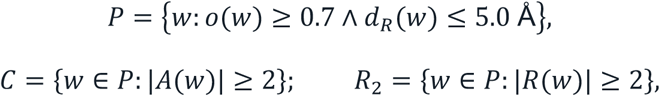

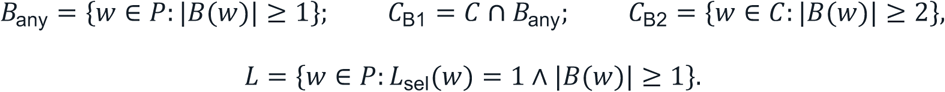

*C*, *R*_2_, and *B*_any_ are subsets of *P*; *R*_2_ is also a subset of *C* because *R*(*w*) ⊆ *A*(*w*); *C*_B2_ ⊆ *C*_B1_ ⊆ *C*; and *L* ⊆ *B*_any_. The designated-receptor two-partner set *R*_2_ is distinct from the exact-BW two-partner set *C*_B2_ . The ligand-contact set *L* is not required to be a subset of *C* . Occupancy sensitivities were 0, 0.3, 0.5, 0.7, 0.9, and 1.0; polar cutoffs were 3.2, 3.5, 3.8, and 4.1 Å. Primary coordinate clustering, one-to-one coordinate matching, and topology recurrence used the 38 default *C*_B2_ waters. The full-panel alignment reference maximized available exact-BW receptor Cα coordinates, with lexicographic structure identifier as the tie breaker; tied DOR maps 4N6H and 4RWD each supplied 240 positions, so 4N6H was selected. Coordinate recurrence was evaluated at 0.8, 1.0, 1.2, 1.5, and 2.0 Å using complete-linkage clusters and connected components of the corresponding single-linkage graph. A cluster was recurrent when it contained waters from at least two independent maps. Pairwise coordinate recurrence used maximum-cardinality one-to-one matching at the same radius. Each *C*_B2_ water contributed every unordered pair among its exact-BW partners to the topology analysis. PLIP event overlap was evaluated separately. The leave-4N6H-out analysis filtered its waters after alignment and retained the 4N6H coordinate frame.

### Normalization, scaling, and denominators

Normalization was prespecified by outcome rather than applied as one global transformation. Node counts, edge counts, and rewired-contact counts remained raw counts. Contact Jaccard similarity normalized conserved contacts by the pairwise union, and its complementary distance equals the symmetric-difference count divided by that union. Shared-contact distance RMS deviation is a root mean square over the conserved-edge set. Site-relative wiring and rewiring fractions divided pooled class-specific counts by the corresponding pooled edge or changed-contact total; they are contact-weighted fractions, not equally weighted map averages. Endpoint means averaged all eligible pairwise outcomes incident on one map so each map contributed one model record within a stated scope. Unadjusted and adjusted percentage contrasts divided an observed difference or fitted exposure coefficient by the absolute observed reference-group mean. Continuous model and matching covariates were standardized within the prespecified analysis scope using the population standard deviation. After centering, a zero population standard deviation was replaced by 1 so a constant covariate had standardized value 0; this numerical safeguard did not impute a missing covariate. Descriptive graph fingerprints were standardized across all 86 structures before principal component analysis; current-flow and perturbation-response diagnostics used their separately stated graph-specific normalizations. Unresolved sites and missing outcomes were excluded from their relevant denominator and were never converted to zero. The SI normalization registry states the numerator, denominator or centering rule, scope, zero and missing-value policy, and interpretation for every reported family of quantities.

### Prevalence, percentage contrasts, and uncertainty

For *k* positive maps among *n*, prevalence was *p* = *k*/*n* and its 95% Wilson binomial-proportion confidence interval used *z* = 1.959963984540054.^31^ Wilson endpoints were calculated on the unit-proportion scale and reported as percentages using CI_W_(%) = 100 CI_W_. For exposed and reference structure means, the unadjusted percentage contrast was

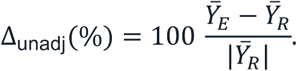

The unadjusted spatial-composition intervals in SI Table S5 used a stratified fixed-map-record bootstrap.^32^ For a pairwise outcome, one endpoint mean was first computed for each of the 77 selected-site-positive maps over its 76 observed pairs. For an intrinsic static outcome, the observed per-map fraction was carried directly. Each of 5,000 replicates sampled exposed and reference map records separately with replacement at their observed group sizes while carrying the relevant fixed map value unchanged. Pair rows were not resampled, the pair graph was not rebuilt, and endpoint means were not recomputed. For estimable contrasts in the separate percentage-effect sensitivity tables, 10,000 independent structure-bootstrap resamples were drawn. Publication clustering attempted 10,000 publication-key resamples with replacement and retained every map belonging to each selected key occurrence; percentile limits used only replicates containing both exposure levels and an absolute reference mean greater than 10^−15^. Publication groups were identified successively by PubMed identifier, DOI, normalized article title, and PDB accession when no publication identifier was available. For matched pair *l*, the percentage contrast and its valid-pair mean were

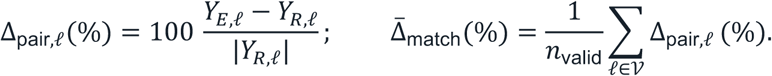

A pair belonged to V only when both outcomes were finite and |*Y_R_*_,*P*_| > 10^−15^; *n*_valid_ = |V|, and the mean was undefined when *n*_valid_ = 0 . Matched-pair percentage intervals used 10,000 bootstrap resamples of the valid pair-level percentages. Deterministic Secure Hash Algorithm 256-bit (SHA-256)-derived seeds included the contrast, outcome, and resampling-mode identifiers. A group-level percentage was undefined if the absolute reference mean was at most 10^−15^.

### State positive-control and component-association models

The state family used inactive-like and active-partner-bound maps only. Its adjustment block was

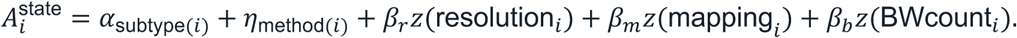

For outcome *Y_i_*, the fitted model was

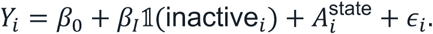

The state-adjusted percentage was

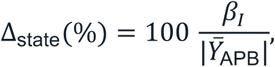

where *Ȳ*_APB_ is the observed active-partner-bound mean in the fitted scope; coefficient confidence limits were scaled by the same denominator. Mapping is the maximum SIFTS-reported coverage among the intervals for the selected expected-accession chain record, and BWcount is the number of nodes in that map’s primary exact-BW Cα graph. The 16 estimable state outcomes formed one literature-directed Benjamini-Hochberg family.

For each component contrast, the nonexposure adjustment block was

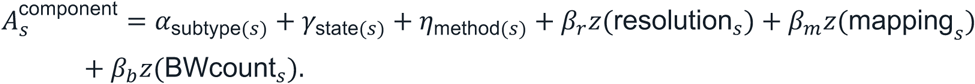

The generic primary model was

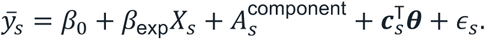

Here, *ȳ_s_* is the endpoint mean for a pairwise outcome or the observed value for an intrinsic outcome; *X_s_* is the binary exposure; *β*_exp_ is its exposed-minus-reference coefficient; and ***c****_s_* is the contrast-specific context vector. Collecting all nonexposure adjustment terms gives **Γ**^T^***Z****_s_*, where ***Z****_s_* is the covariate vector and **Γ** is its coefficient vector.

For a continuous covariate *x* in an *H*-map analysis scope, standardization used

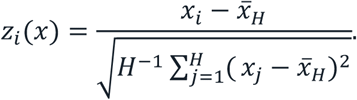

If the population standard deviation in this expression was zero, the scale was set to 1 after centering, so every observed value of that constant covariate mapped to 0; missing covariates were not imputed.

The additional context vector ***c****_s_* was contrast-specific: curated binding mode and resolved G-protein context for sterol, generic lipid, deposited water, and structural water; binding mode alone for G-protein assembly and within-Gα Gβ contact; resolved G-protein and receptor-contacting allosteric-modulator indicators for peptide versus small molecule; and resolved G-protein context plus orthosteric modality for orthosteric-plus-allosteric versus orthosteric-only binding. Within each scope, categorical levels represented by fewer than five maps were combined as an other-sparse category. A missing categorical value was encoded as unknown before sparse-level collapsing, whereas a map missing the outcome, exposure, resolution, mapping coverage, or BWcount was excluded from that model. Eight defined component contrasts were crossed with four outcomes: exact-BW contact-pair count, endpoint-mean Jaccard similarity, endpoint-mean rewired-contact count, and endpoint-mean shared-contact distance RMS deviation. All 32 models were estimable and full rank, meaning that fitted design columns had no exact linear dependence.

Ordinary least squares inference used HC3 covariance with a t reference distribution.^33^ The adjusted percentage was

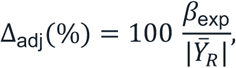

where *Ŷ_R_* is the observed reference-group mean in the fitted scope; coefficient confidence limits were scaled by the same denominator. Publication-cluster covariance was a sensitivity analysis when at least 10 publication groups were available. Matching was exact on experimental method, receptor subtype, and state. Resolution, mapping coverage, and exact-BW mapped-residue count were standardized once across the complete hypothesis-specific analysis scope using population standard deviations. Within each exact stratum, maps were paired without replacement and without a distance caliper by minimum-cost linear assignment on the resulting Euclidean distance.^34^ Structure and publication leave-one-out analyses deleted the unit and all pairwise records involving that unit, recomputed endpoint means, and refit the model.

The Benjamini-Hochberg procedure^35^ was applied to ordered *p* values *p*_(1)_ ≤ ⋯ ≤ *p*_(*m*)_ as follows.

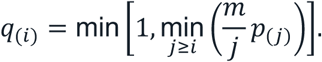

Here, *m* = 16 for the state family and *m* = 32 for the component family. PIP₂-like lipid, β-arrestin, sodium, and selected-ligand/water/receptor bridging were too sparse for the defined component-model family and were restricted to prevalence and structure-specific description.

### Reproducible scientific specification

For every method introduced in this study, the Supporting Information supplies human-readable pseudocode and an equivalent machine-readable scientific specification containing ordered operations, complete parameter values, equations and symbol definitions, authoritative table schemas, a public data dictionary, derivation and validation records, and a claim-to-source map. Separate non-image source tables contain the numerical and categorical data underlying every figure; no figure-rendering code or figure-specific pseudocode is included. Stable artifact identifiers, record keys, field names, rounding rules, and Secure Hash Algorithm 256-bit (SHA-256) cryptographic checksums connect reported values to their scientific inputs and transformations. This design follows JCIM and joint ACS guidance on method, data, software, and workflow transparency.^36^

## Associated Content

### Supporting Information

Expanded cohort curation; exact-BW mapping; graph, pair, component, interface, water, and statistical definitions; complete state and component outcome tables; water recurrence; claim classification; pseudocode restricted to methods introduced in this study; machine-readable parameters, equations, schemas, derivations, validation records, and checksums; and non-image source data underlying every figure (DOCX, CSV, JSON, Markdown, and text).

### Data and Software Availability

All manuscript-derived tables, data underlying each figure, pseudocode of methods introduced in this study, method parameters, equation definitions, public schemas, derivation records, validation records, and checksum manifests are supplied with the Supporting Information. Source coordinate models remain available from the RCSB Protein Data Bank under their accession identifiers.

## Notes

The authors declare no competing financial interest.

## Acknowledgments

This publication was made possible by an Institutional Development Award (IDeA) from the National Institute of General Medical Sciences of the National Institutes of Health under Grant # 2P20GM103432. The content is solely the responsibility of the authors and does not necessarily represent the official views of the National Institutes of Health.

## References

(1) Manglik, A.; Kruse, A. C.; Kobilka, T. S.; Thian, F. S.; Mathiesen, J. M.; Sunahara, R. K.; Pardo, L.; Weis, W. I.; Kobilka, B. K.; Granier, S. Crystal structure of the µ-opioid receptor bound to a morphinan antagonist. Nature 2012, 485 (7398), 321–326. DOI: 10.1038/nature10954.

(2) Wu, H.; Wacker, D.; Mileni, M.; Katritch, V.; Han, G. W.; Vardy, E.; Liu, W.; Thompson, A. A.; Huang, X.-P.; Carroll, F. I.;, et al. Structure of the human κ-opioid receptor in complex with JDTic. Nature 2012, 485 (7398), 327–332. DOI: 10.1038/nature10939.

(3) Granier, S.; Manglik, A.; Kruse, A. C.; Kobilka, T. S.; Thian, F. S.; Weis, W. I.; Kobilka, B. K. Structure of the δ-opioid receptor bound to naltrindole. Nature 2012, 485 (7398), 400–404. DOI: 10.1038/nature11111.

(4) Thompson, A. A.; Liu, W.; Chun, E.; Katritch, V.; Wu, H.; Vardy, E.; Huang, X.-P.; Trapella, C.; Guerrini, R.; Calo, G.;, et al. Structure of the nociceptin/orphanin FQ receptor in complex with a peptide mimetic. Nature 2012, 485 (7398), 395–399. DOI: 10.1038/nature11085.

(5) Huang, W.; Manglik, A.; Venkatakrishnan, A. J.; Laeremans, T.; Feinberg, E. N.; Sanborn, A. L.; Kato, H. E.; Livingston, K. E.; Thorsen, T. S.; Kling, R. C.;, et al. Structural insights into µ-opioid receptor activation. Nature 2015, 524 (7565), 315–321. DOI: 10.1038/nature14886.

(6) Sounier, R.; Mas, C.; Steyaert, J.; Laeremans, T.; Manglik, A.; Huang, W.; Kobilka, B. K.; Déméné, H.; Granier, S. Propagation of conformational changes during μ-opioid receptor activation. Nature 2015, 524 (7565), 375–378. DOI: 10.1038/nature14680.

(7) Koehl, A.; Hu, H.; Maeda, S.; Zhang, Y.; Qu, Q.; Paggi, J. M.; Latorraca, N. R.; Hilger, D.; Dawson, R.; Matile, H.;, et al. Structure of the µ-opioid receptor–Gi protein complex. Nature 2018, 558 (7711), 547–552. DOI: 10.1038/s41586-018-0219-7.

(8) Claff, T.; Yu, J.; Blais, V.; Patel, N.; Martin, C.; Wu, L.; Han, G. W.; Holleran, B. J.; Van der Poorten, O.; White, K. L.;, et al. Elucidating the active δ-opioid receptor crystal structure with peptide and small-molecule agonists. Science Advances 2019, 5 (11), eaax9115. DOI: doi:10.1126/sciadv.aax9115.

(9) Wang, Y.; Zhuang, Y.; DiBerto, J. F.; Zhou, X. E.; Schmitz, G. P.; Yuan, Q.; Jain, M. K.; Liu, W.; Melcher, K.; Jiang, Y.;, et al. Structures of the entire human opioid receptor family. Cell 2023, 186 (2), 413–427.e417. DOI: 10.1016/j.cell.2022.12.026 (accessed 2026/08/07).

(10) Han, J.; Zhang, J.; Nazarova, A. L.; Bernhard, S. M.; Krumm, B. E.; Zhao, L.; Lam, J. H.; Rangari, V. A.; Majumdar, S.; Nichols, D. E.;, et al. Ligand and G-protein selectivity in the κ-opioid receptor. Nature 2023, 617 (7960), 417–425. DOI: 10.1038/s41586-023-06030-7.

(11) Zhao, J.; Elgeti, M.; O’Brien, E. S.; Sár, C. P.; Ei Daibani, A.; Heng, J.; Sun, X.; White, E.; Che, T.; Hubbell, W. L.;, et al. Ligand efficacy modulates conformational dynamics of the µ-opioid receptor. Nature 2024, 629 (8011), 474–480. DOI: 10.1038/s41586-024-07295-2.

(12) Han, J.; Fine, E. J.; Jiang, Q.; Zhuang, Y.; Suomivuori, C.-M.; Chen, Z.-W.; Denn, E.; Whiddon, K.; Li, K.; Evers, A. S.;, et al. Structural dynamics of kappa opioid receptor interactions with β-arrestin 1. Nature Communications 2026, 17 (1), 7171. DOI: 10.1038/s41467-026-73968-3.

(13) Zhang, H.; Wang, X.; Xi, K.; Shen, Q.; Xue, J.; Zhu, Y.; Zang, S.-K.; Yu, T.; Shen, D.-D.; Guo, J.;, et al. The molecular basis of μ-opioid receptor signaling plasticity. Cell Research 2025, 35 (12), 1021–1036. DOI: 10.1038/s41422-025-01191-8.

(14) Kaneko, S.; Imai, S.; Uchikubo-Kamo, T.; Hisano, T.; Asao, N.; Shirouzu, M.; Shimada, I. Structural and dynamic insights into the activation of the μ-opioid receptor by an allosteric modulator. Nature Communications 2024, 15 (1), 3544. DOI: 10.1038/s41467-024-47792-6.

(15) Zhang, H.; Konovalov, K.; Parpounas, A. K.; Provasi, D.; Yang, S.; Abraham, A.; Vela, A. M.; Warren, A. L.; Zilberg, G.; Wang, S.;, et al. Structural and dynamic studies uncover a distinct allosteric modulatory site at the µ-opioid receptor. Nature Communications 2026, 17 (1), 6000. DOI: 10.1038/s41467-026-72633-z.

(16) Wang, Y.; Luo, P.; Xu, H.; Zhan, L.; Sakamoto, K.; Li, M.; Wang, J.; Huang, X.-P.; Zhou, J.; Liu, T.;, et al. Structure-based design of an opioid receptor modulator for enhanced morphine analgesia. Science Advances 2026, 12 (7), eaea9832. DOI: doi:10.1126/sciadv.aea9832.

(17) Venkatakrishnan, A. J.; Ma, A. K.; Fonseca, R.; Latorraca, N. R.; Kelly, B.; Betz, R. M.; Asawa, C.; Kobilka, B. K.; Dror, R. O. Diverse GPCRs exhibit conserved water networks for stabilization and activation. Proceedings of the National Academy of Sciences 2019, 116 (8), 3288–3293. DOI: doi:10.1073/pnas.1809251116.

(18) Fenalti, G.; Giguere, P. M.; Katritch, V.; Huang, X.-P.; Thompson, A. A.; Cherezov, V.; Roth, B. L.; Stevens, R. C. Molecular control of δ-opioid receptor signalling. Nature 2014, 506 (7487), 191–196. DOI: 10.1038/nature12944.

(19) Ople, R. S.; Ramos-Gonzalez, N.; Li, Q.; Sobecks, B. L.; Aydin, D.; Powers, A. S.; Faouzi, A.; Polacco, B. J.; Bernhard, S. M.; Appourchaux, K.;, et al. Signaling Modulation Mediated by Ligand Water Interactions with the Sodium Site at μOR. ACS Central Science 2024, 10 (8), 1490–1503. DOI: 10.1021/acscentsci.4c00525 (accessed 8/7/2026).

(20) Zheng, H.; Zou, H.; Liu, X.; Chu, J.; Zhou, Y.; Loh, H. H.; Law, P.-Y. Cholesterol level influences opioid signaling in cell models and analgesia in mice and humans. Journal of Lipid Research 2012, 53 (6), 1153–1162. DOI: 10.1194/jlr.M024455 (accessed 2026/08/07).

(21) Radoux-Mergault, A.; Oberhauser, L.; Aureli, S.; Gervasio, F. L.; Stoeber, M. Subcellular location defines GPCR signal transduction. Science Advances 2023, 9 (16), eadf6059. DOI: doi:10.1126/sciadv.adf6059.

(22) Janetzko, J.; Kise, R.; Barsi-Rhyne, B.; Siepe, D. H.; Heydenreich, F. M.; Kawakami, K.; Masureel, M.; Maeda, S.; Garcia, K. C.; von Zastrow, M.;, et al. Membrane phosphoinositides regulate GPCR-&#x3b2;-arrestin complex assembly and dynamics. Cell 2022, 185 (24), 4560– 4573.e4519. DOI: 10.1016/j.cell.2022.10.018 (accessed 2026/08/07).

(23) Varga, B. R.; Bernhard, S. M.; El Daibani, A.; Zaidi, S. A.; Lam, J. H.; Aguilar, J.; Appourchaux, K.; Nazarova, A. L.; Kouvelis, A.; Shinouchi, R.;, et al. Structure-guided design of partial agonists at an opioid receptor. Nature Communications 2025, 16 (1), 2518. DOI: 10.1038/s41467-025-57734-5.

(24) Nael, M. A.; Alakonda, L. M.; Elokely, K. M. Contact-Network Phenotyping of the CDK Family Reveals Selective Distal C-Lobe Contact Redistribution by Modern CDK5 Inhibitors and a Quantitative Selectivity Landscape against CDK2 and CDK1. Journal of Chemical Information and Modeling 2026, 66 (12), 7276–7295. DOI: 10.1021/acs.jcim.6c00886 (accessed 8/8/2026).

(25) Vallat, B.; Rose, Y.; Piehl, D. W.; Duarte, J. M.; Bittrich, S.; Bi, C.; Segura, J.; Zalevsky, A.; Sekharan, M. R.; Webb, B. M.;, et al. RCSB Protein Data Bank: Delivering integrative structures alongside experimental structures and computed structure models. Nucleic Acids Research 2026, 54 (D1), D489–D498. DOI: 10.1093/nar/gkaf1187 (accessed 8/8/2026).

(26) Pándy-Szekeres, G.; Caroli, J.; Mamyrbekov, A.; Kermani, A. A.; Keserű, György M.; Kooistra, Albert J.; Gloriam, D. E. GPCRdb in 2023: state-specific structure models using AlphaFold2 and new ligand resources. Nucleic Acids Research 2023, 51 (D1), D395–D402. DOI: 10.1093/nar/gkac1013 (accessed 8/8/2026).

(27) Dana, J. M.; Gutmanas, A.; Tyagi, N.; Qi, G.; O’Donovan, C.; Martin, M.; Velankar, S. SIFTS: updated Structure Integration with Function, Taxonomy and Sequences resource allows 40-fold increase in coverage of structure-based annotations for proteins. Nucleic Acids Research 2019, *47* (D1), D482–D489. DOI: 10.1093/nar/gky1114 (accessed 8/8/2026).

28. Ballesteros, J. A.; Weinstein, H. [19] Integrated methods for the construction of three-dimensional models and computational probing of structure-function relations in G protein-coupled receptors. In Methods in Neurosciences, Sealfon, S. C. Ed.; Vol. 25; Academic Press, 1995; pp 366–428.

(29) Kabsch, W. A solution for the best rotation to relate two sets of vectors. Acta Crystallographica Section A 1976, 32 (5), 922–923. DOI: doi:10.1107/S0567739476001873.

(30) Adasme, M. F.; Linnemann, K. L.; Bolz, S. N.; Kaiser, F.; Salentin, S.; Haupt, V J.; Schroeder, M. PLIP 2021: expanding the scope of the protein–ligand interaction profiler to DNA and RNA. Nucleic Acids Research 2021, 49 (W1), W530–W534. DOI: 10.1093/nar/gkab294 (accessed 8/8/2026).

(31) Wilson, E. B. Probable Inference, the Law of Succession, and Statistical Inference. Journal of the American Statistical Association 1927, 22 (158), 209–212. DOI: 10.1080/01621459.1927.10502953.

(32) Efron, B. Bootstrap Methods: Another Look at the Jackknife. The Annals of Statistics 1979, 7 (1), 1–26, 26.

(33) MacKinnon, J. G.; White, H. Some heteroskedasticity-consistent covariance matrix estimators with improved finite sample properties. Journal of Econometrics 1985, 29 (3), 305–325. DOI: 10.1016/0304-4076(85)90158-7.

(34) Kuhn, H. W. The Hungarian method for the assignment problem. Naval Research Logistics Quarterly 1955, 2 (1-2), 83–97. DOI: 10.1002/nav.3800020109 (accessed 2026/08/07).

(35) Benjamini, Y.; Hochberg, Y. Controlling the False Discovery Rate: A Practical and Powerful Approach to Multiple Testing. Journal of the Royal Statistical Society: Series B (Methodological) 1995, 57 (1), 289–300. DOI: 10.1111/j.2517-6161.1995.tb02031.x (accessed 8/8/2026).

(36) Amaro, R. E.; Batista, V.; Blumberger, J.; Choong, Y. S.; Corminboeuf, C. m.; Cournia, Z.; Cui, Q.; De Vivo, M.; Evangelista, F. A.; Gao, Y. Q.;, et al. Advancing Reproducibility and Open Data in Theoretical and Computational Chemistry. Journal of Chemical Theory and Computation 2026, 22 (9), 4199–4200. DOI: 10.1021/acs.jctc.6c00733 (accessed 8/8/2026).

