## Supplementary material for "An Exact-Residue Atlas of Opioid Receptor Wiring and Rewiring across Ligand and Transducer Contexts": SI files: README.md

# Supporting data and reproducibility files

This directory contains the light, submission-facing data and scientific specifications supporting the Article and Supporting Information.

## Data groups

- `Data_*.csv` files contain compact authoritative summaries and extracted values used in the Article and Supporting Information.
- `Sensitivity_*.csv` files contain robustness, alternate-estimand, matching, leave-one-out, component, spatial, and deposited-water results.
- `Data_component_matched_pair_detail.csv` releases the exposed and unexposed structure identifiers and outcome values for every matched pair, allowing every matched native and percentage difference to be recalculated directly.
- `Figure_*.csv` files contain the numerical or categorical source data used to build each displayed figure panel. 
- The even-numbered `Figure_02`, `Figure_06`, `Figure_08`, `Figure_10`, `Figure_12`, and `Figure_14` evidence CSVs, together with `exact_bw_crosswalk.csv`, contain the residue, component, pair-selection, and deposited-water evidence used for the integrated three-dimensional Article figures. Figure 4 uses the coordinate-provenance table directly.
- `STRUCTURAL_COMPANION_PROVENANCE.csv` identifies the canonical deposited structures and the reduced editable PDB files supplied in the `figures` directory.

## Reproducible scientific specification

- `SUPPLEMENTARY_METHODS_PSEUDOCODE.md` and `SUPPLEMENTARY_METHODS_PSEUDOCODE.json` contain pseudocode only for the six composite methods introduced in this study, P01 through P06.
- `METHOD_PARAMETERS.json`, `METHOD_EQUATIONS.json`, `EQUATION_SYMBOLS.csv`, `ANALYSIS_CONTRASTS.csv`, `STATE_OUTCOMES.csv`, `COMPONENT_TESTS.csv`, and `WATER_ESTIMAND_DEFINITIONS.json` define the complete scientific operations, parameters, equations, contrasts, and estimands.
- `NORMALIZATION_REGISTRY.csv` classifies every reported quantity as raw, normalized, standardized, or percentage-scaled and records its numerator, denominator or reference, analysis scope, zero rule, missing-value rule, and interpretation.
- `PUBLIC_DATA_DICTIONARY.csv`, `PUBLIC_TABLE_SCHEMAS.json`, `DERIVATION_MANIFEST.json`, and `CLAIM_TO_SOURCE_MAP.csv` connect public filenames and fields to derivations and reported claims. Every released CSV has exactly one stable artifact identifier and exactly one schema record.
- `METHODS_ARTIFACTS.schema.json`, `METHODS_ARTIFACT_INDEX.json`, and `VALIDATION_REPORT.json` provide machine-readable validation structure, method-file checksums, and validation outcomes.
- `FIGURE_TO_SOURCE_MAP.csv`, `FILE_INDEX.csv`, and `SHA256SUMS.txt` provide package navigation and byte-level integrity checks. In `FIGURE_TO_SOURCE_MAP.csv`, PDF names are relative to the sibling `figures` directory and source-data names are relative to this directory. Paths in `FILE_INDEX.csv` and `SHA256SUMS.txt` are relative to the package root. To avoid recursive hashes, `FILE_INDEX.csv` omits itself and `SHA256SUMS.txt`, whereas `SHA256SUMS.txt` omits only itself and includes `FILE_INDEX.csv`.

## Normalization and missing values

Normalization was specified by outcome, not applied as one global transformation. Raw graph and rewiring counts remain counts. Contact Jaccard distance is normalized by the pairwise contact union. Site-centered fractions use their stated pooled edge or changed-contact denominators. Percentage effects use the absolute observed reference-group mean and are undefined when its magnitude is at most 1e-15. Matching variables and principal-component inputs use the explicitly stated within-scope standardization. The 3×32/3×35 water topology is reported against both explicit denominators: 3/84 processed maps, 3.57%, and 3/7 C<sub>B2</sub>-positive maps, 42.86%. Blank or unresolved fields represent missing, inapplicable, or non-estimable values according to the relevant schema and are never silently converted to zero.
