## Supplementary material for "An Exact-Residue Atlas of Opioid Receptor Wiring and Rewiring across Ligand and Transducer Contexts": SI files: SUPPLEMENTARY_METHODS_PSEUDOCODE.md

# Pseudocode for introduced composite methods

Only the six composite methods introduced in this work are pseudocoded here. Cohort curation, state and component regression, standard alignment, resampling, and standard graph descriptors remain fully specified in the parameter and equation registries, but they are not presented as newly introduced methods. Every reported scale follows the numerator, denominator or reference, zero, and missing-data rules in `NORMALIZATION_REGISTRY.csv`.

## P01. Exact-BW receptor contact-graph construction

**Inputs:** curated receptor-chain records, exact SIFTS coordinate-to-UniProt mappings, GPCRdb generic-number mappings.

**Parameters:** `mapping`; `coordinate_selection.receptor_graph_construction`; `contact_graphs`.

**Equations:** E001 and E002.

```text
for each curated structure s:
    map selected-chain coordinate residues to the expected target UniProt accession by exact SIFTS correspondence
    map each retained target residue to an exact TM1 through TM7 GPCRdb generic number
    exclude heuristic, alternative-chain, non-target, and duplicate exact-BW mappings
    read the first deposited coordinate model and apply the declared alternate-location selection
    create one graph node for each unique exact-BW residue with the required coordinate
    for each distinct mapped residue pair {u,v}:
        if the CA distance is at most 8.0 A:
            add the canonical undirected primary edge {u,v}
    repeat graph construction under the five declared contact-definition sensitivities
    assert no self-edge, canonical endpoint ordering, and one node per exact-BW position
```

An unmapped residue or a residue lacking the coordinate required by a contact mode is excluded from that graph and is not encoded as a noncontact. The resulting edges are static distance-defined contacts on exactly mapped target-receptor residues.

## P02. Shared-node pair comparison and endpoint aggregation

**Inputs:** exact-BW contact graphs from P01.

**Parameters:** `pairwise_comparison`; `statistics.endpoint_mean`; `statistics.leave_one_out`.

**Equations:** E003 through E007, E022, E035, E036, and E055.

```text
enumerate every unordered curated-structure pair (a,b)
for each pair:
    M_ab = intersection of exact-BW node sets V_a and V_b
    align shared C-alpha coordinates by the standard Kabsch operation
    induce both primary graphs on M_ab
    calculate intersection, union, gained, lost, conserved, and rewired edge sets
    calculate contact Jaccard similarity or distance, aligned RMSD,
        rewired-contact count, and conserved-edge distance change
    repeat edge-set calculations on the uninduced per-structure graphs
        as a mapping-availability sensitivity
for each independent-map endpoint s and requested dyadic outcome y:
    average all retained incident dyads containing s
for each structure or publication deletion:
    remove incident dyads first, then recompute every endpoint mean
pair complete map-level endpoint outcomes and calculate the Pearson redundancy correlation
```

The primary edge universe contains only edges whose endpoints are available in both structures. Endpoint means use only retained incident dyads with an observed requested outcome. A zero denominator remains missing. Pair differences and endpoint means are descriptive static comparisons and do not remove shared-data dependence.

## P03. Role-aware interface, site, and spatial classification

**Inputs:** curated roles, deposited coordinates, P01 graphs, and P02 changed-edge sets.

**Parameters:** `component_interfaces`; `coordinate_selection.component_water_coordinate_selection.general_repeated_atom_ranking`; `site_localization`.

**Equations:** E008 through E012.

```text
for each curated ligand, component, or protein partner:
    retain biological role, assembly presence, and direct receptor contact as separate labels
    select repeated atom records by the declared alternate-location and occupancy ranking
    resolve receptor heavy-atom contacts and map contacted receptor residues to exact BW
for each resolved role-aware site and receptor graph:
    calculate shortest-path distance from every node or edge to the site
    classify direct, adjacent, connected distal, and disconnected strata
for each changed-edge pair comparison:
    assign lost edges to the source site with destination fallback
    assign gained edges to the destination site with source fallback
    assign conserved edges to the most proximal resolved endpoint site
    omit pairs for which neither endpoint site is resolved
pool static and changed contacts only after structure-level and pair-level classification
```

Deposited absence is not biological absence. Unknown site status is missing and is never converted to zero. Site-relative distance and composition do not establish signaling direction, physical propagation, or causation.

## P04. Deposited-water estimands and recurrence

**Inputs:** deposited coordinates and curated selected-ligand roles.

**Parameters:** component and water coordinate selection; `water_analysis.default`; occupancy and polar-cutoff sensitivities; `water_analysis.recurrence`; `water_analysis.water_graph`.

**Equations:** E003 and E013 through E021, plus E038.

```text
for each deposited water residue:
    select one oxygen coordinate by maximum occupancy and deposition-order tie break
    classify membership in the receptor-proximal root set
    independently classify protein contact, receptor contact, exact-BW bridge,
        and selected-ligand-contact branches
retain one coordinate-selection record per default root-set water
repeat the occupancy and polar-cutoff sensitivity grid
for coordinate recurrence:
    select the reference receptor frame by the deterministic exact-BW coverage rule
    align receptor frames
    cluster coordinates separately by complete linkage and single linkage
    perform pairwise one-to-one coordinate matching without replacement
    remove 4N6H, reselect 4RWD, realign, and repeat the reported conditions
for topology recurrence:
    count reference-free exact-BW water-mediated residue pairs across independent maps
deduplicate role-aware interaction events before cross-validation summaries
```

Unprocessed, unresolved, or out-of-scope events remain missing, and absent deposited waters are not imputed. The ligand-contact set is a separate branch rather than a sixth core stage. These estimands describe static deposited waters and recurrence, not solution hydration, residence time, or causal mediation.

## P05. StrucFlow workflow diagnostic

**Inputs:** whole-core contact ontologies and their geometry, energetic, water, dynamic, and recurrence annotations.

**Parameters:** `ancillary_graph_descriptors_and_sensitivities.strucflow_workflow_diagnostic`.

**Equations:** E047 through E050.

```text
for each independent-map structure:
    construct geometry, signed energetic-frustration, water-mediation,
        absolute dynamic-correlation, and recurrence channels
    apply the declared precedence and fallback rules
    combine channels with fixed weights into a sparse composite contact tensor
    calculate the dimensionless graph score relative to the initial tensor
    run seed-7 event mode for 100 proposals with bounded candidates,
        declared acceptance branches, incremental updates, and periodic exact recalculation
    run seed-7 continuous-weight mode for 50 iterations with batches of eight,
        bounded central differences, Gaussian noise, clipping, and periodic exact recalculation
    retain one diagnostic record for each structure and mode
```

Missing physical annotations contribute only their declared channel fallback and never become positive evidence. StrucFlow outputs are contact-topology workflow diagnostics. Event counts are not physical time, continuous weights are not molecular dynamics, and graph scores are not calibrated free energies.

## P06. Contact-propagation network diagnostic

**Inputs:** the unthresholded composite contact ontology and role-matched, target-matched selected-ligand exact-BW contacts.

**Parameters:** `ancillary_graph_descriptors_and_sensitivities.contact_propagation_network_diagnostic`.

**Equations:** E051 through E054.

```text
for each structure:
    certify exact-BW seed residues from role-matched and target-matched selected-ligand contacts
    if no seed can be certified:
        retain the structure as not applicable and do not impute a seed
    otherwise:
        construct baseline and seeded three-state factor graphs from identical contacts
        calculate node aggregates and floored unary potentials
        retain at most 20000 strongest pair factors
        floor each 3 by 3 pair potential and divide it by its matrix mean
        run uniformly initialized damped loopy sum-product separately for both graphs
        retain convergence status and per-residue state-belief differences
        report the maximum absolute seeded-minus-baseline belief shift
verify that the explicit no-seed control has zero maximum belief shift
```

A missing seed remains not applicable. Failed convergence remains explicit rather than becoming a zero shift. CPN output is a belief-space workflow response, not a physical perturbation trajectory, signaling efficacy, mechanistic propagation, or causal effect.
