## Supplementary material for "An Exact-Residue Atlas of Opioid Receptor Wiring and Rewiring across Ligand and Transducer Contexts": SI document

##### Contents

1. Supplementary Methods S1 to S9
2. Supplementary Results and interpretation boundaries S10 to S14
3. Supplementary Tables S1 to S8
4. Supplementary Figures S1 to S8
5. Machine-readable methods and data index

#### Supplementary Methods

All calculations described below were performed with our in-house StrucMind platform. For each method introduced in this study, the accompanying Supporting Information (SI) reproducibility files provide human-readable pseudocode and machine-readable operations, parameters, equations, schemas, provenance links, and validation criteria.

##### S1. Candidate discovery, curation, and independent-map panel

The Research Collaboratory for Structural Bioinformatics Protein Data Bank (RCSB PDB) discovery snapshot was queried. An independent experimental map is one unique experimental data set counted once for structure-level inference; entries sharing an Electron Microscopy Data Bank (EMDB) map were grouped before representative selection. Four exact-match searches targeted the rcsb_polymer_entity_container_identifiers.reference_sequence_identifiers.database_accession field for the mu-opioid receptor (MOR), P35372; kappa-opioid receptor (KOR), P41145; delta-opioid receptor (DOR), P41143; and nociceptin/orphanin FQ receptor (NOP), P41146. Every entry hit was retained at discovery, without an experimental-method, resolution, or release-date filter, and the four hit lists were combined by PDB identifier. The exact query specification, response identifiers, and query date are retained in the machine-readable discovery record. RCSB PDB coordinate models and metadata supplied experimental entries, entity annotations, source organism, experimental method, and resolution. The G protein-coupled receptor database (GPCRdb) supplied receptor annotations, and the Protein Data Bank in Europe Structure Integration with Function, Taxonomy and Sequences resource (PDBe SIFTS) supplied residue-level UniProt mappings. An exact Ballesteros-Weinstein (BW) label is an unambiguous generic transmembrane residue position. Sources are cited in the Article.

For accession $a$ and coordinate chain $c$, the inclusive mapped union span was

$$L_{ac} = \left|\bigcup_k \left(\left[\min\left(u_{\text{start},k}, u_{\text{end},k}\right), \max\left(u_{\text{start},k}, u_{\text{end},k}\right)\right] \cap \mathbb{Z}\right)\right|,$$

where $k$ indexes SIFTS UniProt intervals and $\mathbb{Z}$ denotes integer UniProt positions. When more than one candidate record was available, records were ranked by decreasing $L_{ac}$; decreasing interval-length-weighted sequence identity; agreement with the GPCRdb preferred chain; agreement with prior curated-chain evidence; and lexical author-chain and label-chain identifiers. For 23SO, SIFTS verification selected chain R. Inclusion required human taxonomy identifier 9606, $L_{\text{target},c} \geq 250$, and $L_{\text{target},c}$ at least as large as the mapped union of every alternative accession on $c$. Mapping coverage and identity were recorded as quality-control variables and had no additional hard threshold.

For interval length $\ell_k = |u_{\text{end},k} - u_{\text{start},k}| + 1$ and SIFTS-reported interval identity $i_k$, the ranking identity was

$$I_{ac} = \frac{\sum_k \ell_k\, i_k}{\sum_k \ell_k}.$$

The registry began with 99 candidates. Eleven non-target chimeric or non-opioid constructs, 7T10, 7T11, 7UL2, 7UL3, 7UL5, 7YMJ, 8HNN, 8K2W, 9LFA, 9LFC, and 9LFD, and two mouse receptor structures, 9MQK and 9MQL, were excluded. The retained cohort contains 86 structures. The selected receptor copy was identified by receptor entity, followed by exact author-chain agreement, exact label-chain agreement, and lexical identifiers. The selected receptor chain in the RCSB asymmetric unit, the deposited coordinate unit before biological-assembly transformations, supplied primary receptor graphs. Biological assembly 1, the deposited macromolecular assembly used here for environmental context, supplied component, partner, lipid, ion, and water coordinates; that retained the receptor copy, complete contacting polymer chains, and individual nonpolymer or water residues with a heavy atom within 5.0 Å of the receptor.

Two pairs share experimental maps. PDB entries 10TL and 9PU5 share EMD-71869 and both have nominal resolution 3.5 Å. Entries 10TM and 9PUD share EMD-71871 and both have nominal resolution 3.0 Å. The receptor graphs within each pair have contact Jaccard similarity, the size of the contact-set intersection divided by the size of its union, equal to 1.0; they also have zero gained or lost contacts and zero shared-contact root-mean-square (RMS) distance deviation. The explicitly designated independent-map representatives were 9PU5 and 9PUD because they are the complete receptor- $G_i$ entries, where $G_i$ denotes the inhibitory heterotrimeric G-protein family. Both alternative deposited identifiers remain in the cohort registry. This rule yields 84 independent maps.

GPCRdb state annotation had precedence. GPCRdb Active entries with a populated signaling-protein type were active-partner-bound, Active entries without that field were active-like, and Inactive entries were inactive-like. Intermediate or Other annotations would have retained their corresponding categories but were absent. If GPCRdb state was unavailable, the normalized lowercase RCSB title was evaluated in a fixed order. An inactive, antagonist, or inverse agonist token assigned inactive-like. Otherwise, a g protein, -gi, gi , goa, gz, or arrestin token assigned active-partner-bound. Otherwise, an active or agonist token assigned active-like. A title not meeting these rules was unclear. GPCRdb annotation overrode a conflicting title; 8Y45 was the only retained conflict. The resulting counts are 49 active-partner-bound, 16 inactive-like, 4 active-like, and 15 unclear maps. Evidence sources were GPCRdb/title for 20/29 active-partner-bound maps, 8/8 inactive-like maps, 4/0 active-like maps, and 0/15 unclear maps. State models used only active-partner-bound and inactive-like maps.

Ligand and component roles were curated before statistical analysis. Orthosteric denotes binding at the receptor's primary ligand pocket, whereas allosteric denotes binding at a distinct site. The primary 84-map binding partition is 72 orthosteric-only, 9 orthosteric-plus-allosteric, 1 allosteric-only, and 2 with no resolved ligand. Named short opioid peptides were accepted as orthosteric ligands when chain identity and receptor contact supported that role. The selected-ligand role is the curated union of orthosteric and allosteric targets; its coordinate-defined receptor site is the set of exact-BW residues within 4.5 Å of those targets. In 9L60, chain P is dynorphin A(1-13), whereas A1D6C, MPAM-15, remains allosteric. PIO in 9ZZO is deposited short-chain di$C_8$-phosphatidylinositol 4,5-bisphosphate, abbreviated PI(4,5)$P_2$ or $PIP_2$, and was assigned to the analysis category $PIP_2$-like lipid. Cohort decisions, curated roles, feature prevalence, and mapping quality control are summarized in Tables S1 and S2 and Figure S1.

##### S2. Exact target-core mapping and receptor-contact graphs

The expected-accession target core contains selected-chain polymer residues explicitly covered by the expected receptor's SIFTS segment. This broader core retained 24,360 residues and excluded 715 selected-chain residues. Among exclusions, 489 mapped explicitly to non-target UniProt accessions and 226 lacked an explicit mapping to the expected target accession. Coordinate residues were assigned to the expected accession only by exact selected-chain SIFTS correspondence and then to an exact GPCRdb generic label. No alternative coordinate-chain mapping or heuristic segment-fraction label was used. All 39,914 propagated mapping records, comprising 19,957 asymmetric-unit records and their biological-assembly-1 counterparts, were supported by the PDBe SIFTS range and GPCRdb generic-label sources. Exact-BW graph nodes are a narrower rule-specific subset of the retained target core: a primary node must have an exact transmembrane helix 1 through 7 (TM1 through TM7) GPCRdb/BW label and a resolved alpha-carbon (Cα) atom. Across the 86 primary Cα graphs, this subset comprised 19,957 structure-node records. Fusion proteins, antibodies, nanobodies, G proteins, arrestins, and other components were not receptor-graph nodes, although their coordinate relationships were retained as context.

For structure $s$ and contact definition $m$, the receptor graph was an undirected simple graph, meaning that contacts have no direction, self-loops are excluded, and each residue pair appears once:

$$G_s^{(m)} = \left(V_s, E_s^{(m)}\right).$$

$V_s$ contains exact-BW target-receptor residues with the atoms required by rule $m$. The primary Cα 8 Å edge set was

$$E_s^{(\mathrm{C}\alpha 8)} = \left\{\{u, v\}: u, v \in V_s,\ u < v,\ \left\|\boldsymbol{x}_{su}^{(\mathrm{C}\alpha)} - \boldsymbol{x}_{sv}^{(\mathrm{C}\alpha)}\right\|_2 \leq 8.0\,\text{Å}\right\}.$$

Each unordered residue pair was included once, with no sequence-separation exclusion. Sensitivity rules were Cα at 6.0 Å, Cα at 10.0 Å, beta-carbon (Cβ) at 8.0 Å with Cα used for glycine, minimum inter-residue nonhydrogen-atom distance at 4.5 Å, and minimum side-chain nonhydrogen-atom distance at 5.0 Å. For glycine in the side-chain rule, Cα represented the residue. A residue lacking the atom set required by a rule was absent from that rule's node or contact calculation rather than imputed. The selected coordinate model was the first deposited model. An alternate location is one of multiple deposited coordinate records for the same atom. DOR 4N6H was the only selected-receptor coordinate file with alternate locations; its A and B conformers had equal occupancy, meaning the same deposited fractional population, and conformer A was selected. No selected chain contained a duplicated exact-BW label. Graph construction was defined to fail closed, meaning to stop rather than choose among competing residues, if a duplicate label occurred.

The primary graph domain was the exact-BW-induced target core. The full selected receptor core was analyzed separately as a domain sensitivity. The six contact rules and two graph domains were never averaged. The primary claims use only the exact-BW Cα 8 Å graph.

**S3. Structural alignment, pairwise contact comparison, and endpoint means**

All-to-all comparison means that every unordered structure pair is enumerated once. The $n(n-1)/2$ analysis contained 3,655 pairs for 86 structures, 3,486 for 84 independent maps, and 2,926 for the 77-map selected-ligand-site subset. For structures $a$ and $b$, $M_{ab}$ denotes shared exact-BW positions with Cα coordinates. All pairs had more than 200 shared positions. Mapped coordinates were superposed by Kabsch rigid-body alignment constrained to a proper rotation. Root-mean-square deviation (RMSD) is the residual coordinate difference after that superposition and was calculated as follows.

$$\mathrm{RMSD}_{ab} = \min_{Q,t} \sqrt{\frac{1}{|M_{ab}|} \sum_{r \in M_{ab}} \|Q\boldsymbol{x}_{ar} + \boldsymbol{t} - \boldsymbol{x}_{br}\|_2^2}.$$

$Q$ is a proper rotation with determinant 1, $\boldsymbol{t}$ is a translation, and $\boldsymbol{x}_{ar}$ and $\boldsymbol{x}_{br}$ are mapped Cα coordinate vectors. Positions missing from either structure were excluded from $M_{ab}$.

An estimand is the precisely specified quantity targeted by a calculation. For structure $s$, $E_s$ contained every contact whose two endpoints had exact-BW assignments in that structure. The primary estimand induced both graphs on the exact-BW nodes available in the pair:

$$E_{a|ab} = \{\{u, v\} \in E_a : u \in M_{ab} \wedge v \in M_{ab}\},$$

$$E_{b|ab} = \{\{u, v\} \in E_b : u \in M_{ab} \wedge v \in M_{ab}\}.$$

For these node-comparable edge sets, superscript $c$ denotes complement within $U_{ab}$:

$$S_{ab} = E_{a|ab} \cap E_{b|ab}; \qquad U_{ab} = E_{a|ab} \cup E_{b|ab},$$

$$J_{ab} = \frac{|S_{ab}|}{|U_{ab}|}; \qquad d_{J,ab} = 1 - J_{ab},$$

$$E_{\text{gain}} = E_{b|ab} \cap E^{c}_{a|ab}; \qquad E_{\text{loss}} = E_{a|ab} \cap E^{c}_{b|ab}; \qquad E_{\text{cons}} = S_{ab},$$

$$R_{ab} = |E_{\text{gain}}| + |E_{\text{loss}}| = \left|E_{a|ab} \,\triangle\, E_{b|ab}\right|.$$

$R_{ab}$ is termed the rewired-contact count as shorthand for a static contact symmetric difference. It does not measure a physical transition. No primary edge union was empty. A full-edge-set sensitivity estimand allowed a contact incident on a node unavailable in the other structure to enter the symmetric difference. Let $N_{\text{full}}$ and $N_{\text{shared}}$ be the pooled full-edge and shared-node changed-contact totals within one declared scope. Mapping-availability retention and removal were

$$f_{\text{retained}} = \frac{N_{\text{shared}}}{N_{\text{full}}}; \qquad f_{\text{removed}} = \frac{N_{\text{full}} - N_{\text{shared}}}{N_{\text{full}}} = 1 - f_{\text{retained}}; \qquad P_{\text{removed}} = 100 f_{\text{removed}}.$$

They were undefined when $N_{\text{full}} = 0$. Shared-node coverage ratios and the difference between the primary and full-edge estimands were retained as quality-control variables. The shared-contact distance RMS deviation was

$$D_{ab} = \sqrt{\frac{1}{|S_{ab}|} \sum_{e \in S_{ab}} [d_b(e) - d_a(e)]^2}.$$

For conserved edge $e$, $d_a(e)$ and $d_b(e)$ are the contact-rule-specific distances between its residue endpoints in structures $a$ and $b$. $D_{ab}$ was missing if no conserved edge was available; this case did not affect the reported results.

Pairwise outcomes reuse structures. For eligible structure i, the endpoint mean was

$$\bar{Y}_i = \frac{1}{n_i} \sum_{j \in P_i} Y_{ij},$$

where $P_i$ is the set of eligible comparison partners for structure $i$ and $n_i = |P_i|$. In the complete 84-map universe $n_i = 83$; in the complete 77-map selected-site universe $n_i = 76$. Scope restrictions and missing outcomes changed $n_i$ explicitly. A leave-one-out calculation deletes one analysis unit, removes every pairwise record involving it, recomputes the remaining endpoint means, and refits the model.

For the endpoint-outcome redundancy analysis, $x_i$ was endpoint-mean contact Jaccard similarity and $y_i$ was endpoint-mean rewired-contact count for independent map $i$. Pearson $r$, the linear-correlation coefficient, across the $N = 84$ complete paired endpoint means was

$$r = \frac{\sum_{i=1}^{N}(x_i - \bar{x})(y_i - \bar{y})}{\sqrt{\left[\sum_{i=1}^{N}(x_i - \bar{x})^2\right]\left[\sum_{i=1}^{N}(y_i - \bar{y})^2\right]}}.$$

This correlation quantifies linear redundancy between two primary shared-node outcomes and is not independent biological evidence.

An ancillary semantic-transport analysis, a nonphysical comparison of contact distributions across predefined categorical bins, encoded exact-mapped contact rows in either family or universal semantic space. GPCR family bins combined GPCR pair class, helix-segment pair, and receptor-contact context. Universal bins combined generic contact class, structural segment pair, and intra-chain or inter-chain context. In burden mode, bin mass was proportional to contact-row count. In binary mode, every occupied bin received equal mass:

$$m_k^{(\text{burden})} = \frac{n_k}{\sum_j n_j}; \qquad m_k^{(\text{binary})} = \frac{1}{K_{\text{occ}}}.$$

Here, $n_k$ is the number of contact rows assigned to bin $k$ and $K_{\text{occ}}$ is the number of occupied bins. The four family/universal by burden/binary settings yielded 14,620 records. They were retained separately and are ancillary to the primary biological claims.

##### S4. Ligand, component, and protein-interface estimands

Assembly presence, receptor contact, exact-BW interface, selected-ligand role, and curated mechanistic role were distinct variables. Assembly presence records whether a component is modeled, receptor contact records physical proximity, an exact-BW interface requires a mapped receptor contact, selected-ligand role identifies the curated orthosteric or allosteric target, and mechanistic role records the literature-supported interpretation. Nonpolymer instances were matched by chemical component identifier, chain, and residue identifier; polymeric peptide ligands were matched by curated chain identity. In the component and water coordinate curation, repeated nonhydrogen atoms with the same atom name were ranked by blank alternate-location label, then label A, then other labels; occupancy was ranked from greatest to least within each label tier, followed by the label as a deterministic tie breaker. For component $c$ and exact-BW receptor residue $v$ in structure $s$, the coordinate-defined site at cutoff $\delta$ was

$$S_{sc}(\delta) = \left\{ v \in V_s : \min_{a \in c,\, b \in v} \lVert \boldsymbol{x}_a - \boldsymbol{x}_b \rVert_2 \leq \delta \right\}.$$

The minimum was taken over nonhydrogen atoms. $\delta = 4.5$ Å defined primary ligand, lipid, sterol, allosteric-modulator, and protein-interface sites. The selected-ligand site was the union of coordinate-defined sites for role-selected orthosteric and allosteric targets. A missing target or unresolved contact site remained unknown and was not coded as absence. Protein-Ligand

Interaction Profiler (PLIP) exact-BW interactions were retained as a separate interaction-rule comparator.

This distinction matters for the 9L60 dynorphin/MPAM-15 complex. PLIP maps 14 dynorphin exact-BW contacts. The primary 4.5 Å coordinate rule maps 25 dynorphin positions, 10 MPAM-15 positions, and 35 positions in their role-aware union. Article Figures 9 and 10 and the spatial analysis use the 35-position coordinate-defined union.

Protein-interface residue identifiers were resolved to the designated receptor chain before exact-BW mapping. Gα, Gβ, and Gγ denote the alpha, beta, and gamma subunits of a heterotrimeric G protein. Their assembly vectors each equal the resolved G-protein assembly vector across the 84-map panel, with 66 positive maps. The Gβ-specific exposure therefore asks whether any Gβ heavy atom is within 4.5 Å of the full designated receptor within those 66 Gα-positive maps. Eleven maps meet this definition, 15 meet a 5.0 Å sensitivity, and 3 have a mapped exact-BW Gβ interface. No Gγ residue met the primary receptor-contact criterion.

**S5. Site distance and direct, adjacent, connected-distal, and disconnected classes**

For a resolved site $S_s$ in graph $G_s$, node and edge distances were

$$h_s(v; S_s) = \min_{u \in S_s} \{ \mathrm{dist}_{G_s}(v, u) \},$$

$$h_s(\{u, v\}; S_s) = \min\{ h_s(u; S_s), h_s(v; S_s) \}.$$

An edge was direct when $h = 0$, adjacent when $h = 1$, connected-distal when $1 < h < \infty$, and disconnected when no path existed. A lost edge was classified in the source structure when that site was resolved and otherwise in the destination structure. A gained edge was classified in the destination structure when that site was resolved and otherwise in the source structure. A conserved edge received the most proximal supported classification across resolved endpoint sites. A pair with neither endpoint site resolved was omitted. The primary selected-ligand pair analysis required both endpoint sites to be resolved.

For class $k$, the pooled static and changed-contact fractions were

$$f_k^{(\mathrm{wire})} = \frac{\sum_s |E_{sk}|}{\sum_s |E_s|},$$

$$f_k^{(\mathrm{rewire})} = \frac{\sum_{a<b} R_{abk}}{\sum_{a<b} R_{ab}}.$$

$E_{sk}$ is the subset of structure $s$ edges assigned to class $k$, and $R_{abk}$ is the number of changed contacts assigned to $k$ for pair $\{a, b\}$. The reported percentage forms were $\pi_k = 100 f_k^{(\mathrm{wire})}$ and $\rho_k = 100 f_k^{(\mathrm{rewire})}$. A fraction and its percentage were undefined when their corresponding pooled denominator was zero. The values therefore weight edges or changed contacts, not maps equally. Selected-ligand union sites are resolved in 79/86 structures and 77/84 independent maps. Every Cα 8 Å receptor graph in the 77-map subset is connected, so the disconnected fraction is zero under the primary definition and distal means connected-distal.

**S6. Deposited-water estimands, water graph, and recurrence**

A deposited water is a water molecule represented explicitly by coordinates in a deposited structural model. Recognized water residue names were HOH, WAT, $H_2O$, and DOD. Occupancy is the deposited fractional population assigned to an atomic coordinate record. Among oxygen records in one water residue, the representative was the record with greatest occupancy; an equal-occupancy tie retained deposition order. Receptor proximity used any designated-receptor heavy atom within 5.0 Å. Polar contacts used nitrogen (N) and oxygen (O) atoms within the selected cutoff. At the default 3.5 Å polar cutoff and occupancy threshold 0.7, let $o(w)$ be occupancy, $d_R(w)$ the minimum receptor distance, $A(w)$ the set of protein-residue polar partners, $R(w)$ the subset of designated-receptor partners, $B(w)$ the subset of exact-BW receptor partners, and $l(w)$ indicate a selected-ligand polar partner. The estimands were

$$P = \{w: o(w) \geq 0.7 \wedge d_R(w) \leq 5.0\,\text{Å}\},$$

$$C = \{w \in P: |A(w)| \geq 2\},$$

$$R_2 = \{w \in C: |R(w)| \geq 2\},$$

$$B_{\text{any}} = \{w \in P: |B(w)| \geq 1\},$$

$$C_{\text{B1}} = C \cap B_{\text{any}}; \qquad C_{\text{B2}} = \{w \in C: |B(w)| \geq 2\},$$

$$L = \{w \in P: l(w) = 1 \wedge |B(w)| \geq 1\}.$$

$A(w)$ can include receptor, Gα, Gβ, Gγ, β-arrestin, a nanobody, a single-chain variable fragment (scFv), an antigen-binding fragment (Fab), or other stabilizing protein residues. A nanobody is a single-domain antibody fragment, an scFv joins antibody variable domains in one polypeptide, and a Fab is the antigen-binding antibody fragment. $R(w)$ and $B(w)$ are narrower receptor subsets. $C$, $R_2$, and $B_{\text{any}}$ are subsets of $P$; $R_2$ is also a subset of $C$ because $R(w)$ is a subset of $A(w)$; $C_{\text{B2}}$ is a subset of $C_{\text{B1}}$, which is a subset of $C$; and $L$ is a subset of $B_{\text{any}}$. $L$ is not required to be a subset of $C$. Occupancy sensitivities were 0, 0.3, 0.5, 0.7, 0.9, and 1.0. Polar cutoffs were 3.2, 3.5, 3.8, and 4.1 Å.

For optional water-network descriptors, nodes were receptor-proximal water instances with at least one exact-BW polar partner, and two nodes were joined when their oxygen distance was at most the current polar cutoff. For $H = (W, F)$ with $\kappa$ connected components, cycle rank was

$$\mu = |F| - |W| + \kappa.$$

A water-network hub had degree at least 3. These descriptors were not used to infer dynamic water residence or energetics.

Water coordinates were not compared in raw deposited frames. Primary coordinate clustering, one-to-one coordinate matching, and the reported exact-BW bridge-topology recurrence used the 38 default $C_{\mathrm{B2}}$ waters in 7 maps unless another set was named. The alignment reference was selected once from the full 84-map panel as the map with the greatest number of exact-BW receptor Cα coordinates, with lexicographic structure identifier as the tie breaker. DOR 4N6H and DOR 4RWD each had 240 positions, so 4N6H was selected. Every other map was Kabsch-aligned to 4N6H over their shared exact-BW receptor Cα positions, and the same transformation was applied to its water oxygens. Coordinate radii were 0.8, 1.0, 1.2, 1.5, and 2.0 Å. Complete linkage required every within-cluster separation to be at most the radius. Single-linkage clusters were connected components after joining waters within the radius. A cluster was recurrent when it contained water instances from at least two independent maps. Pairwise recurrence used maximum-cardinality one-to-one matching, which selects the greatest possible number of nonreused cross-map water pairs whose oxygen separation does not exceed the stated radius. Each $C_{\mathrm{B2}}$ water contributed every unordered pair among its exact-BW partners; a topology was recurrent when the same pair was contributed by at least two independent maps. The primary leave-4N6H-out analysis filtered its waters after alignment and retained the common 4N6H coordinate frame. A reference-selection sensitivity applied the same deterministic rule after excluding 4N6H, selected 4RWD with 240 exact-BW coordinates, and aligned the remaining maps to that reference. At 1.0 Å, both frames yielded three complete-linkage recurrent clusters and three single-linkage recurrent clusters; 2 of 15 map pairs had at least one one-to-one coordinate match in both frames. Exact-BW topology recurrence is reference-free. PLIP water events were coordinate-matched to curated waters within 0.10 Å, deduplicated by structure/water/BW endpoint, and treated as a separate operational definition.

##### S7. Prevalence, percentages, and bootstrap procedures

For $x$ positive maps among $n$, prevalence was $\hat{p} = x/n$. The two-sided 95% Wilson confidence interval (CI), a binomial-proportion interval with improved boundary behavior relative to the simple normal approximation, used $z = 1.959963984540054$.

$$\mathrm{CI}_W = \frac{\hat{p} + \frac{z^2}{2n} \pm z\sqrt{\frac{\hat{p}(1-\hat{p})}{n} + \frac{z^2}{4n^2}}}{1 + \frac{z^2}{n}}.$$

Wilson endpoints were computed on the unit-proportion scale and reported in percent as $\mathrm{CI}_W(\%) = 100\,\mathrm{CI}_W$.

Denominators were 84 maps for sterol, generic lipid, deposited water, structural water, G-protein assembly, allosteric contact, sodium, and ligand-water bridging; 76 resolved-modality maps for peptide versus small molecule; 81 resolved-binding maps for orthosteric-plus-allosteric versus orthosteric-only; and 66 Gα-positive maps for direct Gβ contact.

For outcome $Y$, the exposed group, denoted $E$, contained maps with the specified component or context, and the reference group, denoted $R$, contained the prespecified comparison maps without that exposure. The unadjusted percentage contrast was

$$\Delta_{\text{unadj}}(\%) = 100\,\frac{\bar{Y}_E - \bar{Y}_R}{|\bar{Y}_R|}.$$

The percentage was undefined when $|\bar{Y}_R| \leq 10^{-15}$. Article Table 3 and SI Table S5 use only the 77 selected-ligand-site-positive maps. Unadjusted intervals in SI Table S5 used a stratified fixed-map-record bootstrap. For a pairwise outcome, one endpoint mean was first computed for every map over its 76 observed pairwise records. For an intrinsic static outcome, the observed per-map fraction was used directly. Each of 5,000 replicates sampled exposed and reference map records separately with replacement at their observed group sizes while carrying the relevant fixed map value unchanged. Pair rows were not resampled, the pairwise comparison graph was not rebuilt, and endpoint means were not recomputed. Interval limits were the empirical 2.5th and 97.5th percentiles and are conditional on the observed map summaries.

For estimable structure contrasts in the separate component percentage-effect tables, 10,000 independent resamples were drawn. Publication clustering attempted 10,000 publication-key resamples with replacement and retained every map belonging to each selected key occurrence; percentile limits used only replicates containing both exposure levels and an absolute reference mean greater than $10^{-15}$. For matched pair $\ell$, the pair-level percentage and the mean across valid pairs were

$$\Delta_{\text{pair},\ell}(\%) = 100\,\frac{Y_{E,\ell} - Y_{R,\ell}}{|Y_{R,\ell}|}; \qquad \bar{\Delta}_{\text{match}}(\%) = \frac{1}{n_{\text{valid}}}\sum_{\ell\in\mathcal{V}} \Delta_{\text{pair},\ell}(\%).$$

A pair belonged to $\mathcal{V}$ only when both outcomes were finite and $|Y_{R,\ell}| > 10^{-15}$; $n_{\text{valid}} = |\mathcal{V}|$, and the summary was undefined when $n_{\text{valid}} = 0$. Matched-pair percentage intervals used 10,000 bootstrap resamples of the valid pair-level percentages. Structure, publication, and matched-pair seeds were deterministic integers derived from Secure Hash Algorithm 256-bit (SHA-256) cryptographic hashes of the contrast identifier, outcome identifier, and corresponding resampling-mode tag.

**S8. State and component models, covariance, and multiplicity**

State analysis used inactive-like and active-partner-bound maps only. The state-model adjustment block was

$$A_i^{\text{state}} = \alpha_{\text{subtype}(i)} + \eta_{\text{method}(i)} + \beta_r z(\text{resolution}_i) + \beta_m z(\text{mapping}_i) + \beta_b z(\text{BWcount}_i).$$

For outcome $Y_i$, the fitted model was

$$Y_i = \beta_0 + \beta_I I(\text{inactive}_i) + A_i^{\text{state}} + \epsilon_i.$$

$I(\text{inactive}_i)$ is 1 for an inactive-like map and 0 for an active-partner-bound map. The $\alpha$ and $\eta$ blocks are subtype and method effects, $z$ denotes within-scope standardization, $\beta$ coefficients are fitted fixed effects, and $\epsilon_i$ is the residual. Active-partner-bound is the reference. Mapping is receptor mapping coverage, and BWcount is the number of exact-BW-mapped receptor residues. Sixteen estimable outcomes formed one defined positive-control family. Tests were two-sided. The positive-control expectation was lower similarity and greater pairwise structural difference for inactive-like maps; local direct, adjacent, and connected-distal measures were included to locate differences rather than to impose a signed rejection rule. Structurally zero disconnected outcomes were not estimable and were not tested.

The state-adjusted percentage was

$$\Delta_{\text{state}}(\%) = 100\,\frac{\beta_I}{|\bar{Y}_{\text{APB}}|},$$

where $\bar{Y}_{\text{APB}}$ is the observed active-partner-bound mean in the fitted scope. Coefficient confidence limits were scaled by the same denominator.

For each component contrast, the nonexposure adjustment block was

$$A_s^{\text{component}} = \alpha_{\text{subtype}(s)} + \gamma_{\text{state}(s)} + \eta_{\text{method}(s)} + \beta_r z(\text{resolution}_s) + \beta_m z(\text{mapping}_s) + \beta_b z(\text{BWcount}_s).$$

The generic component model was

$$\bar{y}_s = \beta_0 + \beta_{\text{exp}} X_s + A_s^{\text{component}} + c_s^{\mathsf{T}}\theta + \epsilon_s.$$

$\bar{y}_s$ is the endpoint mean for a pairwise outcome or the observed value for an intrinsic outcome. $X_s$ is the binary exposure indicator, $\beta_{\text{exp}}$ is its exposed-minus-reference coefficient, $\gamma$ is the state-effect block, $c_s$ is the contrast-specific context vector, and $\theta$ contains its coefficients. Collecting all nonexposure adjustment terms gives $\Gamma^{\mathsf{T}} Z_s$, where $Z_s$ is the covariate vector and $\Gamma$ is its coefficient vector.

Mapping coverage was the maximum SIFTS-reported coverage among the intervals for the selected expected-accession chain record. Mapped-BW count was the number of nodes in that map's primary exact-BW Cα graph. For a continuous covariate x in an H-map analysis scope, standardization was

$$z_i(x) = \frac{x_i - \bar{x}_H}{\sqrt{\frac{1}{H}\sum_{j=1}^{H}(x_j - \bar{x}_H)^2}}.$$

If the population standard deviation was zero, the scale was set to 1 after centering, so all observed values of the constant covariate mapped to 0. This safeguard did not impute a missing covariate.

Categorical variables used treatment coding, meaning that each nonreference level was represented by an indicator relative to one omitted reference level. After levels represented by fewer than 5 maps within the scope were combined as other_sparse, the lexicographically first retained level was the reference; these references were DOR for subtype, active-partner-bound for state after the four active-like maps were combined as other_sparse, electron microscopy for method, orthosteric for curated binding mode, and other-or-unresolved for modality when that category was included. A missing categorical value was labeled unknown before sparse-level collapsing. Binary G-protein and allosteric-contact indicators used 0 as reference. Rows missing the outcome, exposure, resolution, mapping coverage, or mapped-BW count were excluded from that model. A model was retained only when both exposure levels contained at least 5 maps and the design matrix was full rank, meaning that no included design column was an exact linear combination of the others.

H1 through H4, sterol, generic lipid, deposited water, and structural water, additionally included curated binding mode and G-protein context. H5, G-protein assembly, included binding mode but omitted a separate G-protein covariate. H6, direct Gβ contact, was restricted to Gα-positive maps, included binding mode, and omitted the G-protein covariate. H7, peptide versus small molecule, was restricted to 76 resolved-modality maps, omitted binding mode, and included receptor-contacting allosteric-modulator and G-protein indicators. H8, orthosteric-plus-allosteric versus orthosteric-only, was restricted to 81 resolved-binding maps, omitted binding mode, and included orthosteric modality and G-protein context.

The four outcomes were exact-BW contact-pair count, endpoint-mean contact Jaccard similarity, endpoint-mean rewired-contact count, and endpoint-mean shared-contact distance RMS deviation. All $8 \times 4 = 32$ models were estimable and full rank. Article Table 3 displays, for each contrast, the outcome with the smallest nominal $p$ value among its four defined outcomes. This is an exploratory display rule; inference uses all 32 rows.

Ordinary least squares used type 3 heteroskedasticity-consistent (HC3) covariance and a $t$ reference distribution. With design matrix $X$, ordinary least-squares estimate $\hat{\boldsymbol{\beta}}$, residual $e_i$, and leverage $h_{ii}$, the $i$th diagonal element of the regression hat matrix,

$$V_{\text{HC3}}(\hat{\boldsymbol{\beta}}) = (X^{\mathsf{T}}X)^{-1}X^{\mathsf{T}}\text{diag}\left[\frac{e_i^2}{(1-h_{ii})^2}\right]X(X^{\mathsf{T}}X)^{-1}.$$

Here, diag forms a diagonal matrix from the bracketed residual terms. Confidence intervals used residual degrees of freedom. The adjusted percentage was

$$\Delta_{\text{adj}}(\%) = 100\,\frac{\beta_{\text{exp}}}{|\bar{Y}_R|},$$

where $\bar{Y}_R$ is the observed reference-group mean within the fitted scope, not a model-predicted counterfactual. Coefficient confidence limits were scaled by the same denominator. Publication-cluster covariance was a sensitivity calculation when at least 10 publication groups were available.

For ordered $p$ values $p_{(1)} \le \cdots \le p_{(m)}$, Benjamini-Hochberg adjusted $p$ values, denoted $q$, were calculated as follows.

$$q_{(i)} = \min\left\{1, \min_{j\ge i}\left(\frac{m}{j}\right)p_{(j)}\right\}.$$

$m = 16$ for the state family and $m = 32$ for the component family. The families were not combined. Zero of 32 component tests had $q < 0.05$.

#### S9. Matching, leave-one-out analysis, ancillary calculations, and reproducible specification

**Normalization, scaling, and denominator registry.** No single normalization was applied across outcomes. For pair $\{a, b\}$, Jaccard union normalization divided conserved contacts by the pairwise union, $J_{ab} = |S_{ab}|/|U_{ab}|$, and the complementary Jaccard distance satisfied

$$1 - J_{ab} = \frac{R_{ab}}{|U_{ab}|}.$$

Shared-node induction was not a numerical normalization. It restricted both endpoint graphs to BW positions resolved in both structures before their edge sets were compared. The rewired-contact quantity $R_{ab}$ remained a raw, unnormalized symmetric-difference count. It was reported together with union-normalized Jaccard measures, and regression models included mapped-BW count to adjust for graph-size differences. This covariate adjustment does not mathematically normalize $R_{ab}$. Shared-contact distance RMS deviation used the conserved-contact set as its root-mean-square denominator. An unresolved site, missing outcome, or unavailable node was excluded from its stated denominator and was not converted to zero.

Endpoint averaging divided the sum of eligible incident pairwise outcomes by the number $n_i$ of eligible partners, thereby producing one model record per map within a stated scope. Pooled static wiring fractions divided the total number of class-specific edges by the total number of pooled edges. Pooled rewiring fractions divided class-specific changed contacts by all pooled changed contacts. These are contact-weighted fractions, not equally weighted map averages. Unadjusted percentage contrasts divided the exposed-minus-reference mean difference by the absolute observed reference-group mean. Adjusted percentage contrasts divided the fitted exposure coefficient by that same type of observed reference-group mean within the fitted scope. These percentage transformations scale effects and do not normalize probabilities.

Continuous covariates used analysis-scope z-standardization by subtraction of the scope mean and division by the population standard deviation; after centering, a zero population standard deviation was replaced by 1, and missing values were not imputed. Categorical covariates were not z-standardized. The 78 graph-fingerprint fields were centered and scaled across all 86 structures before principal component analysis (PCA), with scale 1 assigned to a zero-variance field. Fingerprint spectral fields used the unweighted symmetric normalized Laplacian. Current-flow node and edge betweenness used conductance weights, the largest connected component, and the stated algorithmic graph-size normalization. The perturbation-response scanning (PRS) matrix was normalized separately by dividing each row by its diagonal self-response, with denominator 1 used only when that diagonal was nonpositive. StrucFlow instead used an unnormalized weighted Laplacian. In the contact-propagation network (CPN), weighted degree was divided by the map-specific maximum degree, evidence means used their explicitly eligible row weights, and each floored pair-potential matrix was divided by its own $3 \times 3$ matrix mean. These graph-specific transformations have different targets and are not interchangeable.

Matching was exact on experimental method, receptor subtype, and state. Resolution, mapping coverage, and mapped-BW count were standardized once across the complete hypothesis-specific analysis scope by the population mean and population standard deviation. Within an exact stratum, exposed and reference maps were paired without replacement and without a distance caliper, meaning no maximum allowable within-pair distance was imposed, by minimum-cost linear assignment. Distance was

$$d(i,j) = \sqrt{\sum_{k=1}^{3} \left(z_{ik} - z_{jk}\right)^2}.$$

Matched differences used full enumeration of all $2^n$ sign assignments when the number of pairs $n$ was at most 18. A sign assignment reverses or retains the direction of each paired difference under the paired null. Otherwise, the test used 100,000 seeded Monte Carlo sign assignments, meaning random draws from those possible sign assignments, and applied a plus-one correction to the Monte Carlo tail probability only. The statistic was the absolute mean paired difference. In the analyzed contrasts, every matched test used full enumeration. The Monte Carlo fallback seed was derived from the hypothesis identifier, outcome identifier, and prespecified method identifier. Publication identity was assigned by PubMed identifier, then persistent digital publication identifier, normalized title, and PDB accession as fallback. A within-publication contrast required both exposure levels within the same publication. For each leave-one-out analysis, the unit and every pairwise record involving it were excluded, endpoint means were calculated from the remaining pairwise records, and the specified model was fitted. The same full endpoint-recomputation procedure was applied to the six retained state outcomes; all eligible deletions completed and all six coefficients preserved their signs.

**Graph attributes, fingerprints, dimensional reduction, clustering, and communities.** Each receptor graph carried nine scalar residue descriptors: dynamic coupling index (DCI), a contact-energy proxy, weighted constraint coordination, rigidity, travel depth, side-chain entropy, dehydron count, buried unsatisfied polar count, and local backbone entanglement. Helix, sheet, and coil membership supplied three binary indicators. For a 13 Å Cα elastic-network Hessian pseudoinverse $H^+$, define the off-diagonal mean block response as

$$\bar{r}_{\text{off}} = \frac{1}{N(N-1)} \sum_{p \neq q} \left\| (H^+)_{pq} \right\|_F.$$

The operational dynamic-coupling score was

$$\text{DCI}_i = \frac{\frac{1}{N-1} \sum_{j=1}^{N} \left\| (H^+)_{ij} \right\|_F}{\bar{r}_{\text{off}}}.$$

Thus, the row numerator included the diagonal self-response block, whereas its scale factor and the reference mean used $N-1$ and off-diagonal blocks, respectively. The contact-energy descriptor summed a Lennard-Jones plus Coulomb proxy over heavy-atom pairs separated by strictly less than 6 Å whose residue indices differed by at least two, replaced pair distances below 2.2 Å by 2.2 Å, used distance-dependent dielectric $4r$, clipped each pair contribution to the interval from $-12$ to $12$, and assigned half of each pair value to each endpoint. Weighted constraint coordination was the sum of rule-defined constraints incident on a residue. The geometric rules used a 2.0 Å backbone carbon-to-nitrogen cutoff, 4.5 Å minimum-heavy-atom cutoff, and 8.0 Å Cβ cutoff; each base constraint had weight 1, with an increment of 1 for a backbone nitrogen-to-oxygen hydrogen bond within 3.5 Å and an increment of 1 for a salt bridge within 4.0 Å. Rigidity was $\min(1, c_i/3)$, where $c_i$ is weighted coordination. Travel depth used a 1.5 Å voxel grid, a 1.4 Å solvent probe, and shortest travel to bulk solvent. Side-chain entropy was the residue-specific maximum entropy multiplied by $1 - \min(1, n_{10}/24)$, where $n_{10}$ is the number of neighboring residues within 10 Å. A dehydron was a backbone hydrogen bond with fewer than 19 surrounding carbon atoms within 6.5 Å. A buried unsatisfied polar site had exposure at most 0.15 and no nitrogen or oxygen partner within 3.5 Å. Entanglement used a discrete Gauss-integral contribution over nonadjacent Cα bond segments.

The seven directed edge descriptors were residue distance, sequence separation, interchain status, contact-geometry surprisal, contact-density percentile, and the secondary-structure codes of the two endpoints. Each undirected contact supplied two directed rows, so $m_d = 2|E|$. Missing or nonfinite edge values used the following fixed defaults: 8.0 Å for residue distance, 0 for sequence separation and interchain status, 0 bits for contact-geometry surprisal, 0.5 for contact-density percentile, and coil code 2 for either secondary-structure endpoint. The contact-geometry null used 18 equal-area $\theta$ bins, 36 uniform $\phi$ bins, add-0.5 smoothing over 648 bins, and global backoff when a secondary-structure-pair context contained fewer than 200 reference contacts. Surprisal was $-\log_2$ of the smoothed bin probability, and density percentile was the total probability mass of bins no more probable than the observed bin. The graph fingerprint concatenated the mean, population standard deviation, maximum, and minimum of each of the 12 node fields; mean, population standard deviation, maximum, and median degree; the first 16 eigenvalues of the unweighted symmetric normalized Laplacian; the mean of each edge field; $\log(1 + n)$; $\log(1 + m_d)$; and directed density $m_d/[n(n-1)]$. The resulting 78 dimensions were finite. Each field was centered by its all-86-structure mean and divided by its population standard deviation; a zero-variance field used unit scale and therefore remained 0 after centering. Principal component analysis was fitted to all 86 structure fingerprints, and the 84 independent-map representatives were displayed. Ward minimum-variance clustering used Euclidean distance at $k = 4$ and $k = 8$. These coordinates and partitions are descriptive and are not state or ligand classifiers.

Louvain communities were calculated on each unweighted undirected receptor graph at resolution 1.0 with deterministic seed 0. Standard modularity and the community assignment were retained. Community labels are algorithmic partition identifiers, not biological modules.

**Current-flow, curvature, elastic-network, and perturbation-response descriptors.** Parallel directed contact records were collapsed to one undirected edge. Graph-theory calculations used the largest connected component. For edge $\{u, v\}$, conductance and length were

$$c_{uv} = \frac{\exp(-s_{uv})}{\max(d_{uv}, 10^{-9})}; \qquad \ell_{uv} = \frac{1}{c_{uv}},$$

where $s_{uv}$ is contact-geometry surprisal and $d_{uv}$ is residue distance. Normalized node and edge current-flow betweenness used the full weighted Laplacian system over all source-target pairs. Effective resistance was

$$R_{ij} = (\boldsymbol{e}_i - \boldsymbol{e}_j)^{\top} L^{+} (\boldsymbol{e}_i - \boldsymbol{e}_j),$$

where $L^{+}$ is the weighted Laplacian pseudoinverse. The node resistance summary was the mean of $R_{ij}$ over $j \neq i$. Every primary Cα 8 Å graph was connected, so this restriction removed no primary-graph node.

For unweighted edge $\{u, v\}$, the augmented triangle Forman summary was

$$F_{uv} = 4 - k_u - k_v + 3t_{uv},$$

where $k_u$ and $k_v$ are endpoint degrees and $t_{uv}$ is the number of common-neighbor triangles containing the edge. Sensitivity comparisons used the incident mean triangle contribution

$$T_u = \frac{1}{k_u} \sum_{v \sim u} 3\, t_{uv},$$

rather than total curvature, thereby avoiding duplication of the degree term.

The anisotropic network model (ANM), a harmonic elastic-network approximation of directional residue fluctuations, used finite Cα coordinates, a 14 Å connection cutoff, uniform spring constant $\gamma = 1$, and $k_{\mathrm{B}}T = 1$. For connected residues $i$ and $j$ with displacement vector $\boldsymbol{r}_{ij,\mathrm{ANM}}$,

$$H_{ij,\mathrm{ANM}} = -\gamma \frac{\boldsymbol{r}_{ij,\mathrm{ANM}} \boldsymbol{r}_{ij,\mathrm{ANM}}^{\top}}{\|\boldsymbol{r}_{ij,\mathrm{ANM}}\|^2}; \qquad H_{ii,\mathrm{ANM}} = -\sum_{j \neq i} H_{ij,\mathrm{ANM}}.$$

Modes with eigenvalues at most $10^{-8}\max(\max|\lambda|, 1)$ were excluded when forming $H^{+}$. The mean-square fluctuation (MSF), the modeled positional fluctuation magnitude for one residue, was

$$\mathrm{MSF}_i = k_{\mathrm{B}}T\, \mathrm{tr}((H^{+})_{ii}).$$

The perturbation-response matrix was

$$P_{ij} = \frac{1}{3} \|(H^{+})_{ij}\|_F^2,$$

followed by row normalization to $P_{ii}$, with denominator 1 when $P_{ii}$ was nonpositive. Perturbation-response scanning (PRS) summarizes the modeled response pattern from localized harmonic perturbations. PRS effectiveness was the off-diagonal row mean, and PRS sensitivity was the off-diagonal column mean. These are harmonic-network proxies, not measurements of dynamics, kinetics, or causal allostery.

**Contact-definition and graph-domain sensitivities.** Each of the six exact-BW-induced graph definitions was calculated independently for every structure and compared with the primary Cα 8 Å graph. Agreement measures were edge-set Jaccard similarity, Spearman correlation of current-flow betweenness on shared exact-BW nodes, Spearman correlation of $T_u$, cosine similarity of raw 78-dimensional fingerprints, adjusted Rand index for Louvain assignments on shared nodes, and largest-connected-component fraction. For vectors $\boldsymbol{x}$ and $\boldsymbol{y}$, cosine similarity is $\boldsymbol{x}^\mathsf{T}\boldsymbol{y}/(\|\boldsymbol{x}\|_2\|\boldsymbol{y}\|_2)$. The adjusted Rand index is the chance-corrected agreement between two graph partitions. A rank correlation required at least five finite shared values and nonzero variance. Figure S3 reports the median and the 5th and 95th percentiles across 86 structures.

For domain sensitivity, the exact-BW-induced graph was compared with the corresponding whole expected-accession receptor core. Current-flow, triangle Forman, ANM MSF, and PRS-effectiveness correlations and Louvain adjusted Rand indices used exact-BW nodes shared by both domains. Fingerprint cosine similarity compared the complete domain-specific vectors. Figure S4 reports all 86 comparisons. The whole-core calculation is a domain-boundary sensitivity, not a second biological cohort. Graph size is summarized in Figure S2.

**Semantic contact transport.** Four settings retained family or universal bins and burden or binary mass separately. Family bins combined GPCR contact-pair class, segment pair, and receptor-contact context. Universal bins combined generic contact class, structural-segment pair, and intra-chain or interchain context. For contact-row count $n_k$ and $K_{\text{occ}}$ occupied bins,

$$p_k^{(\text{burden})} = \frac{n_k}{\sum_j n_j}; \qquad p_k^{(\text{binary})} = \frac{1}{K_{\text{occ}}}.$$

Conserved mass was $\sum_k \min(p_k, q_k)$; the residual source and destination masses were recorded as loss and gain. Semantic cost $C_{kl}$ was 0 for identical bins and 1 for a namespace mismatch. Otherwise, 0.45 was added for a contact-class mismatch, $0.35[1 - J(S_k, S_l)]$ for segment-token disagreement, and 0.20 for a context mismatch, with total cost capped at 1.5. Token-set Jaccard similarity was $J(S_k, S_l) = |S_k \cap S_l|/|S_k \cup S_l|$. Transport distance was

$$D(p,q) = \min_{\pi \geq 0} \sum_{k,l} \pi_{kl}\, C_{kl},$$

subject to $\sum_l \pi_{kl} = p_k$ and $\sum_k \pi_{kl} = q_l$. The finite minimum-cost-flow problem was solved by successive shortest augmentation without entropic regularization. The four modes remained separate for all 3,655 pairs, yielding 14,620 records. This is a semantic contact-distribution comparison, not physical transport or a signaling trajectory.

**StrucFlow workflow diagnostic.** StrucFlow is a study-introduced graph-perturbation diagnostic that combines geometry, energetic-frustration, water-mediation, dynamic-correlation, and recurrence evidence in a weighted receptor contact network. It is an internal workflow diagnostic, not a physical simulation. StrucFlow used the whole expected-accession receptor-core Cα 8 Å contact ontology for each of 84 independent maps, not the narrower exact-BW-induced graph. Each possible edge carried five sparse channels: geometry $G_{ij}$, signed energetic frustration $E_{ij}$, water mediation $H_{ij,\mathrm{hydr}}$, absolute dynamic correlation $D_{ij}$, and recurrence or persistence $P_{ij,\mathrm{pers}}$. For finite residue distance $d_{uv}$, the geometry channel was

$$G_{ij} = \begin{cases} 1, & d_{uv} \le 8, \\ \frac{1}{2}\left[1 + \cos\left(\frac{\pi(d_{uv} - 8)}{7}\right)\right], & 8 < d_{uv} < 15, \\ 0.25 q_{ij}, & d_{uv} \ge 15, \end{cases}$$

where $q_{ij}$ is the explicit direct-contact indicator. Thus, $G_{ij} = 0$ at or beyond 15 Å when $q_{ij} = 0$. If distance was missing, $G_{ij}$ was the maximum of $q_{ij}$ and any supplied contact-strength value. $E_{ij}$ used the first available value in the order frustration index, geometric-frustration index (GFI) contact score, geometric-frustration score, and essential score divided by 10, with clipping to $[-1,1]$. If those fields were absent, salt, hydrogen-bond, aromatic, pi, or cation contact text gave $E_{ij} = -0.5$; clash or frustration text gave $E_{ij} = +0.5$; otherwise $E_{ij} = 0$. $H_{ij,\mathrm{hydr}}$ was 1 for a mediated contact or water annotation and 0 otherwise. $D_{ij}$ was the clipped absolute value of the first available ANM covariance, ANM correlation, or dynamic correlation. $P_{ij,\mathrm{pers}}$ was the clipped first available cohort recurrence, recurrence fraction, or contact recurrence. Absent $D_{ij}$ and $P_{ij,\mathrm{pers}}$ values were 0. Composite edge weight was

$$A_{ij} = 0.35 G_{ij} + 0.20 I(E_{ij} \ne 0)\left[1 - \min(|E_{ij}|, 1)\right] + 0.15 H_{ij,\mathrm{hydr}} + 0.15 D_{ij} + 0.15 P_{ij,\mathrm{pers}}.$$

Weights below 0.05 were removed. With weighted degree matrix $D_A$, the unnormalized Laplacian was $L = D_A - A$, and algebraic connectivity was its second-smallest eigenvalue. The dimensionless score was

$$S_{\mathrm{SF}} = S_{\mathrm{strain}} + 0.8 S_{\mathrm{pack}} + 0.05 S_{\mathrm{edge}} + 0.2 S_{\mathrm{frust}} + 0.1 S_{\mathrm{hydr}} + 0.2 S_{\mathrm{pers}}.$$

Using the initial contact tensor as reference,

$$S_{\mathrm{strain}} = \frac{1}{24} \sum_{q=1}^{12} \left[\frac{\lambda_q - \lambda_{q0}}{\max(\lambda_{q0}, 10^{-6})}\right]^2,$$

$$S_{\mathrm{pack}} = \frac{1}{2} \mathrm{mean}_i \left[\frac{k_{i,\mathrm{SF}} - k_{i0}}{\max(k_{i0}, 1)}\right]^2,$$

$$S_{\mathrm{edge}} = \mathrm{mean}_{ij} A_{ij}\left[1 - \mathrm{clip}(A_{ij}, 0, 1)\right],$$

$$S_{\mathrm{frust}} = \mathrm{mean}_{ij} G_{ij} |E_{ij}|; \qquad S_{\mathrm{hydr}} = \mathrm{mean}_{ij} H_{ij,\mathrm{hydr}} (1 - G_{ij})^2; \qquad S_{\mathrm{pers}} = \mathrm{mean}_{ij} (G_{ij} - P_{ij,\mathrm{pers}})^2.$$

The means followed the defined nonzero-channel edge sets in the parameter registry. Event mode used 100 stochastic proposals, seed 7, and at most 3,000 candidates per proposal. One candidate edge was selected uniformly after any candidate subsampling. Available moves increased or decreased its geometry weight by 0.1; additionally, a weight above 0.2 could be set to 0 and a weight below 0.2 could be formed at $\max(0.1, w + 0.1)$. A move that reduced an edge between residue-index positions separated by at most two below weight 0.3 was rejected by the topology constraint. At inverse temperature 1, the Metropolis acceptance probability was

$$\Pr(\text{accept}) = \begin{cases} 1, & \Delta S \leq 0, \\ \exp(-\Delta S), & 0 < \Delta S < 700, \\ 0, & \Delta S \geq 700. \end{cases}$$

Incremental score differences used first-order unnormalized-Laplacian eigenvalue perturbation and exact local terms; exact scores were recalculated at proposal 1, every 25 proposals, and at the last proposal. Continuous-weight mode used 50 iterations, a uniform random batch of 8 candidate weights, central finite differences with step 0.025, learning rate 0.01, independent Gaussian noise scale 0.001, clipping to the interval from 0 to 1, the same topology constraint, and exact recalculation at iteration 1, every 10 iterations, and at the last iteration. For candidate edge $e$, let $\boldsymbol{w}_{t,e\leftarrow x}$ denote the iteration-$t$ weight vector with the weight of edge $e$ replaced by $x$. Then

$$w_e^+ = \min(1, w_{e,t} + 0.025); \qquad w_e^- = \max(0, w_{e,t} - 0.025); \qquad g_{e,t} = \frac{S_{\mathrm{SF}}(\boldsymbol{w}_{t,e\leftarrow w_e^+}) - S_{\mathrm{SF}}(\boldsymbol{w}_{t,e\leftarrow w_e^-})}{w_e^+ - w_e^-},$$

$$w_{e,t+1} = \mathrm{clip}(w_{e,t} - 0.01 g_{e,t} + 0.001 \xi_{e,t}, 0, 1); \qquad \xi_{e,t} \sim \mathcal{N}(0,1).$$

An auxiliary reconstruction used 10 ANM reconciliation modes with damping 0.7. Figure S7A and S7B report final-minus-initial graph score and algebraic connectivity for 168 validated map-mode records. Event index is not physical time, edge updates are not molecular dynamics, and the score is not a calibrated free energy.

**Contact-propagation-network workflow diagnostic.** The study-introduced contact-propagation network (CPN) is a three-state probabilistic factor-graph diagnostic that perturbs certified selected-ligand contact seeds and measures the resulting change in approximate node-state beliefs. A factor graph represents node-specific state scores as unary factors and edge-specific compatibility scores as pair factors. The CPN used the whole expected-accession receptor-core Cα 8 Å factor graph. Selected-ligand seeds were exact-BW residues supported by role-matched and target-matched PLIP direct-contact records. No missing seed was imputed. Seventy-nine of 86 structures had certified seed sets and seven were not applicable; Figure S7C uses the 77 seeded independent-map representatives.

Each residue had constrained $C_{\text{CPN}}$, adaptive $A_{\text{CPN}}$, and strained $S_{\text{CPN}}$ states. Composite weighted degree was calculated before the 0.05 StrucFlow sparsity threshold. Let $J_{E,i}$ and $J_{P,i}$ denote the nonzero energetic-evidence and persistence-evidence neighbor sets, respectively. The CPN node aggregates were

$$d_{i,\text{CPN}} = \frac{\text{degree}_i}{\max_j \text{degree}_j}; \qquad f_{i,\text{CPN}} = \frac{\sum_{j \in J_{E,i}} G_{ij} E_{ij}}{\sum_{j \in J_{E,i}} G_{ij}}; \qquad p_{i,\text{CPN}} = \frac{\sum_{j \in J_{P,i}} G_{ij} P_{ij,\text{pers}}}{\sum_{j \in J_{P,i}} G_{ij}}; \qquad h_{i,\text{CPN}} = \sum_j H_{ij,\text{hydr}}.$$

For each geometry-weighted row mean, only nonzero evidence entries with positive geometry weight contributed. If no positive geometry weight was available for a nonempty evidence row, the unweighted row mean was used, and an empty row returned 0. The certified-seed indicator was $z_{i,\text{CPN}}$. Unary scores were

$$U_i(C_{\text{CPN}}) = 1 + 1.25 d_{i,\text{CPN}} + 0.75 p_{i,\text{CPN}} - 0.5 \max(f_{i,\text{CPN}}, 0),$$

$$U_i(A_{\text{CPN}}) = 1 + 0.4 h_{i,\text{CPN}} + 0.5(1 - p_{i,\text{CPN}}) + 0.4 |f_{i,\text{CPN}}|,$$

$$U_i(S_{\text{CPN}}) = 1 + 1.2 \max(f_{i,\text{CPN}}, 0) + 0.5 |f_{i,\text{CPN}}| + 2 z_{i,\text{CPN}}.$$

Unary potentials were floored at 0.05, $\psi_i(x_{\text{CPN}}) = \max[U_i(x_{\text{CPN}}), 0.05]$. Pair potentials began at 1. For edge strength $x_{\text{edge}}$ from the unthresholded composite contact weight, define $f^+ = \max(f_{\text{edge}}, 0)$ and $f^- = \max(-f_{\text{edge}}, 0)$, with hydration $h_{\text{edge}}$ and persistence $p_{\text{edge}}$. The pair potentials were

$$\psi_{\text{CC}} = 1 + x_{\text{edge}}(0.6 + 0.8 p_{\text{edge}} + 0.5 f^-), \qquad \psi_{\text{AA}} = 1 + x_{\text{edge}}(0.25 + 0.4 h_{\text{edge}}).$$

$$\psi_{\text{SS}} = 1 + x_{\text{edge}}(0.25 + 0.8 f^+), \qquad \psi_{\text{CA}} = \psi_{\text{AC}} = 1 + 0.25 x_{\text{edge}}.$$

$$\psi_{\text{AS}} = \psi_{\text{SA}} = 1 + 0.25 x_{\text{edge}}(1 + f^+ + h_{\text{edge}}).$$

$$\psi_{\text{CS}} = \psi_{\text{SC}} = \max(0.35, 1 - 0.35 x_{\text{edge}}).$$

Entries were floored at 0.05 and normalized by their matrix mean. At most the 20,000 strongest pair factors were retained.

Loopy sum-product belief propagation, an iterative approximation to marginal-state inference on a graph containing cycles, used

$$m_{i \to j}(x_{\text{CPN},j}) \propto \sum_{x_{\text{CPN},i}} \psi_i(x_{\text{CPN},i}) \psi_{ij}(x_{\text{CPN},i}, x_{\text{CPN},j}) \prod_{\substack{k \in N_{\text{CPN}}(i) \\ k \neq j}} m_{k \to i}(x_{\text{CPN},i}),$$

with uniform initial messages, damping 0.5, at most 150 iterations, and convergence tolerance $10^{-6}$. A node belief is the resulting normalized approximate probability distribution over that node's three states. Baseline beliefs used $z_{i,\text{CPN}} = 0$; perturbed beliefs added the strain score of 2 at certified seed sites. The displayed response was

$$\Delta_{\text{max}} = \max_{i,x_{\text{CPN}}} \left| b_{i,\text{seed}}(x_{\text{CPN}}) - b_{i,\text{base}}(x_{\text{CPN}}) \right|.$$

One hundred seeded belief samples were an ancillary diagnostic and did not enter the displayed shift. An explicit no-seed control returned zero shift. These values are belief-space workflow diagnostics, not physical conformations, molecular dynamics, signaling efficacy, or causal propagation.

**Target and coordinate evidence completeness.** Completeness was evaluated at the independent-map level and kept separate from biological prevalence. Coordinate evidence was the role-aware selected-ligand union: an exact-BW receptor residue was included when a nonhydrogen atom lay within 4.5 Å of a curated orthosteric or allosteric target atom. Target-level evidence came from independently curated selected-ligand records. For each map, the analysis counted selected targets and PLIP-resolved targets and identified whether every defined target was PLIP-resolved or whether the direct set was partial. Maps without a defined selected target were retained with zero target records, and all-target completeness was not applicable rather than false. Coordinate and PLIP evidence were not substituted for one another. Figure S8 measures selected-target annotation and coordinate-evidence completeness, not ligand absence or biological prevalence.

The public reproducibility supplement is a scientific specification for the methods introduced in this study. It contains ordered pseudocode in human-readable and JavaScript Object Notation (JSON) forms, complete parameters, equations and symbol definitions, authoritative table schemas, water-set definitions and membership, a public data dictionary, checksum-linked derivation and validation records, and a claim-to-source map. Separate non-image source tables contain the data underlying each figure; figure-rendering code and figure-specific pseudocode are excluded. Stable artifact identifiers connect inputs, transformations, outputs, validation criteria, record keys, and reported rounding. This design follows Journal of Chemical Information and Modeling (JCIM) and joint American Chemical Society (ACS) guidance on method, data, software, and workflow transparency, with the corresponding sources cited in the Article.

#### Supplementary Results and Interpretation Boundaries

##### S10. State result is a literature-directed positive control

Six of 16 state outcomes survive within-family correction (Table S3). Their directions are consistent with inactive, active, and transducer-bound structural differences described and cited in the Article. Because structures that established these conformational distinctions are represented, this is a literature-directed workflow positive control, not independent replication or confirmation of a specific activation mechanism. The shared-node-restricted full endpoint-recomputation leave-one-out analysis comprised 378 outcome-specific fits, and every retained coefficient preserved its sign in 100% of deletions.

##### S11. Site-relative spatial atlas

For the selected-ligand union, pooled static wiring is 13.39% direct, 32.53% adjacent, and 54.08% connected-distal. The primary shared-node comparison contains 239,736 changed contacts: 15.28% direct, 40.13% adjacent, and 44.59% connected-distal (Table S4). The full-edge sensitivity contains 281,166 changed contacts and gives 13.19%, 36.31%, and 50.49%, respectively. Of those 281,166 full-edge changed-contact records, 41,430, or 14.74%, have at least one endpoint outside the exact-BW node set shared by the pair. These percentages locate graph edges relative to a resolved site. They are not a physical signal trajectory.

Structural-water sites place 64.61% of static wiring and 65.89% of shared-node changed contacts in connected-distal graph space. Ligand-contact-water sites place 82.67% and 86.01%, respectively, in connected-distal space. Sterol, generic-lipid, allosteric-modulator, and Gα sites also have majority connected-distal wiring. These patterns identify distal regions represented in deposited models but do not show that the component caused the contact distribution.

##### S12. Percentage-scale associations remain exploratory after global correction

None of the 32 component associations survives global correction (Tables S5 and S6; Figure S5). Percentage tables remain useful because they show scale, direction, coverage, and uncertainty. The peptide-versus-small-molecule RMS coefficient is negative in every structure and publication leave-one-out refit. Eighteen matched pairs have a mean difference of $-0.01889$ Å, and three of five discordant publications agree with the adjusted direction. The global $q$ value is 0.3165. For structural-water-positive maps, the minimum-$p$ result under the shared-node primary estimand is 7.67% higher shared-contact RMS distance, with a 95% interval from $-0.21\%$ to $+15.55\%$ and $q = 0.4511$. Only seven maps are exposed, with four matched pairs and one discordant publication. Sterol is the only nonligand component with substantial prevalence and six discordant publications; its adjusted interval crosses zero. These are graded evidence statements, not discovery claims.

**S13. Deposited-water branches, recurrence, and dominant-map sensitivity**

At default settings, $P$ contains 137 waters. The structural branch contains 61 core waters, 52 designated-receptor bridges, 44 core waters with an exact-BW contact, and 38 core waters bridging at least two exact-BW residues (Table S7; Figure S6). $B_{\mathrm{any}}$ independently contains 85 proximal waters with any exact-BW contact. The ligand-contact branch $L$ contains 10 waters. Three of those 10 also belong to the 38-water $C_{\mathrm{B2}}$ set, and seven lie outside the structural core. Thus, the structural and ligand-contact branches answer different deposited-coordinate questions.

DOR 4N6H supplies 98/137 receptor-proximal waters, 71.5%, and 39/61 core waters, 63.9%. Without it, 39 proximal waters, 29 any-BW-contact waters, 22 core waters, and 17 core exact-BW bridges remain. The ligand branch retains 8 waters in 3 maps, 3 of which overlap $C_{\mathrm{B2}}$ and 5 of which lie outside $C$.

At a 1.0 Å radius, the complete-linkage and single-linkage definitions each yield six recurrent coordinate clusters, and each yields three after removing 4N6H. Among 21 unordered pairs of seven $C_{\mathrm{B2}}$-positive maps, 6/21 have at least one one-to-one coordinate match; after removing 4N6H, 2/15 do. Twenty-five exact-BW bridge topologies recur and 15 persist without 4N6H. The 3×32/3×35 topology occurs in 3/84 processed maps, 3.57%, and 3/7 $C_{\mathrm{B2}}$-positive maps, 42.86%; it spans DOR, KOR, and NOP and persists in KOR and NOP after 4N6H exclusion. The first percentage uses the full processed-map denominator, whereas the second conditions on maps capable of contributing a $C_{\mathrm{B2}}$ topology. Coordinate recurrent clusters occur only among X-ray diffraction structures. Topological recurrence also contains one mixed-method DOR pair, X-ray diffraction 4N6H and cryogenic electron microscopy (cryo-EM) 9YDP, at 3×54/5×64. These are static coordinate and topology observations, not solution hydration or dynamic water wires.

**S14. Sparse systems are structure-specific recapitulations**

Short-chain PI(4,5)$P_2$ (PDB ligand PIO) and β-arrestin 1 occur together only in the engineered KOR/vasopressin V2 receptor (V2R)-tail 9ZZO preparation. The receptor construct, PIO, β-arrestin 1, Fab30, Nb32, and other stabilizing elements cannot be separated statistically. PIO contacts the full receptor within 5.0 Å but does not contact an exact-BW core residue within 5.0 Å. Sodium appears in three maps, and selected-ligand/water/receptor contacts appear in four. Deposited absence often means unresolved or not modeled. These records support exact structure queries and contextual comparison, not a cohort-general $PIP_2$, β-arrestin, sodium, or hydration effect (Table S8). StrucFlow and CPN diagnostics and structure-target evidence completeness are shown in Figures S7 and S8 as workflow quality control, not mechanistic evidence.

#### Supplementary Tables

##### Table S1. Cohort and curated-role counts

| Category | Count or identifiers |
|---|---|
| Candidate entries | 99 |
| Included human opioid structures | 86 |
| Independent experimental maps | 84 |
| MOR/KOR/DOR/NOP, full | 35/28/18/5 |
| MOR/KOR/DOR/NOP, primary | 33/28/18/5 |
| Primary states: active-partner/inactive/unclear/active-like | 49/16/15/4 |
| Primary acquisition: electron microscopy/X-ray diffraction | 72/12 |
| Curated binding modes: orthosteric/orthosteric plus allosteric/allosteric only/none | 72/9/1/2 |
| Curated allosteric-contact maps | 8K9L, 8Y71, 8Y72, 8Y73, 9L60, 9PU5, 9XC6, 9XDQ, 9XDR, 9XF4 |
| 9L60 selected ligand | Dynorphin A(1-13), chain P; PLIP 14 exact-BW contacts; coordinate dynorphin 25, MPAM-15 10, union 35 |
| 9ZZO PIO identity | Short-chain diC$_8$-PI(4,5)P$_2$; analysis category PIP$_2$-like lipid |
| Shared-map relationships | 10TL with 9PU5; 10TM with 9PUD |

##### Table S2. Primary-map activity and component prevalence

| Feature | n/N | Percent | Wilson 95% interval | Reporting level |
|---|---|---|---|---|
| Unspecified agonist | 50/84 | 59.5% | 48.8% to 69.4% | Descriptive category |
| Antagonist | 15/84 | 17.9% | 11.1% to 27.4% | Descriptive category |
| Inverse agonist | 4/84 | 4.8% | 1.9% to 11.6% | Sparse |
| Biased agonist | 1/84 | 1.2% | 0.2% to 6.4% | Singleton |
| Partial agonist | 1/84 | 1.2% | 0.2% to 6.4% | Singleton |
| Full agonist | 1/84 | 1.2% | 0.2% to 6.4% | Singleton |
| Unresolved activity | 9/84 | 10.7% | 5.7% to 19.1% | Preserved unknown |
| No orthosteric ligand | 3/84 | 3.6% | 1.2% to 10.0% | Not applicable |
| Receptor-contacting sterol | 26/84 | 31.0% | 22.1% to 41.5% | Group descriptive |
| Receptor-contacting generic lipid | 11/84 | 13.1% | 7.5% to 21.9% | Limited overlap |
| Deposited receptor-proximal water | 9/84 | 10.7% | 5.7% to 19.1% | Coverage-limited |
| Two-protein-residue structural water | 7/84 | 8.3% | 4.1% to 16.2% | Coverage-limited |
| Resolved G-protein assembly | 66/84 | 78.6% | 68.7% to 86.0% | Confounded context |
| Direct Gβ contact within Gα-positive maps | 11/66 | 16.7% | 9.6% to 27.4% | Within-Gα only |
| Orthosteric peptide in resolved-modality subset | 19/76 | 25.0% | 16.6% to 35.8% | Group descriptive |

| Feature | n/N | Percent | Wilson 95% interval | Reporting level |
|---|---|---|---|---|
| Orthosteric plus allosteric in resolved-binding subset | 9/81 | 11.1% | 6.0% to 19.8% | Publication-limited |
| Receptor-contacting allosteric modulator | 10/84 | 11.9% | 6.6% to 20.5% | Group descriptive |
| $PIP_2$-like lipid analysis category | 1/84 | 1.2% | 0.2% to 6.4% | Structure-specific |
| β-arrestin | 1/84 | 1.2% | 0.2% to 6.4% | Structure-specific |
| Receptor-contacting sodium | 3/84 | 3.6% | 1.2% to 10.0% | Structure-specific |
| Selected-ligand/water/receptor contact | 4/84 | 4.8% | 1.9% to 11.6% | Structure-specific |

#### Table S3. Complete 16-outcome state positive-control family

| Outcome | $n$ | Adjusted % of active-partner mean (95% CI) | $p$ | $q$ | Reported result |
|---|---|---|---|---|---|
| Exact-BW contact-pair count | 65 | -1.67% (-2.97%, -0.38%) | 0.0122 | 0.0326 | Yes; full endpoint leave-one-out sign 100% |
| Endpoint-mean contact Jaccard similarity | 65 | -2.49% (-3.74%, -1.24%) | 0.0001906 | 0.00152 | Yes; full endpoint leave-one-out sign 100% |
| Endpoint-mean rewired-contact count | 65 | +34.84% (+16.78%, +52.90%) | 0.0002918 | 0.00156 | Yes; full endpoint leave-one-out sign 100% |
| Endpoint-mean shared-contact distance RMS deviation | 65 | +15.15% (+4.84%, +25.45%) | 0.00471 | 0.0151 | Yes; full endpoint leave-one-out sign 100% |
| Direct static-edge count | 59 | -13.84% (-39.67%, +11.99%) | 0.2869 | 0.4779 | No |
| Adjacent static-edge count | 59 | +2.62% (-13.68%, +18.92%) | 0.7482 | 0.8551 | No |
| Connected-distal static-edge count | 59 | -0.07% (-13.66%, +13.52%) | 0.9916 | 0.9916 | No |
| Direct static-edge fraction | 59 | -12.63% (-38.35%, +13.08%) | 0.3286 | 0.4779 | No |
| Adjacent static-edge fraction | 59 | +3.92% (-11.78%, +19.61%) | 0.6182 | 0.7608 | No |
| Connected-distal static-edge fraction | 59 | +0.89% (-13.38%, +15.16%) | 0.9007 | 0.9607 | No |
| Endpoint-mean changed-direct count | 59 | +25.05% (+9.87%, +40.24%) | 0.00171 | 0.00686 | Yes; full endpoint leave-one-out sign 100% |

| Outcome | $n$ | Adjusted % of active-partner mean (95% CI) | $p$ | $q$ | Reported result |
|---|---|---|---|---|---|
| Endpoint-mean changed-adjacent count | 59 | +34.78% (+18.64%, +50.92%) | 0.0000722 | 0.00116 | Yes; full endpoint leave-one-out sign 100% |
| Endpoint-mean changed connected-distal count | 59 | +40.31% (+5.36%, +75.26%) | 0.0247 | 0.0564 | No |
| Endpoint-mean changed-direct fraction | 59 | -10.23% (-27.18%, +6.71%) | 0.2310 | 0.4619 | No |
| Endpoint-mean changed-adjacent fraction | 59 | -2.99% (-12.68%, +6.70%) | 0.5380 | 0.7173 | No |
| Endpoint-mean changed connected-distal fraction | 59 | +7.31% (-6.70%, +21.32%) | 0.2998 | 0.4779 | No |

Disconnected static and changed outcomes were structurally zero under the primary graph and were not tested.

##### Table S4. Site-relative pooled wiring and changed-contact distributions

| Exact-BW site group | Positive maps | Static %, D/A/CD | Eligible pairs | Changed %, D/A/CD | Interpretation |
|---|---|---|---|---|---|
| Selected-ligand union | 77 | 13.39 / 32.53 / 54.08% | 2,926 | 15.28 / 40.13 / 44.59% | 239,736 shared-node-restricted changed contacts; all primary receptor graphs connected |
| Structural water | 7 | 10.73 / 24.66 / 64.61% | 21 | 10.44 / 23.67 / 65.89% | Site location, not water causality |
| Ligand-contact water | 4 | 2.76 / 14.57 / 82.67% | 6 | 1.27 / 12.72 / 86.01% | Structure-specific site geometry |
| Sterol | 26 | 17.25 / 30.82 / 51.93% | 325 | 9.88 / 36.61 / 53.51% | Deposited sterol-contact geometry |
| Generic lipid | 11 | 16.56 / 27.47 / 55.97% | 55 | 14.87 / 36.13 / 49.00% | Deposited lipid-contact geometry |
| Allosteric modulator | 10 | 9.45 / 16.39 / 74.16% | 45 | 6.22 / 22.79 / 70.99% | Site location, not effect |
| Gα interface | 66 | 9.21 / 15.12 / 75.67% | 2,145 | 5.67 / 11.27 / 83.06% | Intracellular interface geometry |

Disconnected is 0% for every listed site group under the Cα 8 Å graph. Pair denominators require both maps to resolve the specified site.

In Table S4, D, A, and CD denote direct, adjacent, and connected-distal, respectively.

##### Table S5. Unadjusted spatial percentage shifts in the 77-map selected-site subset

| Exposed versus reference context | n exposed/ref. | Outcome | Raw difference as % of reference mean | 5,000-bootstrap 95% interval | Interpretation |
|---|---|---|---|---|---|
| Sterol contact versus reference | 23/54 | Changed-direct fraction | +14.92% | +6.24%, +24.11% | Descriptive composition |
| Sterol contact versus reference | 23/54 | Changed connected-distal fraction | -9.20% | -15.22%, -3.14% | Descriptive composition |
| Generic-lipid contact versus reference | 9/68 | Changed-direct fraction | -17.03% | -22.78%, -11.36% | Limited overlap |
| Generic-lipid contact versus reference | 9/68 | Changed connected-distal fraction | +15.75% | +5.77%, +25.45% | Limited overlap |
| Deposited water versus reference | 9/68 | Changed-direct fraction | -13.65% | -21.20%, -5.43% | Coverage-limited |
| Deposited water versus reference | 9/68 | Changed connected-distal fraction | +12.18% | +0.66%, +23.80% | Coverage-limited |
| Structural water versus reference | 7/70 | Changed-direct fraction | -15.99% | -24.00%, -5.84% | Seven-map pattern |
| Structural water versus reference | 7/70 | Changed connected-distal fraction | +18.03% | +5.35%, +29.00% | Seven-map pattern |
| G-protein present versus reference | 60/17 | Changed-direct fraction | +16.42% | +9.48%, +23.96% | Partner/state/construct bundle |
| G-protein present versus reference | 60/17 | Changed connected-distal fraction | -13.85% | -18.57%, -8.91% | Partner/state/construct bundle |
| Peptide versus small molecule | 19/57 | Static direct fraction | +24.28% | +8.12%, +41.95% | Ligand-modality context |
| Peptide versus small molecule | 19/57 | Static connected-distal fraction | -10.17% | -16.97%, -3.84% | Ligand-modality context |
| Orthosteric plus allosteric versus orthosteric | 9/67 | Changed-direct fraction | +20.05% | +9.65%, +32.81% | Publication-limited |

| Exposed versus reference context | n exposed/ref. | Outcome | Raw difference as % of reference mean | 5,000-bootstrap 95% interval | Interpretation |
|---|---|---|---|---|---|
| Orthosteric plus allosteric versus orthosteric | 9/67 | Changed connected -distal fraction | -15.84% | -24.43%, -7.90% | Publication-limited |
| Allosteric contact versus reference | 10/67 | Changed-direct fraction | +17.70% | +7.50%, +30.11% | Physical-contact context |
| Allosteric contact versus reference | 10/67 | Changed connected -distal fraction | -13.56% | -22.02%, -5.02% | Physical-contact context |

These intervals describe unadjusted structure-level composition and do not control confounding, publication clustering, or multiplicity.

##### Table S6. Multivariable-adjusted percentage-scale component associations

| Contrast | Minimum-p outcome among four | Adjusted % of reference mean (95% CI) | p | Global q | Matched pairs/discordant publications | Evidence boundary |
|---|---|---|---|---|---|---|
| Sterol contact | Endpoint-mean shared-contact distance RMS deviation | -7.28% (-17.56%, +3.01%) | 0.163 | 0.6970 | 17/6 | Direction unclear |
| Generic-lipid contact | Endpoint-mean shared-contact distance RMS deviation | -5.76% (-13.39%, +1.87%) | 0.137 | 0.6970 | 5/2 | Limited overlap |
| Deposited receptor-proximal water | Endpoint-mean shared-contact distance RMS deviation | +3.42% (-3.69%, +10.52%) | 0.340 | 0.7251 | 6/1 | Coverage-limited |
| Two-protein-residue structural water | Endpoint-mean shared-contact distance RMS deviation | +7.67% (-0.21%, +15.55%) | 0.0564 | 0.4511 | 4/1 | Exploratory, replication-limited |
| Resolved G-protein assembly | Endpoint-mean shared-contact distance RMS deviation | +1.58% (-14.08%, +17.25%) | 0.841 | 0.9737 | 2/1 | Confounded context |
| Direct Gβ contact within Gα | Exact-BW contact-pair count | +0.31% (-1.22%, +1.84%) | 0.688 | 0.9737 | 9/5 | No association detected |
| Peptide versus small molecule | Endpoint-mean shared-contact distance RMS deviation | -6.18% (-10.82%, -1.54%) | 0.00989 | 0.3165 | 18/5 | Three of five publications align; exploratory |
| Orthosteric plus allosteric versus orthosteric | Endpoint-mean contact Jaccard similarity | +0.47% (-0.24%, +1.19%) | 0.193 | 0.6970 | 9/1 | Publication-limited |

The displayed outcome is selected by minimum nominal p within each contrast's four defined outcomes. All 32 rows, not the eight display rows, define the global multiplicity family. Zero of 32 has global q < 0.05. Jaccard similarity and rewired count are highly redundant and are not independent support.

##### Table S7. Default water estimands, overlap, and recurrence

| Estimand group | Quantity | All maps, waters/maps | Without 4N6H, waters/maps |
|---|---|---|---|
| Root | Receptor-proximal deposited water, $P$ | 137/9 | 39/8 |
| Exact-BW contact | Any proximal exact-BW contact, $B_{\text{any}}$ | 85/8 | 29/7 |
| Structural | Two-protein-residue core, $C$ | 61/7 | 22/6 |
| Structural | At least two designated-receptor partners, $R_2$ | 52/7 | 18/6 |
| Structural plus exact BW | Core plus at least one exact-BW partner, $C_{\text{B1}}$ | 44/7 | 18/6 |
| Structural plus exact BW | Core plus at least two exact-BW partners, $C_{\text{B2}}$ | 38/7 | 17/6 |
| Ligand-contact branch | Selected-ligand/water/receptor, $L$ | 10/4 | 8/3 |
| Branch overlap | $L \cap C_{\text{B2}}$ | 3/2 | 3/2 |
| Ligand-contact branch | $L$ outside $C$ | 7 waters | 5 waters |
| Dominant-map share | 4N6H share of $P/C$ | 71.5%/63.9% | Not applicable |
| PLIP comparison | Raw/unique exact-mapped events | 19/16 | Not applicable |
| PLIP comparison | Unique events overlapping $C$ | 7/16 (43.8%) | Not applicable |
| Coordinate recurrence | Recurrent clusters at 1.0 Å | 6 | 3 |
| Coordinate recurrence | Map pairs with at least one one-to-one match | 6/21 | 2/15 |
| Topological recurrence | Recurrent exact-BW pairs | 25 | 15 |
| Topological recurrence | 3×32/3×35 occurrence | 3/84 maps, 3.57%; 3/7 $C_{\text{B2}}$-positive maps, 42.86% | 2 maps |

##### Table S8. Claim-language boundaries

| Supported statement | Unsupported expansion |
|---|---|
| State directions are consistent with inactive, active, and transducer-bound structural differences described and cited in the Article | Independent replication, rediscovery of activation, or validation of every microswitch |
| Static wiring or changed-contact percentage relative to a resolved site | Dynamic signal propagation or information-flow direction |
| Percentage magnitude with interval and coverage label | General component effect when global q is at least 0.05 |
| Multiplicity-qualified exploratory hypothesis | Discovery-level mechanism |

| Supported statement | Unsupported expansion |
|---|---|
| Receptor-contacting sterol, lipid, or water in deposited coordinates | Component experimentally added or biologically absent in reference maps |
| Resolved heterotrimeric G-protein context | Isolated Gβ/Gγ effect |
| Within-Gα direct receptor/Gβ contrast | Gβ assembly effect |
| Static deposited-water coordinate and topology atlas | Solution hydration, residence time, energetics, or causal water wire |
| 9ZZO records an engineered KOR/V2R-tail/β-arrestin 1 preparation with short-chain PI(4,5)$P_2$ | Cohort-general $PIP_2$ or β-arrestin effect |
| Cohort-wide quantitative extension | Priority claim or first discovery of allostery, distal communication, or water networks |

### Supplementary Figures

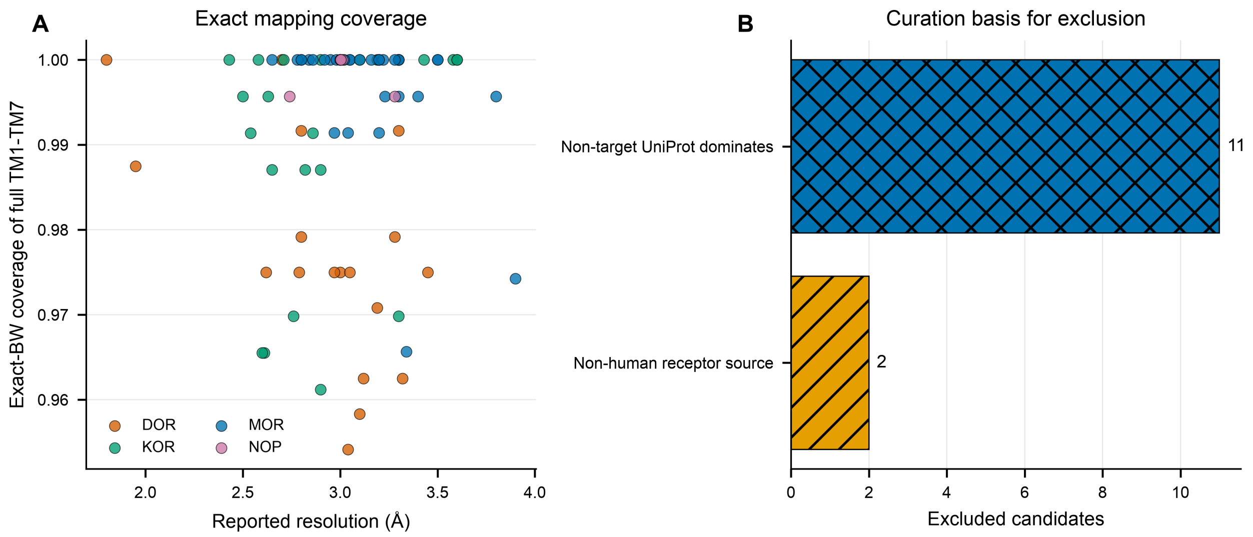


**Figure S1. Cohort curation and exact-mapping quality control.** (A) Exact-BW coverage versus reported resolution for 86 included structures. Exact-BW coverage is the resolved exact-BW Cα count divided by the full GPCRdb TM1 through TM7 exact-BW position count for that receptor. This plot quantity is distinct from the SIFTS mapping-coverage covariate used in the regression and matching analyses. (B) Curation basis for 13 exclusions: 11 non-target chimeric or non-opioid receptor polymers and 2 non-human receptor structures. No excluded candidate contributes to the analysis cohort.

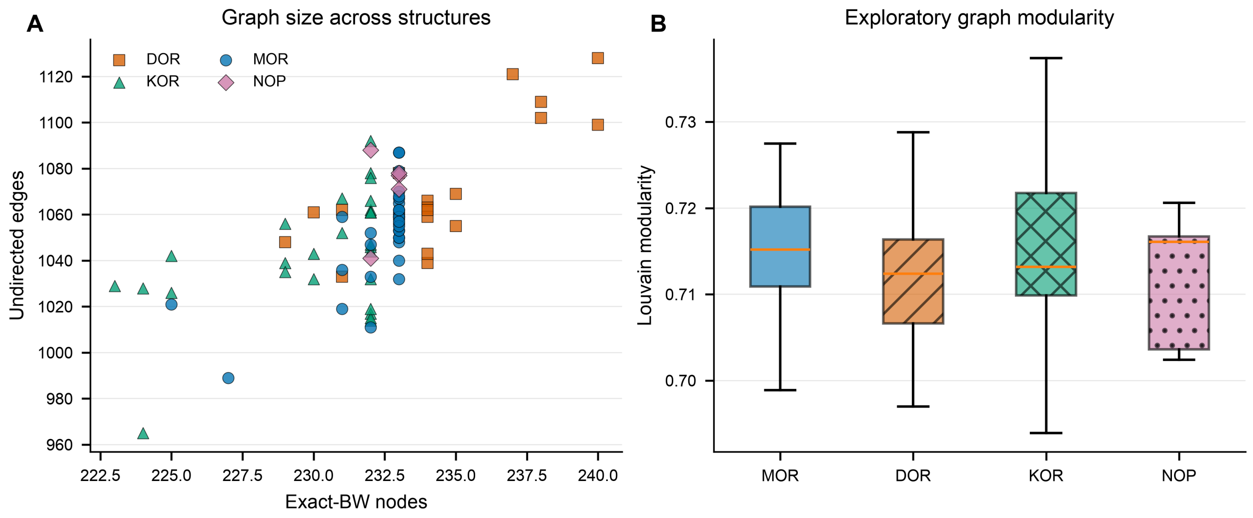


**Figure S2. Exact-BW graph size and exploratory modularity.** (A) Node and edge counts for all 86 exact-BW Cα 8 Å graphs. (B) Louvain modularity by subtype. Louvain communities are algorithmic partitions and are not validated biological modules.

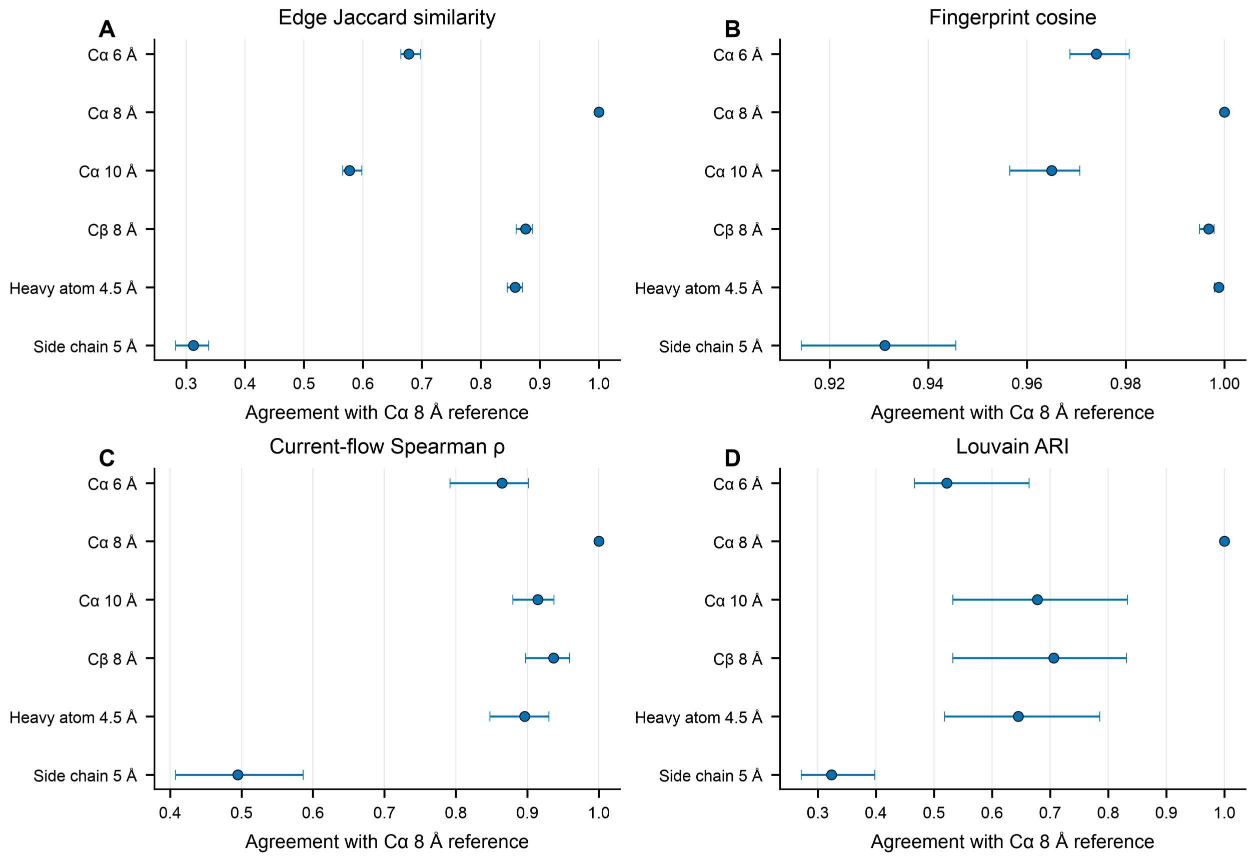


**Figure S3. Contact-definition sensitivity.** Median and 5th-95th percentile agreement with the exact-BW Cα 8 Å reference across 86 structures for 6 contact definitions. Cα 8 Å is the identity reference. These metrics quantify algorithmic sensitivity and do not validate a biological contact mechanism.

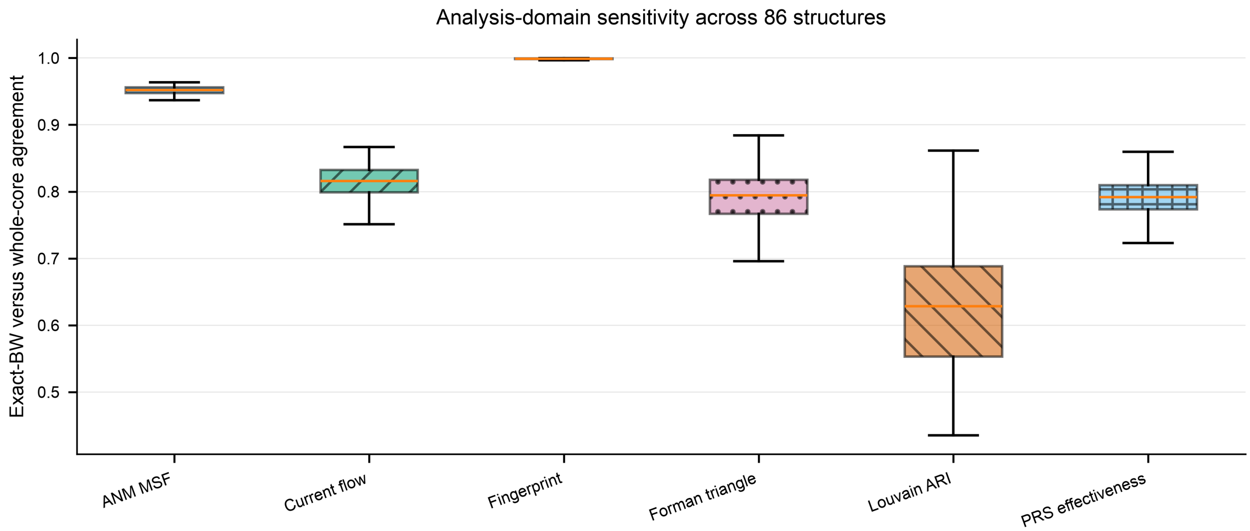


**Figure S4. Analysis-domain sensitivity.** Agreement between exact-BW induced graphs and whole exact-target receptor cores for 6 graph descriptors across 86 structures. Boxes show median and interquartile range; whiskers extend to 1.5 times the interquartile range. Whole-core results are a domain-boundary sensitivity, not a second cohort.

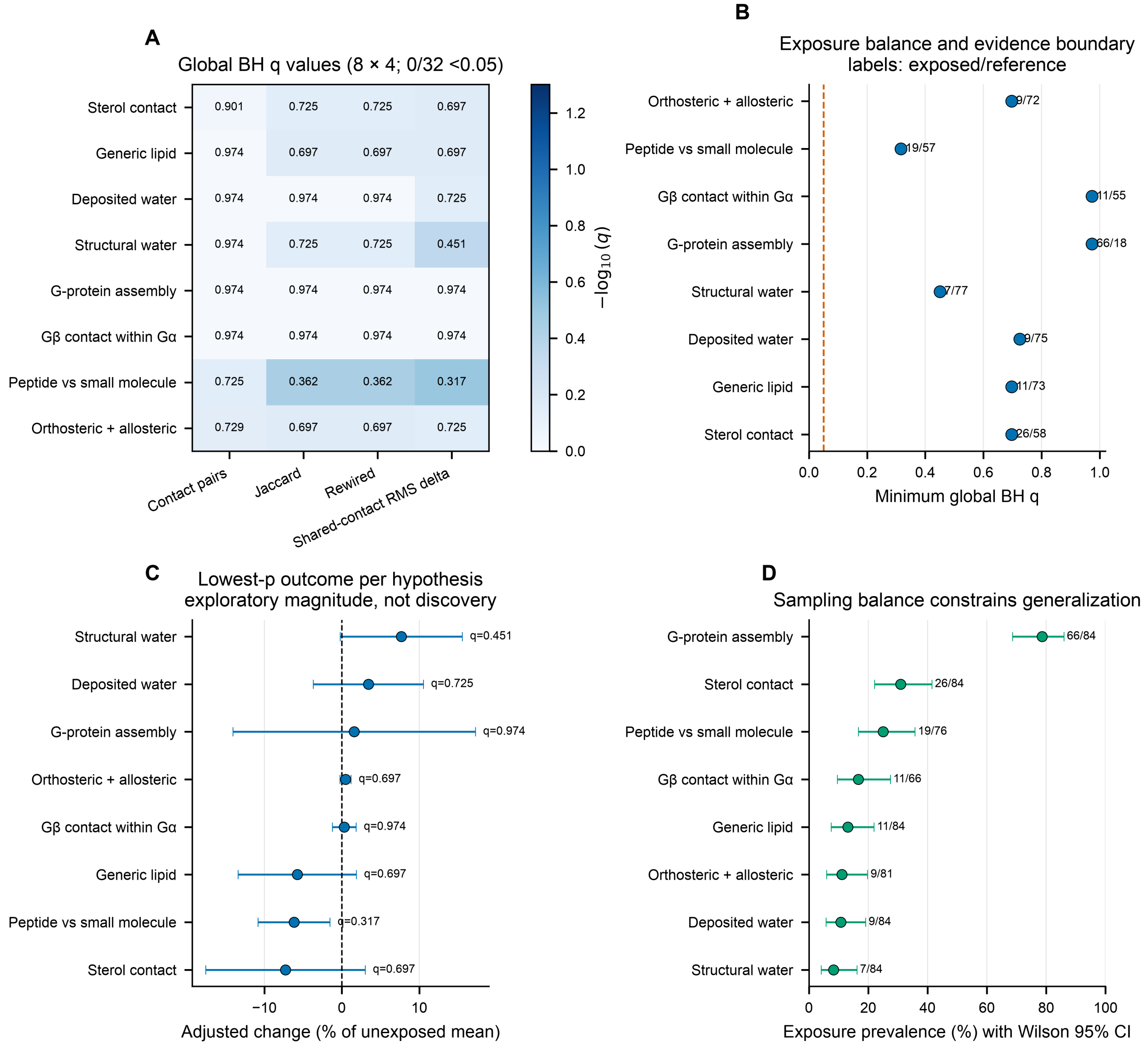


**Figure S5. Complete deduplicated 32-test component association family.** All eight defined hypotheses and four outcomes are displayed with one multivariable-adjusted estimate each and one global Benjamini-Hochberg correction. Zero of 32 tests passes $q < 0.05$. The smallest $q$ is 0.3165 for peptide versus small-molecule shared-contact RMS distance; its magnitude and direction are exploratory only. Descriptive percentages and sensitivity analyses may motivate hypotheses but do not override multiplicity. No hydration, sterol, lipid, $PIP_2$, sodium, G-protein, Gβ, allosteric, efficacy, or other component effect is FDR-supported. Adjusted percentage magnitudes and 95% intervals are shown for each hypothesis's lowest-$p$ outcome: sterol RMS −7.28%, generic-lipid RMS −5.76%, deposited-water RMS +3.42%, structural-water RMS +7.67%, G-protein RMS +1.58%, Gβ exact-contact count +0.31%, peptide-versus-small-molecule RMS −6.18%, and orthosteric-plus-allosteric Jaccard +0.47%. These percentages are conditional descriptive estimates; intervals, overlap limitations, matched-pair direction, leave-one-out stability, and global $q$ values remain in the source tables.

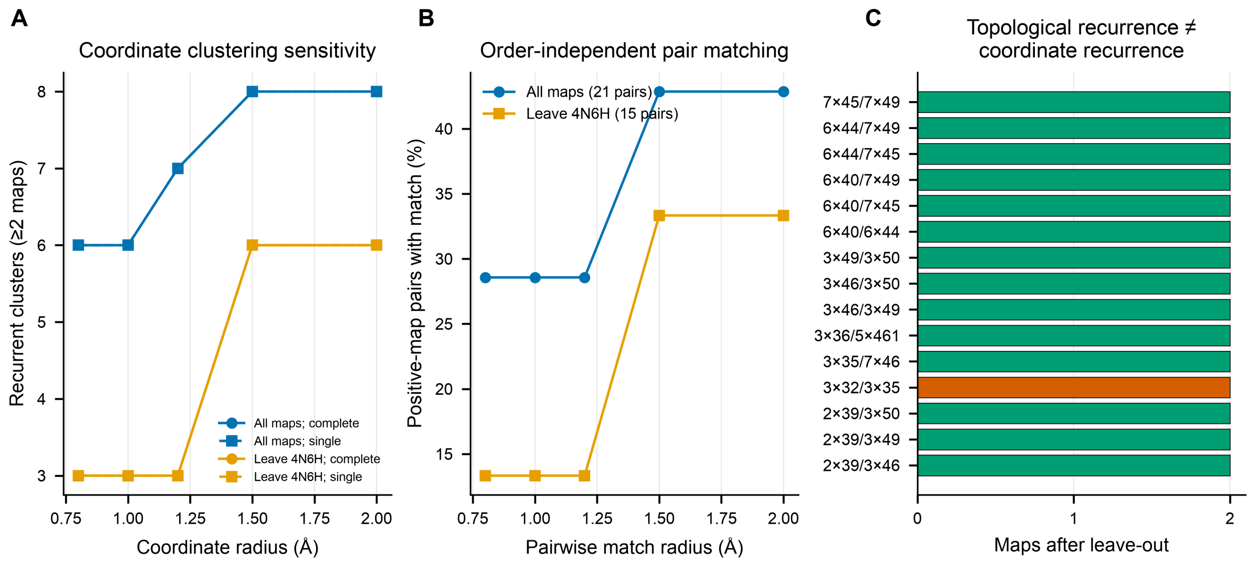


**Figure S6. Coordinate and topological water-recurrence sensitivity.** For default exact-BW bridging waters, a complete-linkage radius of 1.0 Å gives 6 recurrent coordinate clusters and 3 after excluding 4N6H. Pairwise matching finds overlap in 6/21 positive-map pairs and 2/15 after exclusion. Topological recurrence is distinct: 24 BW sites and 25 BW pairs recur in at least two maps, decreasing to 15 and 15 after exclusion. The 3×32/3×35 pair is highlighted. These summaries do not establish a general hydration effect.

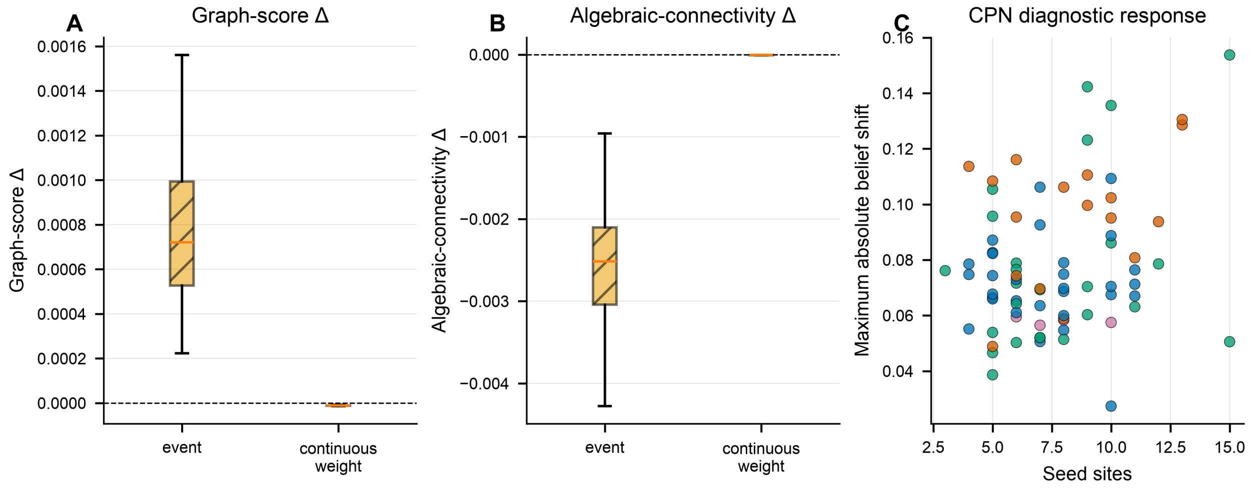


**Figure S7. StrucFlow and CPN computational diagnostics.** (A,B) Graph-score and algebraic-connectivity changes for event and continuous-weight modes across 84 independent maps per mode. (C) Maximum CPN belief shift versus seed count for 77 source-qualified primary-map seeded runs. Events are not physical time; weights and samples are not molecular dynamics; scores are not free energies; no ligand, state, component, subtype, efficacy, signaling, mechanistic, or causal effect is established.

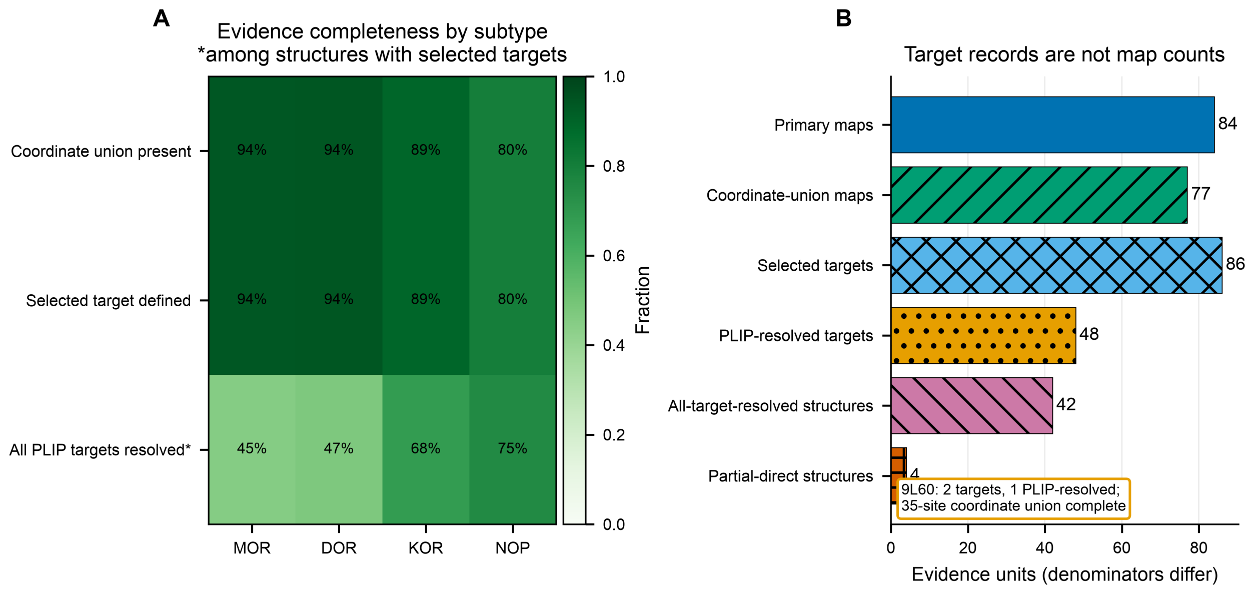


**Figure S8. Target- and coordinate-level evidence completeness.** Completeness is calculated from the curated coordinate union and selected-ligand target annotations. Coordinate role-aware sites occur in 77/84 maps. The target annotation set contains 86 targets across 77 structures; PLIP resolves 48 targets, 42 structures have all selected targets resolved, and 4 have a partial direct set. In 9L60, dynorphin resolves by PLIP while MPAM does not, yet the coordinate union is complete at 35 sites. These values describe evidence layers, not biological absence.

**Machine-Readable Methods and Data Index**

The accompanying public supplement contains the curated cohort, exact-BW mappings, single-structure graph summaries, exhaustive pair summaries, component and water estimands, spatial classifications, state and component statistical outputs, and non-image source data underlying every figure. Pseudocode and workflow specifications are limited to methods introduced in this study. The following stable artifact groups define the reproducibility record.

| Artifact group | Content | Reproducibility role |
|---|---|---|
| MR01 | Human-readable ordered pseudocode for study-introduced methods | Scientific operations and claim boundaries |
| MR02 | Machine-readable JSON specification for study-introduced methods | Ordered operations, inputs, parameters, equations, outputs, and quality checks |
| MR03 | Method parameters and analysis-contrast matrix | Every threshold, scope, reference level, bootstrap count, seed rule, and multiplicity family |
| MR04 | Equation and symbol definitions | Formulas, units, denominators, and missing-value rules |
| MR05 | Water estimand definitions and instance membership | Boolean membership for $P$, $C$, $R_2$, $B_{\text{any}}$, $C_{\text{B1}}$, $C_{\text{B2}}$, $L$, and branch overlap |
| MR06 | Authoritative table schemas and data dictionary | Field types, keys, units, allowed values, null policy, and analysis role |
| MR07 | Checksum-linked derivation and claim-to-source records | Input/output artifact IDs, record keys, fields, rounding rules, and validation status |
| MR08 | Non-image figure source data and package checksums | Panel-level numerical and categorical sources, without rendering code, plus integrity verification |

Each reported Article or SI value is indexed by a stable claim identifier, artifact identifier, record key, and field name. Scientific inputs and outputs are connected by SHA-256 checksums. Human-readable rounding does not replace the full machine-readable value.
